# Gene contribution of *Streptococcus dysgalactiae* subspecies *equisimilis*, an emerging pathogen, to experimental primate necrotizing myositis

**DOI:** 10.64898/2026.02.10.705160

**Authors:** S. M. Nayeemul Bari, Jesus M. Eraso, Randall J. Olsen, Luchang Zhu, James M. Musser

## Abstract

*Streptococcus dysgalactiae* subspecies *equisimilis* (SDSE) is an emerging human pathogen closely related to group A streptococcus. However, its genetic requirements for survival and growth in different conditions and for causing invasive infections remain poorly understood. To address this gap, we used <u>T</u>ransposon-<u>D</u>irected Insertion-site <u>S</u>equencing (TraDIS) to identify genes contributing to fitness in experimental necrotizing myositis in non-human primates (NHPs). Using two SDSE *stG62647* human clinical isolates, MGCS36044 and MGCS36089, we generated highly saturated transposon mutant libraries and analyzed them following *in vitro* growth and *in vivo* infection in eight NHPs. We identified 398 essential genes shared by both strains during growth *in vitro* and *in vivo*, and 17 and seven conditionally essential genes required only *in vitro* or only *in vivo*, respectively. Additionally, we identified 117 and 110 genes in MGCS36044 and MGCS36089, respectively, that were associated with fitness during necrotizing myositis. Transposon insertions in 34 MGCS36044 genes conferred increased fitness, whereas mutation of 83 genes conferred decreased fitness. Similarly, in MGCS36089, mutations in 38 and 72 genes conferred increased or decreased fitness, respectively. Importantly, both strains shared 46 fitness-associated genes, including an enrichment of transporter genes, highlighting nutrient acquisition as a dominant requirement during infection. The results provide critical information for guiding future translational efforts to develop preventive and therapeutic strategies against human SDSE infections.

## INTRODUCTION

*Streptococcus dysgalactiae* subspecies *equisimilis* (SDSE) is a Gram-positive beta-hemolytic bacterial pathogen that is closely related to group A streptococcus (GAS) ^1^. SDSE isolates commonly express Lancefield group C or G carbohydrate antigens ^2^. SDSE infects humans and several animal species ^3^. In humans, SDSE causes a spectrum of diseases similar to those caused by GAS. However, compared to GAS, the molecular pathogenesis of SDSE infections are less well understood ^4^. SDSE and GAS share some putative or proven virulence factors ^5–8^, including streptokinase and C5a peptidase and potentially FasBCA ^9^. The severity of SDSE infections can mirror that of GAS infections ^10, 11^. For example, SDSE and GAS disease manifestations range from mild pharyngitis, tonsillitis, and skin infections such as erysipelas, to severe invasive infections, such as cellulitis, bacteremia, osteoarticular infections (OAI), necrotizing fasciitis, and toxic shock syndrome ^12, 13^. The disease burden due to SDSE has not been quantified extensively but increasing incidence rates of invasive SDSE infections have been reported recently in several countries ^9, 14–16^. As a consequence, many investigators consider SDSE an emerging infectious agent.

Similar to GAS, SDSE isolates have been classified based on sequence variation in the M protein, a surface-displayed molecule that is highly variable. SDSE strains assigned to *emm* type *stG62647* are known to cause severe clinical manifestations, including necrotizing soft-tissue infections. Human infections caused by *emm stG62647* isolates have been increasingly reported globally, but the underlying cause is unknown ^15, 17–24^. More recently, genomic sequencing analysis, mouse virulence analyses, *in vitro* and *in vivo* transcriptome analyses using non-human primates (NHPs) as infection models, and analyses of large, comprehensive collections of SDSE clinical isolates ^17, 25–27^ have contributed much-needed new insight into SDSE pathobiology.

We used genome-wide transposon mutagenesis ^28–36^ to identify SDSE genes contributing to fitness in experimental necrotizing myositis in NHPs. Specifically, we used TraDIS (<u>Tra</u>nsposon-<u>D</u>irected Insertion-site <u>S</u>equencing) to identify genes contributing to survival and pathogenesis in two closely related *stG62647* SDSE human isolates, MGCS36044 and MGCS36089 ^17, 25, 26^. Our data suggest that SDSE transporters play a pivotal role in severe invasive infection. We also identified essential genes shared by both strains during growth *in vitro* in rich media. Collectively, these results provide information critical to guiding future translational efforts to better define the molecular basis of SDSE-human interactions and develop preventive and therapeutic strategies.

## MATERIALS AND METHODS

### Bacterial isolates and growth conditions

The two SDSE isolates MGCS36044 and MGCS36089 have been described previously ^17^. They are genetically closely related, differing from one another by 571 single nucleotide polymorphisms (SNPs) in the core genome. With one important exception (*fasB* in the *fas* operon), they have identical alleles of all genes known to encode major regulators such as two-component systems and stand-alone regulatory genes ^26^. SDSE strains were grown *in vitro* in Todd-Hewitt broth (Becton, Dickinson & Co.) supplemented with 0.2% yeast extract (THY medium) as previously described ^37^. Trypticase soy agar supplemented with 5% sheep red blood cells (Becton, Dickinson & Co.) was used as solid media for colony-forming units (CFU) determination. When required, THY was supplemented with erythromycin at a final concentration of 0.5 µg/mL. *Escherichia coli* strains were grown in Luria-Bertani (LB) medium at 37°C unless otherwise indicated. The LB medium was supplemented with 150 µg/mL erythromycin as needed.

### Generation of SDSE transposon mutant libraries

The plasmid pGh9:*ISS1* ^38^ was extracted from *E. coli* DH5α with a QIAprep miniprep kit (Qiagen, MD), suspended in 100 µL water, and electroporated into SDSE isolates MGCS36044 and MGCS36089 as described ^35^. After transformation, cells were grown in Medium II ^35^ for 1.5 h at 30°C to allow for phenotypic outgrowth, i.e., expression of the gene conferring erythromycin resistance. Each strain was plated on THY-E and incubated at 30°C for three days. Subsequently, the 20 plates corresponding to each strain were scraped into a tube containing THY/glycerol, resulting in a very dense bacterial solution (∼10^10^ CFUs) that was aliquoted and stored at –80°C for subsequent use.

We inoculated 100 µL aliquot of these cells into each of 12 tubes containing 40 mL pre-warmed THY-E. Plasmid pGh9:ISS*1* is temperature-sensitive (*ts*) for replication, where 30°C is the permissive temperature and 37° is non-permissive ^38^. In addition to 37°C, we also used 40°C in this work as the non-permissive temperature because this plasmid had not been used before with SDSE. Cells were grown for 3 h at both non-permissive temperatures. After growth, cells in each tube were centrifuged at room temperature for 10 min at 4000 rpm, washed in phosphate buffer solution (PBS), suspended in 2 mL THY/glycerol, and all cells per strain were mixed, aliquoted, and stored at –80°C.

### Preparation of the input transposon mutant libraries

We followed a protocol (**Fig. S1)** previously used for experimental GAS infections of NHPs and modified for use with SDSE. We created a 350 µL-mixture by combining 175 µL from the 37°C and 175 µL from the 40°C heat-shocked transposon mutant libraries for each of the MGCS36044 and MGCS36089 isolates. These mixtures were inoculated into four 50 mL Falcon tubes containing 30 mL THY-E. Cells were grown for 2 h at 37°C to an ∼OD_600_ = 1.5. We re-inoculated 4 mL from each tube into two (A and B) fresh 50 mL Falcon tubes containing 30 mL THY-E and grown for an additional 6 h. This step resulted in library expansion due to selection on erythromycin and elimination of unwanted dead cells. After growth for 6 h, the OD_600_ values ranged between 1.2–1.4, corresponding to 1.8–2.1 x 10^9^ cells/mL, based on the correspondence of 1.5 x 10^9^ SDSE when OD_600_ = 1. The A and B tubes corresponding to each of the 4 replicates for each strain (n = 16) were centrifuged for 10 min at 4°C, and the pellets were suspended and washed with 20 mL ice-cold PBS. The tubes were centrifuged again, and the cell pellets were suspended in 5 mL cold PBS/glycerol. We used 1 mL to determine the titer on THY-E and trypticase soy agar supplemented with 5% sheep blood plates (Becton Dickinson), and the remaining cells were stored in 16 cryovials (1 mL each from A and B tubes for four replicates of the two strains) at –80°C. Before the expansion of the input libraries, ∼1000 times more CFUs grew on blood agar plates, compared to the selective THY-E plates, suggesting that a some of the bacteria did not contain transposon insertions. By comparison, the number of CFUs was 6.2 x 10^7^–1.1 x 10^8^ on both selective and non-selective plates after the expansion, indicating that virtually all bacteria lacking transposon insertions had been eliminated from the libraries (**Fig. S1**).

Prior to extraction of chromosomal DNA, one of each of the A and B frozen stocks from the four replicates from both strains (n = 16) was serially diluted to the 10^-4^ dilution, plated on THY-E, and grown O/N at 37°C. The cells were subsequently scraped from each plate into individual Fastprep tubes (MP Bio, CA) containing 900 µL cold water and stored at –80°C before chromosomal DNA extraction (**Fig. S1**).

### Preparation of output libraries for TraDIS analysis after infection of NHPs

We used a well-described NHP model of necrotizing myositis ^35, 37, 39–41^. Briefly, eight cynomolgus macaques (3 to 5-year-old, 2–4 kg males and females) were used for the MGCS36044 and MGCS36089 transposon mutant library screens. NHPs were sedated, inoculated in the right quadriceps with the MGCS36044 input library and left quadriceps with the MGCS36089 input library, observed for 24 h, and necropsied. Each inoculum had 1 × 10^9^ CFU (**Fig. S1**). To analyze the output libraries, an approximately 0.5 g biopsy (0.5–1.0-cm diameter) of necrotic muscle was obtained from the center of the myositis lesion (eight NHPs x two strains; n = 16 replicates), homogenized in 1 mL PBS, transferred to 40 mL selective THY-E, and incubated for 6 h at 37°C for library expansion. Before incubation, 100 μL of homogenate was removed from the 40 mL-culture, serially diluted in PBS, and plated for CFU quantification. After growth, the output libraries were pelleted by centrifugation, washed with PBS, suspended in 2 mL PBS with 20% glycerol, aliquoted into cryogenic tubes, and stored at –80°C.

Frozen stocks corresponding to the eight replicates from each strain (n = 16) were serially diluted to the 10^-4^ dilution, plated on THY-E plates, and grown O/N at 37°C. The cells were subsequently scraped from each plate into individual FastPrep tubes (MP Bio, CA) containing 900 µL cold water and stored at –80°C before chromosomal DNA extraction (**Fig. S1**).

### DNA preparation and transposon mutant library sequencing

Preparation of genomic DNA and sequencing of transposon mutant libraries were performed as previously described ^35^ with slight modifications. Genomic DNA of each collected mutant pool was isolated with the GenElute bacterial genomic DNA kit (SIGMA_Aldrich) according to manufacturer’s protocol. DNA concentrations corresponding to either 10^-1^ to 10^-3^ (input libraries) or 10^-1^ to 10^-4^ dilutions (output libraries) were measured with a Nanodrop 8000 (Thermo Fisher Scientific, Waltham, MA). The dilutions were pooled, provided that the DNA concentration was ≥ 100 ng/mL. The final concentration of the pooled chromosomal DNA samples was measured again.

To prevent either over- or under-fragmentation of the chromosomal DNA, a pilot experiment was performed with 3 µg of chromosomal DNA treated with DNA fragmentase (NEB, MA) at 37°C in a time course experiment. After visual inspection of the DNA samples in an agarose gel, the optimal fragmentase reaction time for 3 µg of chromosomal DNA was determined to be 35 min. Subsequent library generation steps were performed using the NEBNext Ultra II DNA library prep kit for Illumina (NEB, MA) following manufacturer’s instructions. These steps were: (i) end Prep reaction to repair DNA fragment ends, (ii) ligation of a custom adapter, and (iii) polymerase chain reaction (PCR) amplification. Bam*HI* (NEB, MA) digestion was performed to cut the pGh9:*ISS1* plasmid backbone before PCR amplification to amplify the insertion site transposon-chromosome junction. The PCR-amplified libraries were sequenced with a NextSeq500/550 instrument (Illumina) using a single-end 75-cycle protocol and custom primers. All primers used in this study are listed in **Table S1**.

The average size for DNA fragments in each library was evaluated with DNA ScreenTape on a TapeStation 2200 (Agilent Technologies). The concentration of each sample was measured fluorometrically with Qubit dsDNA Broad Range and High Sensitivity kits (Thermo Fisher Scientific, Waltham, MA).

### Processing of DNA sequencing reads and data analysis

FastQ files were demultiplexed, and adapter contamination and read-quality filtering were performed with Trimmomatic v. 0.40 ^42^. Sequence data quality was determined with FastQC ^43^.

The mean per-base sequence quality was ≥ Q30 for all libraries. The sequencing reads were mapped to the respective cognate closed reference isolate genome ^17^ with SMALT (https://ftp.sanger.ac.uk/pub/resources/software/smalt/) as part of the BioTraDIS v. 1.4.5 pipeline ^44^, as previously described ^35^. The eight TraDIS input libraries (two isolates X four replicates) produced between 18–34 M reads, which resulted in 15–29 M after removing adapters using Trimmomatic v. 0.4.

Bacteria_tradis ^44^ was used to trim transposon tag sequences and map the reads to the reference genome of isolates MGCS36044 and MGCS36089. The plot files generated by bacteria_tradis were analyzed by tradis_gene_insert_sites to create spreadsheets listing the read count, insertion count, and insertion index for each gene. The output files from the tradis_gene_insert_sites analysis were transferred to tradis_comparison.R to compare the compositions of the input mutant libraries grown on THY-E at 0.5 µg/mL to output libraries after infection of the NHPs. Gene mutations with significantly altered frequency (log_2_ fold change of > 1 or < –1, q value of < 0.05) in the recovered mutant pools were considered as potentially altering SDSE fitness. To determine which genes were essential, conditionally essential, or involved in fitness, the input and output libraries from both isolates were compared (**Fig. S2)**. The functional analyses of fitness and essential genes were based on the clusters of orthologous groups (COGs) ^45^ using FACop.v2 (http://facop.molgenrug.nl/), NCBI, and manually assigning COG categories based on the genome annotation provided by Prokka ^46^.

## RESULTS

### SDSE isolate selection and generation of a highly saturated transposon mutant library

We hypothesized that, similar to GAS ^35^, SDSE adapts to conditions found in necrotizing myositis in NHPs. We also hypothesized that, compared to *in vitro* growth in rich media, certain SDSE genes contribute either positively or negatively to this adaptation, resulting in measurable fitness changes. To test these hypotheses, we used TraDIS, which has been used successfully to analyze several pathogenic streptococcal species ^35, 38^, but not SDSE.

We generated transposon mutant libraries in two well-characterized invasive human isolates, MGCS36044 and MGCS36089, and analyzed them in quadruplicate (**Fig. S1**). These two isolates were selected because (i) they are genetically closely related representatives of an SDSE *emm* type (*stG62647*) that has been recently reported to cause severe human invasive infections ^15, 17, 26^; (ii) experimental virulence and transcriptome data are available for them ^17, 25, 26^; and (iii) although genetically related, these isolates mainly differ in that MGCS36044 has wild-type alleles for all major transcriptional regulators, whereas MGCS36089 has a 40-nt deletion early in *fasB*, a gene encoding one of the two histidine kinases in the *fasBCAX* operon. This deletion has been associated with several transcript changes ^26^.

High-density transposon mutant libraries were generated for both isolates with integrative plasmid pGh9:IS*S1* ^35, 38^. The MGCS36044 library had an average of 191,293 unique insertions, representing an average spacing of one insertion every 11.45 nucleotides (**Fig. 1A** and **Fig. S3A**). Similarly, the MGCS36089 library had 196,089 unique insertions, corresponding to an average spacing of one insertion every 11.25 nucleotides (**Fig. 1B** and **Fig. S3B**). Collectively, these libraries represented extensive genomic coverage, with at least one insertion identified in 83% (n = 1778 of 2138 genes) of MGCS36044 genes and 81% (n = 1737 of 2145 genes) of MGCS36089 genes (**Fig. S4**). This level of saturation provided an average of 89 and 91 insertions per open reading frame (ORF) for MGCS36044 and MGCS36089, respectively (**Fig. S3**).

**Figure 1.**
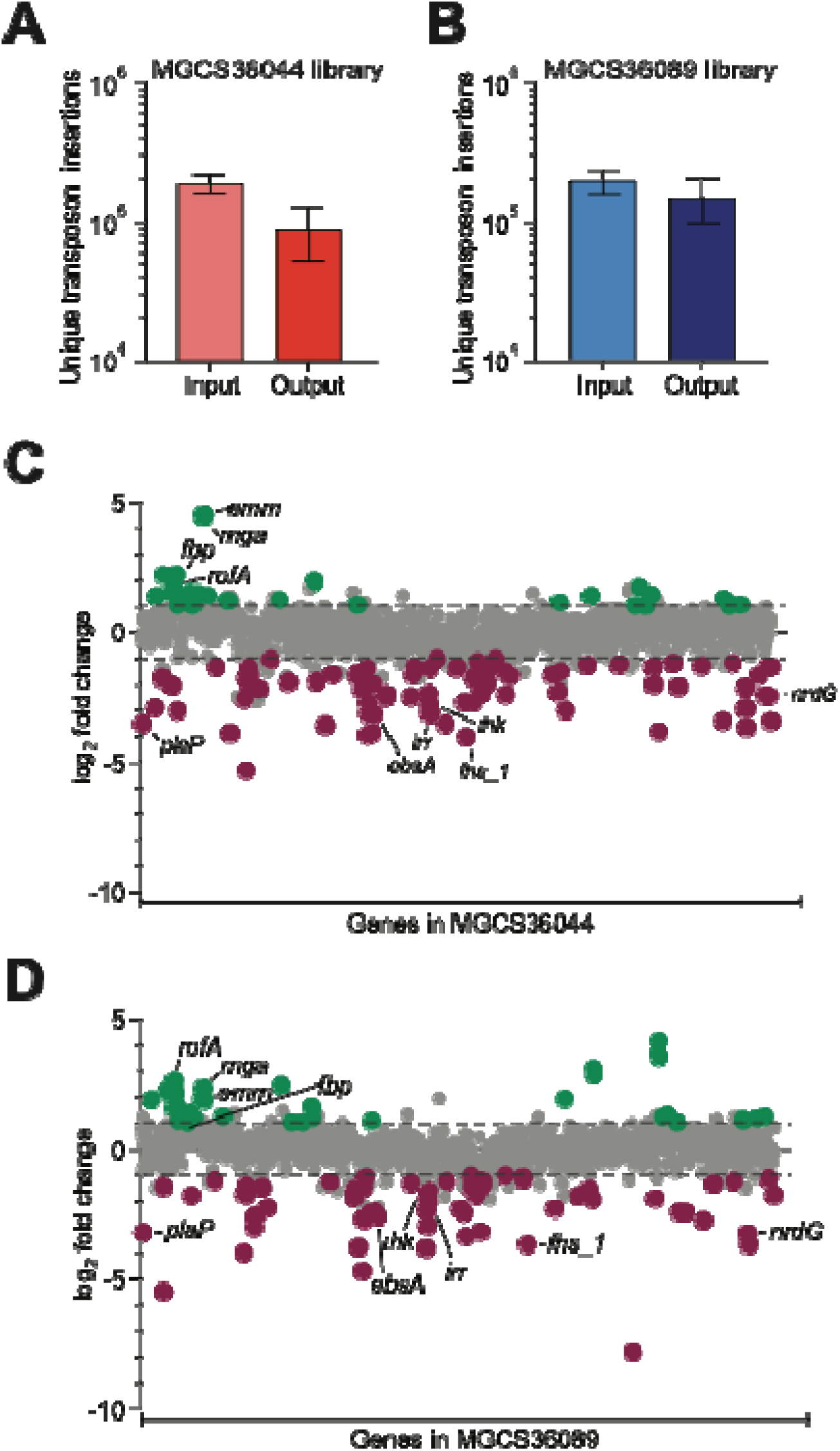
TraDIS analysis of SDSE gene fitness during necrotizing myositis in experimental NHP infection. Comparison of the complexity of (**A**) MGCS36044 and (**B**) MGCS36089 mutant libraries before (n = 4 replicates) and after 24-hour experimental NHP infection (n = 6 NHPs), as assessed by the number of unique transposon insertions identified. Data shown as mean ± SD (Standard Deviation). Genome-wide view of fitness gene changes for (**C**) MGCS36044 and (**D**) MGCS36089. The y-axis denotes the log_2_ fold change (log_2_FC) in mutant abundance for each gene (x-axis) in output libraries relative to input libraries. Gene mutations (insertions) conferring significantly increased fitness (log_2_FC > 1, q < 0.05) are highlighted in green, while those resulting in significantly decreased fitness (log_2_FC < –1, q = 0.05) are highlighted in purple. Grey dots represent genes that didn’t meet the defined cut-off, as indicated by the horizontal dashed lines. Representative fitness genes shared by both isolates are labelled. NHP: non-human primate; SDSE: *Streptococcus dysgalactiae* subspecies *equisimilis*.

To assess *in vivo* fitness and gene essentiality we used an infection protocol used previously for GAS ^35^. Eight cynomolgus macaques were inoculated with 1.0 × 10^9^ cells from input transposon libraries. MGCS36044 and MGCS36089 libraries were inoculated into the right and left anterior leg muscle, respectively. All animals developed signs and symptoms consistent with necrotizing myositis and, as in previous studies, were necropsied at 24 h post-infection ^35, 37, 47^. Biopsies were collected from the purulent infection site and from an immediately adjacent tissue layer. Samples corresponding to the TraDIS output libraries, consisting of eight replicates, were designated as *in vivo* and compared to the input libraries used for NHP inoculation, which had been grown *in vitro* in rich media. Total bacterial cell counts (CFUs) were calculated from homogenized biopsy samples (**Fig. S5**). To ensure sufficient library representation, two samples for each strain (corresponding to two biological replicates) that yielded cell counts lower than the required threshold of 1.0 × 10^6^ cells were excluded from further studies (**Fig. S5**) ^35^.

### Identification of essential genes in SDSE

SDSE is understudied compared to GAS, and until now there were no studies addressing gene essentiality. Using MGCS36044 and MGCS36089, we set out to determine either separately or jointly: (i) the number and identity of essential genes after growth *in vitro*, *in vivo*, and in both conditions simultaneously, and whether these essential genes were encoded by the core or accessory genome (regions of difference [RODs]) ^17^; (ii) whether conditionally essential genes (essential either *in* vivo or *in vitro*, but not in both conditions simultaneously) could be identified; and (iii) the extent of essential gene overlap between MGCS36044 and MGCS36089, including strain-specific conditionally essential genes (**Fig. S6**).

For this purpose, we used the gene essentiality function in the BioTraDIS pipeline and compared the four replicates corresponding to the input (*in vitro*) library to the six replicates of the output (*in vivo*) library recovered from NHP skeletal muscle for both isolates (**Fig. S2**). We hypothesized that most essential genes would be shared by both *in vivo* and *in vitro* libraries. However, given the substantial differences between growth conditions, we expected a minority of genes to be conditionally essential, depending on the growth condition or the SDSE isolate.

Analysis of the MGCS36044 input library grown *in vitro* in rich media identified 531 genes (∼25% of the genome) essential for growth and survival. Similarly, 503 essential genes were identified for growth *in vivo* in primate skeletal muscle (**Fig. 2A**). Of these, 475 genes were shared by both *in vivo* and *in vitro* growth (**Table S2A**). In agreement with our hypothesis, most essential genes were shared under both growth conditions and belonged to the core genome. However, we also identified 56 (**Table S2B**) and 28 (**Table S2C**) conditionally essential genes that were essential *in vitro* or *in vivo*, respectively, but not simultaneously in both growth conditions (**Fig. 2A** and **Fig. S6**). Conditionally essential MGCS36044 genes *in vitro* included three genes involved in fatty acid (FA) synthesis: *fabH*, *fabG_2*, and *acpP_2* were conditionally essential *in vitro* but not *in vivo* (**Table S2B**), whereas 10 additional FA genes were essential both *in vivo* and *in vitro*. In addition, *polA*, encoding DNA polymerase I, was conditionally essential *in vitro*. *In vivo*, four genes involved in sulfur mobilization and iron-sulfur cluster biosynthesis, (*sufCDEB*) were among the 28 conditionally essential genes only *in vivo* (**Table S2C**).

**Figure 2.**
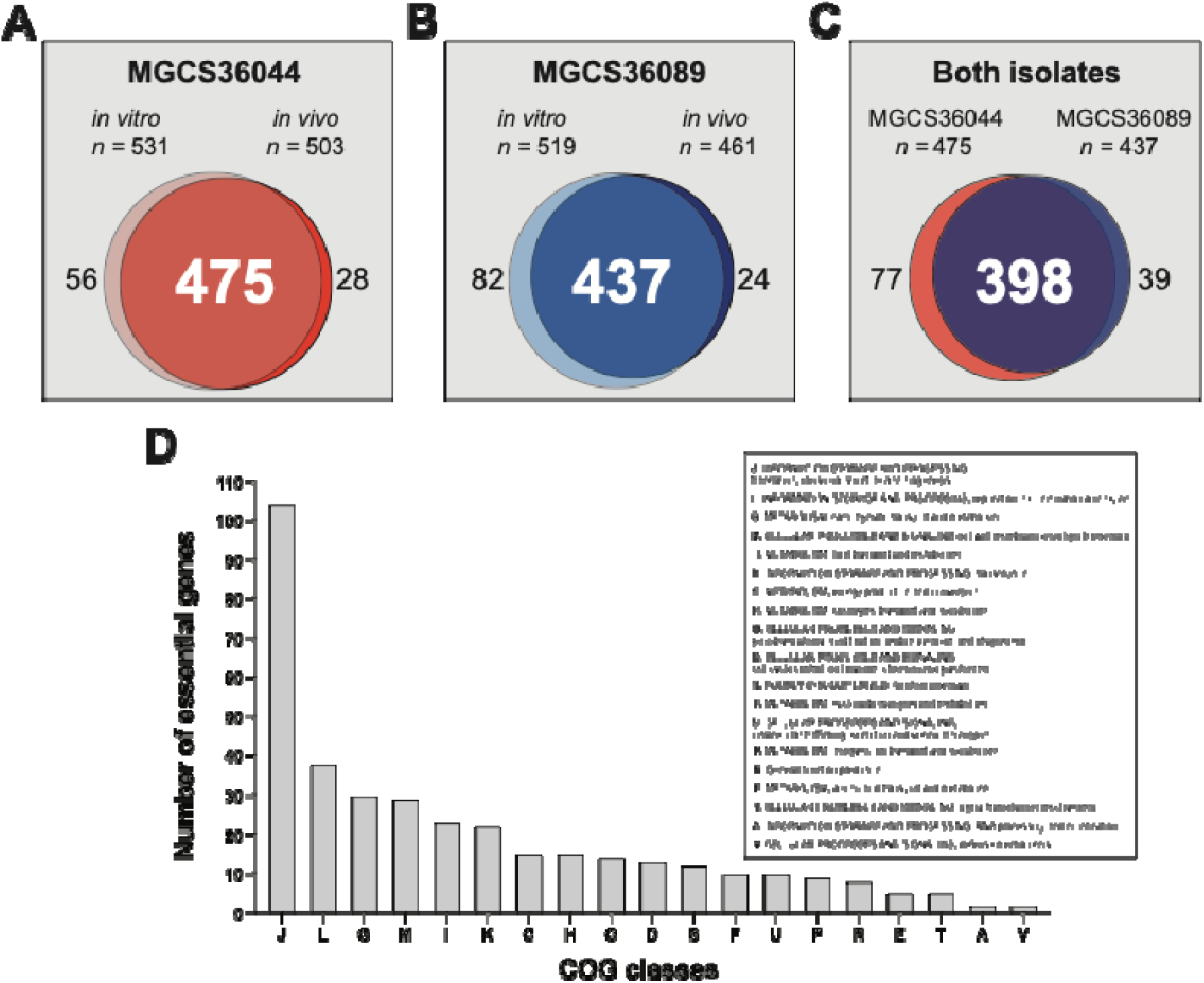
**Essential genes in SDSE isolates MGCS36044 and MGCS36089 *in vitro,* and *in vivo* during NHP necrotizing myositis**. Venn diagrams for MGCS36044 and MGCS36089 grown either *in vitro* (n = 4 replicates) or *in vivo* (n = 6 NHPs). For each condition, the genes reported were identified in all replicates. **(A)** For isolate MGCS36044, 475 genes were shared in both growth conditions, and 84 were conditional essential (56 genes identified only *in vitro*, and 28 genes identified only *in vivo*). **(B)** For isolate MGCS36089, 437 genes were shared in both growth conditions, and 106 were conditional essential (82 genes identified only *in vitro*, and 24 genes identified only *in vivo*). **(C)** 398 essential genes were shared by both isolates, and 116 were conditional essential (77 in MGCS36044, and 39 in MGCS36089). **(D)** Functional analysis, based on Cluster of Orthologous groups (COG) categories, for the 398 essential genes shared by MGCS36044 and MGCS36089. Transposase and small RNA genes were excluded. NHP: non-human primate; SDSE: *Streptococcus dysgalactiae* subspecies *equisimilis*.

Very similar results were obtained for MGCS36089. A total of 519 and 461 genes were essential *in vitro* or *in vivo*, respectively, with 437 shared genes (**Fig. 2B** and **Table S3A**). In addition, 82 (**Table S3B**) and 24 (**Table S3C**) genes were conditionally essential *in vitro* or *in vivo*, respectively. Thus, as hypothesized, most essential genes in both strains were shared *in vivo* and *in vitro*, with a smaller subset essential only under one growth condition.

When the data for these two isolates were compared, 475 and 437 genes were essential under both growth conditions in MGCS36044 and MGCS36089, respectively, and 398 of these were shared (**Fig. 2C** and **Table S4A**). Additionally, 77 genes were conditionally essential only in MGCS36044 (**Table S4B**), whereas 39 were conditionally essential only in MGCS36089 (**Table S4C**), also under both growth conditions (**Fig. 2C**). Several putative virulence genes were conditionally essential: *ccpA*, *vicR*, and *sagGHI* were essential in both strains and under both growth conditions. However, *covR*, *vicK*, and *liaR* were conditionally essential only in MGCS36044 (**Table S2B**), and *covS* was conditionally essential only in MGCS36089 (**Table S2C**). Several genes encoding ribosomal proteins were also conditionally essential. Among 49 ribosomal protein genes, 39 were essential and shared between both strains, whereas an additional seven genes were conditionally essential in MGCS36044 and 3 in MGCS36089. Importantly, 7 genes were conditionally essential in both strains *in vivo* but not *in vitro* (**Table S4D**), and 17 were conditionally essential *in vitro* but not *in vivo* (**Table S4E**). Thus, consistent with our hypothesis, conditionally essential genes were also identified when comparing these two closely related SDSE isolates, either under both growth conditions simultaneously, or individually, either *in vivo*, or *in vitro*.

Functional analysis of essential genes was based on COG categories ^45^, with the caveats that (i) some essential genes might have been originally assigned to more than one category and (ii) some genes have an unknown function ^48–50^. Among essential genes shared by both MGCS36044 and MGCS36089 under both growth conditions, the translation, ribosomal structure and biogenesis COG category (J) contained the largest number of essential genes (*n* = 104; **Fig. 2D**), including 47 genes encoding ribosomal proteins. This result is consistent with previous data obtained for GAS and other bacteria ^48–52^, and it underscores the overall importance of protein synthesis. An additional seven COG categories contained ≥ 15 genes/each (**Fig. 2D**).

Also, *ccpA* (encoding a global regulator that controls gene expression related to carbon metabolism), *vicR* (encoding a two-component system involved in cell wall and surface biogenesis and virulence), and *sagGHI* (encoding an ATP-binding cassette transporter responsible for the export of streptolysin S toxin) were essential in both strains and under both growth conditions.

### Essential genes in the core and accessory genome

Five genes in the RODs were essential in both strains under both *in vivo* and *in vitro* growth conditions (**Table S4A**). Located in ROD.3, MGCS36044_00610, MGCS36044_00622, and MGCS36044_00668 encode a putative XRE-type (<u>x</u>enobiotic response <u>e</u>lement) transcriptional regulator absent in GAS, a DNA cytosine methyltransferase, with roles in virulence and gene expression via epigenetic mechanisms ^53–55^, and a putative Cro/CI transcriptional regulator, respectively. MGCS36044_01838 and *xerS* are in ROD.4 and ROD.6, respectively, and encode a putative AbiEi-family antitoxin, part of a toxin-antitoxin (TA) system, located on an ICE in other bacteria ^56, 57^, and a site-specific tyrosine recombinase, which might have a role in genome maintenance and mobile genetic element integration ^58^.

### Identification of SDSE genes contributing to fitness in NHP necrotizing myositis

The TraDIS pipeline analysis was used to compare the input (in *vitro*) and output (in *vivo*) mutant libraries to identify genes with significantly altered frequencies *in vivo*. The MGCS36089 output library maintained high complexity, with a reduction of only ∼24% in unique insertion sites (**Fig. 1B**). Despite this reduction, genomic saturation remained robust, with at least one insertion in 83% (n = 1781 of 2145) of genes (**Fig. S4B**) compared to 81% (n = 1737 of 2145) in the input library. Notably, although the MGCS36044 output library had a greater than 50% decrease in unique insertion sites, the breadth of genomic coverage remained stable. That is, 85% (n = 1813 of 2138) of genes harbored at least one insertion in the output library compared to 82% (n = 1754 of 2138) in the input library. Even at a high threshold (e.g., five or more insertions per gene), the vast majority of the genome was represented in both the *in vitro* (input) and *in vivo* (output) libraries (**Fig. S4**). Together, these data suggest that although *in vivo* pressure resulted in a reduction in the number of unique insertion events, the library maintained sufficient diversity to provide confidence for the correct identification of fitness and essential genes with minimal error.

Fitness genes were identified as those in which transposon insertions caused a log_2_ fold change of ≥ 1.0 fitness increase, or log_2_ fold change of ≤ –1.0 fitness decrease, with q value ≤ 0.05. The contribution of a gene to fitness was determined relative to its wild-type allele; therefore, genes with an insertional inactivation that led to decreased necrotizing myositis pathogenesis were positive contributors to fitness, and *vice versa* (**Fig. S7**). According to these criteria, we identified 117 and 110 genes in MGCS36044 and MGCS36089, respectively that contribute to fitness in this infection model (**Fig. 3A-B**). For MGCS36044, insertional inactivation of 34 genes conferred increased fitness and 83 genes conferred decreased fitness (**Fig. 3A-B**). Similarly, for MGCS36089, 72 and 38 genes were identified as positive and negative contributors to fitness respectively (**Fig. 3A-B**). In total, 46 genes contributing to fitness were shared by both strains, including 13 genes identified as negative, and 33 genes identified as positive contributors (**Fig. 3A-B, Fig. S7,** and **Tables S5-S7)**.

**Figure 3.**
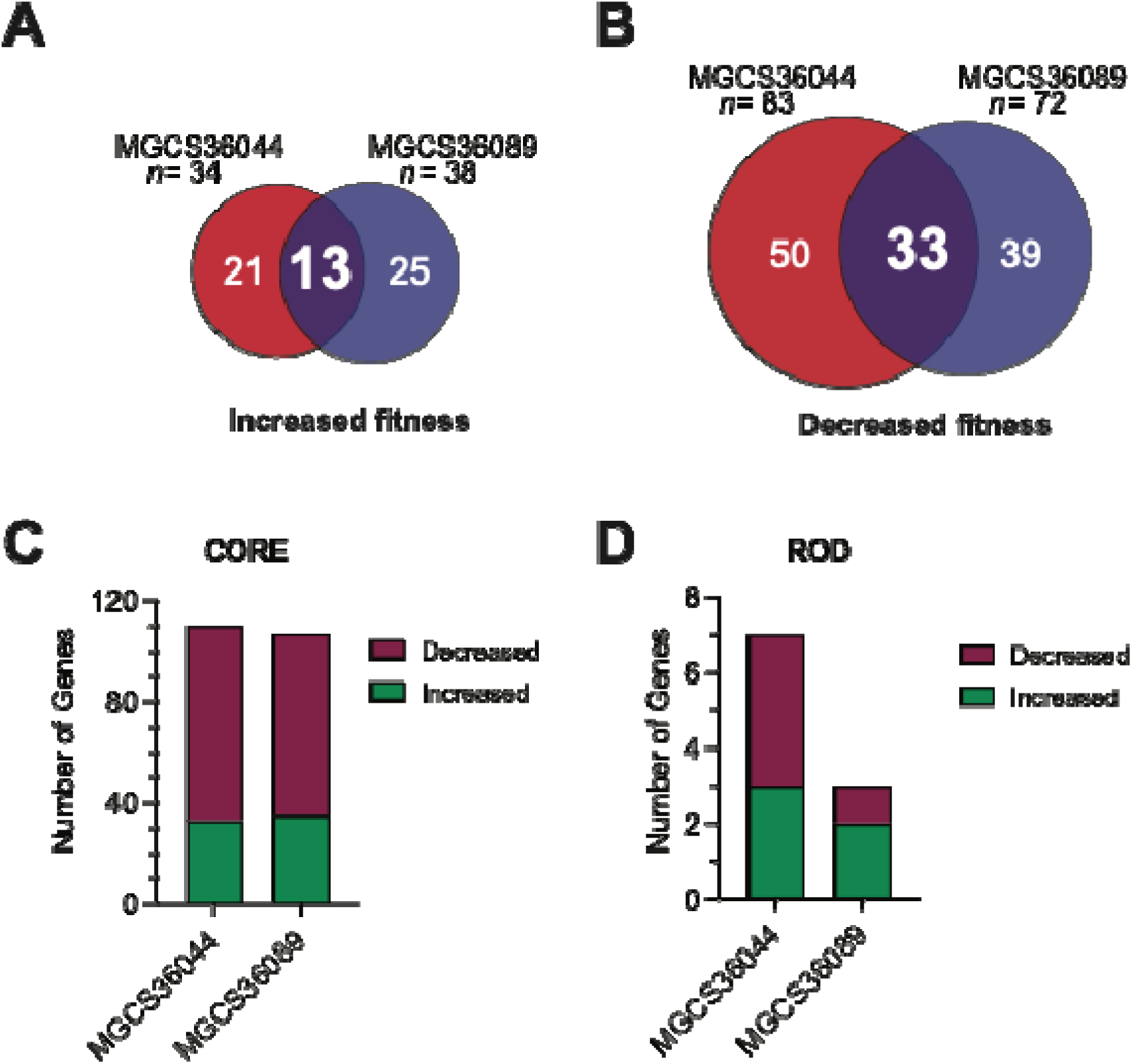
Comparative analysis of *in vivo* fitness gene determinants for SDSE isolates MGCS36044 and MGCS36089. Venn diagram showing the number of mutated genes conferring significantly (**A**) increased fitness and (**B**) decreased fitness in SDSE isolates MGCS36044 and MGCS36089 in NHP infections. (**C** and **D**) Distribution and number of fitness genes based on genomic location. The total number of genes conferring significantly altered fitness is categorized into those belonging to the core genome and region of difference (ROD). The bars are segmented to show the proportion of each set of genes where mutations resulted in either increased (green) or decreased (purple) fitness. NHP: non-human primate; SDSE: *Streptococcus dysgalactiae* subspecies *equisimilis*.

### Overview of genes influencing fitness in experimental necrotizing myositis

Although SDSE is closely related to GAS ^3, 17^, particularly with respect to the core genome, the accessory genome differs substantially between these two species. We identified 110 fitness genes in MGCS36044 that are part of the core genome, whereas seven genes were located in RODs (**Fig. 3C-D**). Transposon insertions in 33 core genes resulted in increased fitness, whereas mutational insertion in 77 core genes conferred decreased fitness. In the accessory genome, mutations in three and four ROD genes conferred increased and decreased fitness, respectively. Similarly, in MGCS36089, there were 107 core fitness genes (35 negative and 72 positive contributors, respectively) and three ROD genes, one with a negative and two with a positive contribution to fitness (**Fig. 3C-D**).

Functional analysis of MGCS36044 and MGCS36089 fitness genes based on COGs ^45^ revealed that the most prevalent COG functional categories included genes involved in (i) cell wall/membrane/envelope biogenesis (M), (ii) transcription (K), (iii) amino acid transport and metabolism (E), and (iv) ion transport and metabolism (P) (**Fig. 4**). These categories were also the most prevalent among the 46 fitness genes shared by both strains, highlighting a conserved functional backbone. With some notable exceptions, the overall COG distribution of SDSE fitness genes was similar to that reported for GAS ^35^, underscoring the genetic similarities between these two species ^3, 17^.

**Figure 4.**
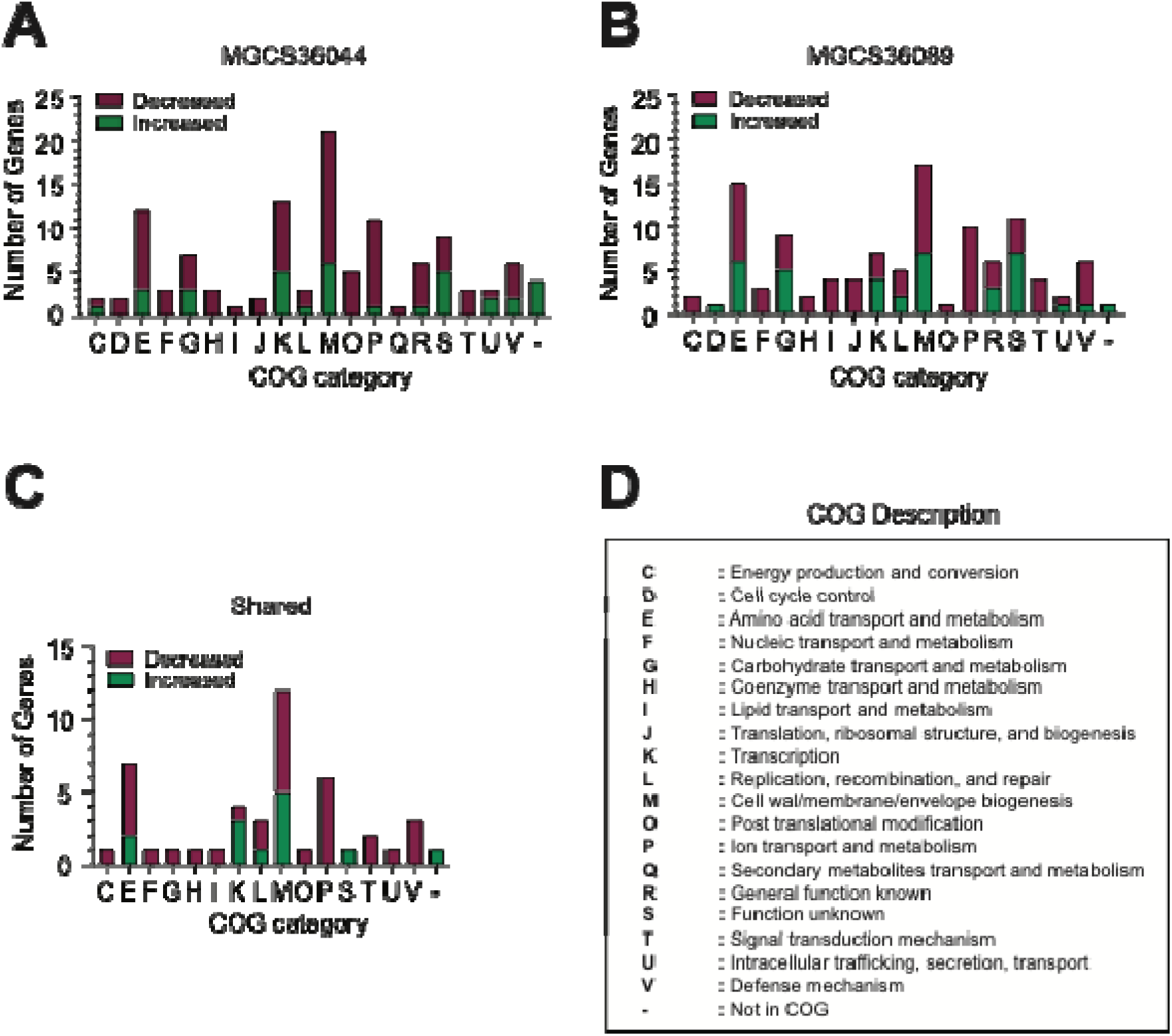
Functional categorization of the identified SDSE fitness-related genes in experimental NHP necrotizing myositis. Bar plots categorize the identified fitness genes (**A**) in MGCS36044, (**B**) in MGCS36089, and (**C**) shared by both isolates according to Cluster of Orthologous groups (COG). The bars are segmented to show the proportionate role of each set of genes for which mutations resulted in either increased (green) or decreased (purple) fitness. (**D**) Description of the functional categories. NHP: non-human primate; SDSE: *Streptococcus dysgalactiae* subspecies *equisimilis*.

In MGCS36044, genes contributing positively to fitness during NHP infection encoded proteins with functions related to either core metabolism (*galE*, *guaA*), carbon and nitrogen acquisition (*palP*), and stress management (*perR*, encoding a transcriptional regulator, and *sufB*, encoding a protein involved in iron-sulfur cluster synthesis). In addition, the *ihk*-*irr* two-component system, which is highly expressed *in vivo* in NHP skeletal muscle in both GAS and SDSE ^26, 37^, *tig* (encoding the trigger factor involved in general maintenance), and *fhs_1* (encoding formyl-tetrahydrofolate synthase), also contributed positively during NHP infection. Similar results were found for MGCS36089, where the major positive contributor to fitness included genes encoding proteins involved in core metabolism (*glgP*, *deoC, atoB* etc.), nutrient acquisition (*oppABCD, metP_1* etc.), and stress response (*sufD*). Notably, 33 fitness genes contributing positively were shared between the two isolates (**Fig. 3B** and **Table S7B**), including genes associated with carbon-nitrogen acquisition (*oppBC*), putative virulence (*sagE*), and transcriptional regulators, such as *ihk* and *irr*.

Unexpectedly, in both clinical isolates studied, several putative virulence genes showed the greatest fitness decrease contribution, including *mga*, a master virulence regulator in GAS ^59, 60^, *emm*, which encodes the M protein and in GAS is positively regulated by Mga ^61^; *rofA*, a pilus regulator ^62, 63^; and *fbp*, which encodes a fibronectin-binding protein and is part of the FCT-1 pilus region in SDSE (**Fig. 1C-D** and **Table S7A**).

Data for the glucose sensor and transporter operon (*manLMN*) pointed to a notable difference between the two clinical isolates studied here (**Table S6A**). This operon had a negative contribution to fitness in MGCS36089, but not in MGCS36044. Also, and in contrast, in GAS it had a positive contribution to fitness ^35^, highlighting strain- and species-specific differences in metabolic regulation during infection.

### Genes encoding putative transporters are major fitness determinants in SDSE in experimental NHP necrotizing myositis

Based on our data, the most significant genetic requirement for SDSE proliferation in primate skeletal muscle may be the efficient acquisition and assimilation of relatively limited host resources, as suggested by the largest functional class of fitness determinants identified represented genes encoding various putative transport functions. In MGCS36044, 29 of 117 genes (24.7%) encode putative transporters (**Fig. 5A**) and transposon mutagenesis in 25 of these genes resulted in decreased fitness (**Fig. 5B-C**), underscoring their essential role during infection. Similarly, in MGCS36089, 32 of 110 MGCS36089 fitness genes (29.0%) encode putative transporters. Of these, the majority (n = 23) contributed positively to fitness during infection (**Fig. 5A-B-C**). Collectively, enrichment of transporter genes among positive fitness determinants in both strains underscores the importance of acquiring necessary growth substrates during infection.

**Figure 5.**
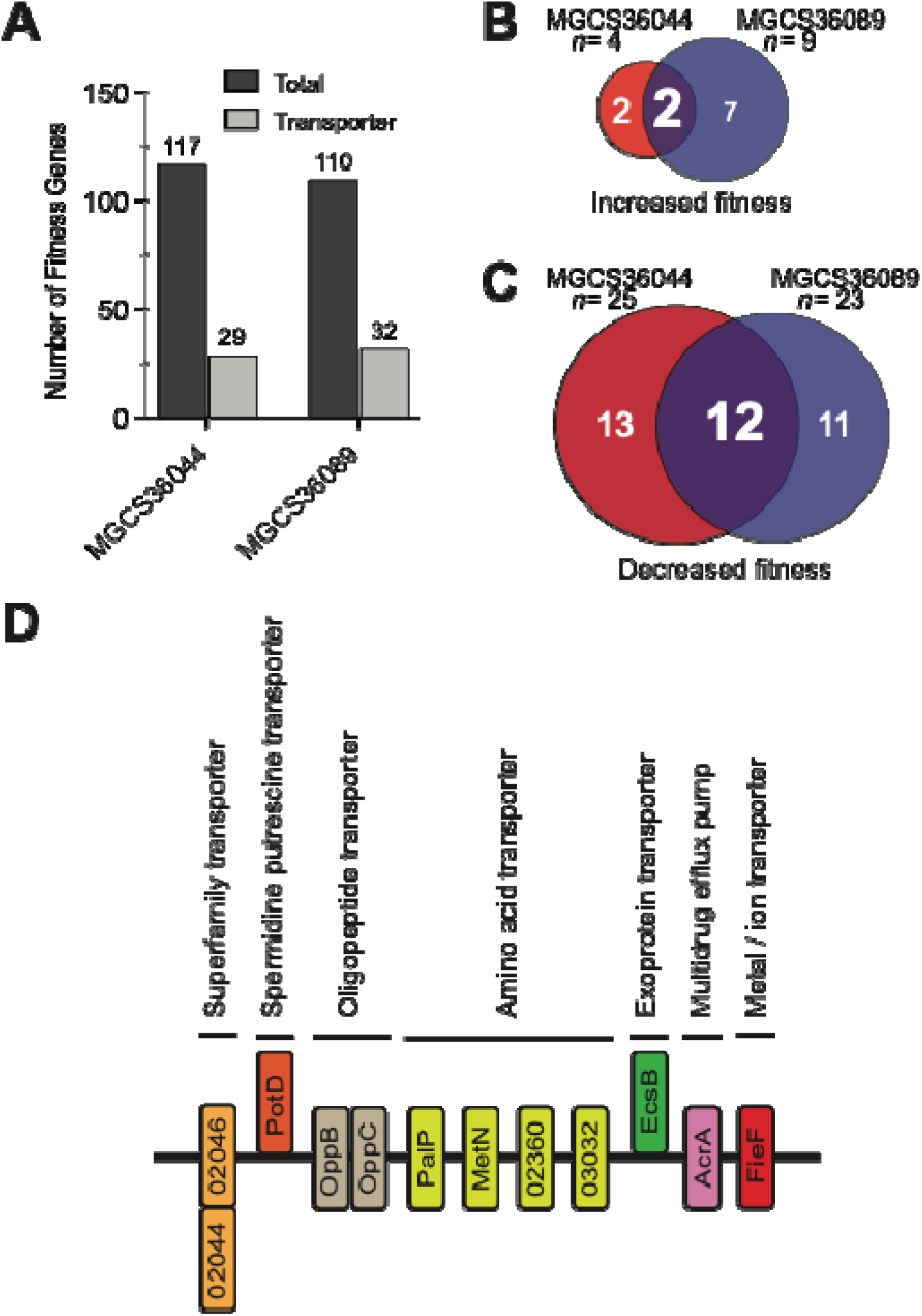
Genes encoding putative transporters are an abundant portion of fitness genes that are required in experimental NHP necrotizing myositis. **(A)** MGCS36044 fitness genes (n = 29, 24.7%) and MGCS36089 fitness genes (n = 32, 29.0%) that encode putative transporters. Venn diagram showing the relationship between MGCS36044 and MGCS36089 transporter genes in the context of mutation conferring (**B**) increased and (**C**) decreased fitness in NHP infection. (**D**) Schematic showing the putative transporter genes shared by both isolates that contribute positively during NHP infections. Putative substrate binding proteins are positioned outside the cell membrane. Elements that are positioned in the membrane and inside the bacterial cell are putative transmembrane and cytosolic proteins, respectively. Locus tag numbers refer to the annotation for isolate MGCS36044. NHP: non-human primate.

Importantly, the 12 transporter genes contributing positively to fitness were shared between MGCS36044 and MGCS36089 (**Fig. 5D**). These included genes likely involved in scavenging small peptides and exoproteins from the damaged host tissue, such as *oppB* and *oppC,* which encode proteins in an oligopeptide transport system, and *escB*, implicated in uptake of exoproteins. These findings indicate that SDSE relies heavily on peptide scavenging during NHP infection, perhaps to address potential amino acid scarcity. In addition to genes for peptide transport, conserved fitness determinants included *potD, acrA*, and *fief,* encoding a putative spermidine/putrescine transporter, a multidrug efflux pump, and a protein predicted to be involved in iron homeostasis, respectively. Together, these data suggest that acquisition of amino acids, polyamines, and metal ions represents an important fitness strategy for SDSE during necrotizing myositis in NHP.

## DISCUSSION

SDSE has increasing clinical importance due to the growing number of human infections it is causing. However, compared to its close genetic relative GAS, the molecular determinants contributing to SDSE survival, growth, and molecular pathogenesis are poorly understood. We recently showed that SDSE undergoes significant transcript remodeling during experimental NHP necrotizing myositis, marked by upregulation of metabolic and virulence gene expression ^26^. Although those transcriptome data identify genes active during infection, they do not identify essential genes or the specific contributions of individual genes to fitness in the host. Here, we addressed this knowledge gap using highly saturated transposon insertion sequencing (TraDIS) of two closely related SDSE isolates. These data defined (i) genes essential for growth *in vitro* and *in vivo* in NHP necrotizing myositis and (ii) genes that contribute positively or negatively to fitness in this experimental infection model. One strength of this study is the use of two genetically very similar *stG62647* clinical isolates that differ by only 571 core SNPs ^26^. By design, we used two isolates with slightly different genetic backgrounds due to mutation of *fasB*, which encodes one of the two histidine kinases in the *fasBCAX* operon. This design allowed us to infer fitness and biologically meaningful isolate-specific differences.

Although essential and conditionally essential genes establish the baseline requirements for viability, successful infection is shaped by genes that modulate fitness under varying host conditions. Across both isolates, we identified 117 and 110 fitness genes, respectively, with substantial overlap (46 shared genes). This overlap indicates that SDSE relies on a conserved set of functions to adapt from nutrient-rich conditions to the more restrictive environment of necrotic skeletal muscle. A striking finding in our study was the identification of genes whose inactivation conferred a significant fitness advantage (log_2_FC ≥ 1.0, q ≤ 0.05) in both isolates. This group includes well-characterized mediators of virulence in GAS, such as the transcriptional regulators *mg*a, and *rofA*, the M-protein encoding gene *emm*, and the fibronectin-binding protein encoding gene *fbp*, which is located in the FCT-1 pilus region (**Tables S7A**). In GAS, Mga is a global transcriptional regulator that positively regulates multiple virulence genes such as *emm*, *scpA*, *sof*, *fbpA*, and *sic* ^64–69^. Of note, these genes are often expected to contribute positively during NHP skeletal muscle infection, but they did not emerge as fitness determinants in our previous TraDIS screen in GAS ^35^, a finding underscoring fundamental differences between these two related species. Since in GAS both Mga and RofA positively regulate the expression of proteins involved in initial adhesion to the host, such as FbpA and pilus proteins ^61, 70–73^, their unexpected negative contribution to SDSE fitness suggests a possible adhesion-dispersion trade-off, where the loss of positive regulators for initial adhesion and/or adhesin gene expression may remove a dissemination barrier, as evidenced by the fact that binding to host plasma fibronectin can physically tie bacteria to the host matrix and limit spreading ^74^. Additionally, since adhesin genes are redundant, the loss of one might be compensated by others. Importantly, changes in *mga* and pili gene expression allow bacteria to adapt to environments, and to evade immunity ^75–80^. Of note, recent transcriptomic analysis of SDSE causing necrotizing fasciitis showed no significant differential upregulation of *mga*, *rofA*, *emm*, and *fbp* compared to *in vitro* growth ^26^. Altogether, our data suggest that SDSE fitness during necrotizing myositis might be governed by a strategic coordination of initial attachment to the host, dispersion, and immune evasion.

Our analysis identified 33 genes indispensable for positive fitness (**Fig. 3B**) in both strains during necrotizing myositis in the NHPs. The most salient feature of the SDSE fitness landscape was a substantial reliance on the nutrient acquisition systems, particularly transporters. Approximately one-third of the identified positive fitness determinants in both isolates were putative transporters. This finding mirrors previous data from GAS, where peptide/amino acid transport was crucial for deep tissue proliferation ^35^.

Although the overall fitness landscapes of MGCS36044 and MGCS36089 were very similar, notable isolate-specific differences were observed. In particular, the glucose sensor and transporter operon *manLMN* contributed negatively to fitness in MGCS36089 (**Table S6A**), contrasting with its reported role in GAS ^35^. This divergence highlights how subtle genetic differences can reshape metabolic priorities and regulatory networks, even among closely related strains, and alerts against assuming direct functional equivalence between GAS and SDSE.

The vast majority of fitness and essential genes were in the core genome, and only a few were encoded by RODs. Whereas in GAS the accessory genome is commonly associated with phage content, in SDSE it consists of RODs ^17^, with gene content resulting from integrative conjugative elements (ICEs) ^81^ and to a very limited extent phages ^82^. The fact that we found essential genes encoding putative transcriptional regulators, a DNA methyltransferase, an antitoxin in a TA system, and a site-specific recombinase, with putative roles in gene expression, virulence, resistance to phage infections ^83^, and chromosome dimer resolution ^58^, respectively, attests to an advantage potentially provided by the accessory genome in SDSE.

A novel finding was the identification of seven genes shared by MGCS36044 and MGCS36089 that were conditionally essential *in vivo* but not *in vitro* (**Table S4D)**. Although their exact functions are unknown in SDSE, and thus their putative function can only be inferred, they might have some future therapeutic potential. While we found *sufD* to contribute to fitness, *sufBC* (part of the *sufCDSEB* operon involved in iron-sulfur [Fe-S] cluster biosynthesis) was conditionally essential *in vivo*. Fe-S clusters are involved in multiple pathways, such as sensing stress due to oxidative and nitrosative damage, gene regulation, RNA modification, and DNA replication and repair ^84, 85^. Other conditionally essential genes *in vivo* were *whiA*, which encodes a transcriptional regulator acting as a switch incorporating environmental cues and internal signals and is involved in cell division ^86, 87^; *trxB,* which encodes a thioredoxin-fold lipoprotein involved in extracellular oxidative stress ^88^; *lgt,* which is involved in the attachment of prolipoproteins to the bacterial membrane (notably, its deletion leads to a marked decrease in virulence in *Streptococcus pneumoniae)* ^89^; and *murA_1,* which encodes one of the *murA* homologs with different functions ^90^, but catalyzing the first step in peptidoglycan synthesis. Thus, since these genes wer essential only *in vivo*, in the primate skeletal muscle, they constitute prime targets for future therapeutic intervention.

Finally, although we identified a partial overlap with GAS ^35^, suggesting conserved molecular mechanisms during necrotizing myositis, most fitness gene determinants were distinct, suggesting that the ability of SDSE to cause severe invasive disease is not merely a mimicry of GAS. As an example, *ccpA* was essential in SDSE but not in GAS ^91^, underscoring the importance of SDSE research to develop targeted therapies.

As clearly documented for SARS-CoV-2, for there emerging pathogens, it is important to rapidly acquire as much information as possible to enhance understanding of pathogen-human interactions. Studies based on genome-scale analyses such as those conducted here and in our prior work are ideal for identifying potential leads for subsequent investigation.

## ACKNOWLEDGMENTS

We declare no conflicts of interest.

These studies were funded in part by the Fondren Foundation (to JMM). We are grateful to Dr. Stephen B. Beres for his essential contribution to the setup and optimization of our TraDIS bioinformatics workflow. We thank Drs. Stara Robertson and Alphina Ho, and the Comparative Medicine Veterinary technologists, for veterinary technical assistance; Alma Amaya, Ryan Gadd, Regan Mangham, Eleanor Nichols, Matthew Ojeda Saavedra, Jordan Pachuca, and Sindy Pena for expert technical assistance; and Drs. Sasha Pejerrey and Heather McConnell for editorial assistance.

## Study approval

All animal experiments were approved by the Institutional Animal Care and Use Committee of Houston Methodist Research Institute (protocol IS00007782).

