## Supplemental Figures for "Gene contribution of *Streptococcus dysgalactiae* subspecies *equisimilis*, an emerging pathogen, to experimental primate necrotizing myositis"

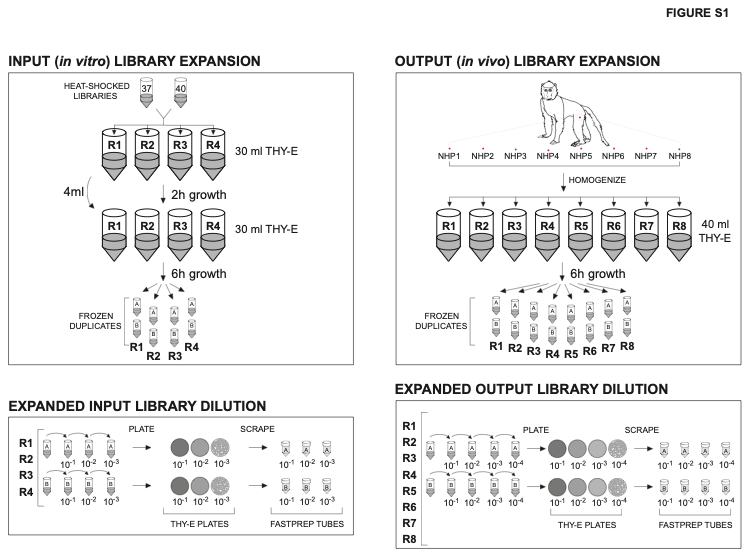

**Figure S1. Transposon library expansion and dilution**. **Left panel**. Input libraries (*in vitro*) for 4 replicates. **Top**. Equivalent amounts of dense (~10^10^ CFUs) 37°C and 40°C heat-shocked transposon mutant libraries of either isolate MGCS36044 or MGCS36089 were expanded in THY-E for ~8 h and frozen. **Bottom**. Expanded libraries for the 4 replicates were serially diluted, titered, and the 10^-1^–10^-3^ dilutions were plated on THY-E, individually scraped, and frozen. **Right panel**. Output libraries (*in vivo*) for 8 replicates. **Top**. 8 NHPs were each inoculated with MGCS36044 and MGCS36089 in the right and left legs, respectively. Biopsies corresponding to the site of infection were homogenized, expanded in THY-E, and frozen. **Bottom**. Expanded libraries for the 8 replicates were serially diluted, titered, and the 10^-1^–10^-4^ dilutions were plated on THY-E, individually scraped, and frozen. THY-E: THY containing 0.5 µg/mL erythromycin; R: replicate; NHP: non-human primate.

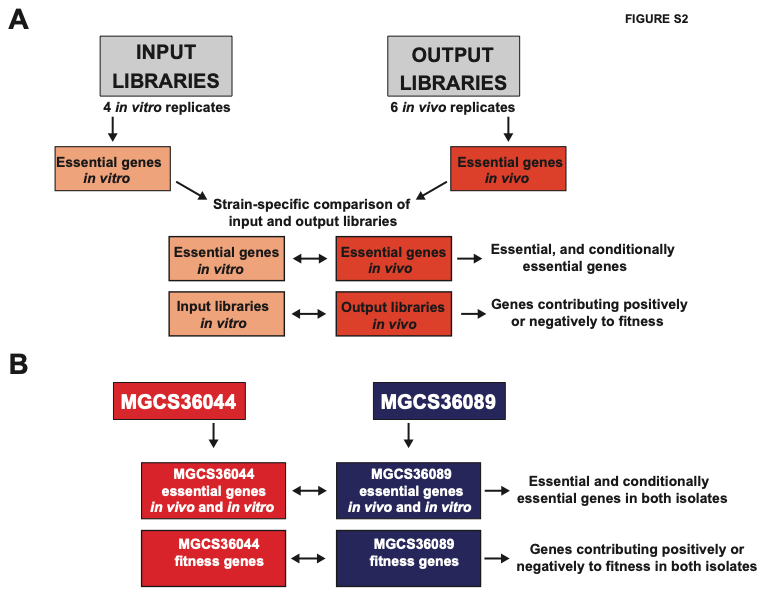

**Figure S2. Schematic for transposon library analysis**. **A**. Four input (*in vitro*) and 6 output (*in vivo*) libraries corresponding to MGCS36044 and MGCS36089 were analyzed with the BioTraDIS pipeline. Two of the original 8 *in vivo* output libraries were excluded (see Materials and Methods). For both isolates, essential genes were separately identified in the input and output libraries, and compared to conditionally essential genes, identified either *in vivo* or *in vitro*. Fitness genes were also identified comparing input and output libraries. **B.** Essential genes found both *in vivo* and *in vitro* in one isolate were compared to those in the second isolate to identify shared essential genes in both isolates. Fitness genes shared by both isolates were also identified by isolate comparison.

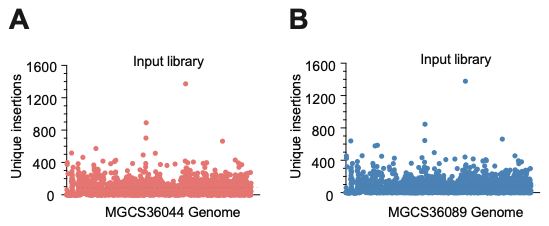

**Figure S3. Characterization of mutant input libraries.** Unique transposon insertion counts for each gene in the (**A**) MGCS36044 and (**B**) MGCS36089 input libraries. Dotted lines indicate the average number of unique insertions per open reading frames in the genome of each isolate (89 and 91 for MGCS36044 and MGCS36089, respectively).

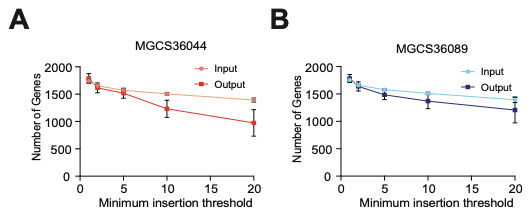

**Figure S4. Library saturation and depth across the SDSE genome.** The total number of genes with a minimum of 1, 2, 5, 10, and 20 unique transposon insertion sites was quantified to assess the depth of (**A**) MGCS36044 and (**B**) MGCS36089 mutant libraries. Data points represent the average of multiple biological replicates (Input n = 4; Output n = 6), with error bars showing the variation (SD) between replicates. SDSE: *Streptococcus dysgalactiae* subspecies *equisimilis,* SD: Standard Deviation.

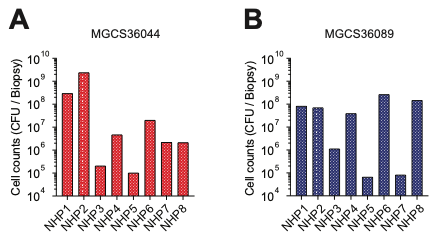

**Figure S5. Bacterial load and sample quality control of NHP tissue biopsies.** Quantification of total bacterial colony-forming units (CFUs) recovered from homogenized skeletal muscle biopsies 24 h post-infection for (**A**) MGCS36044 and (**B**) MGCS36089. Replicates (NHP3 and NHP5 for MGCS36044, and NHP5 and NHP7 for MGCS36089) yielding CFU below the established quality control threshold of 1X10^6^ CFU/biopsy were excluded from all subsequent downstream processing and analysis. The 10^6^ CFU limit ensures that the recovered transposon mutant population is sufficiently representative of the input library to provide statistically significant fitness data. NHP: non-human primate; SDSE: *Streptococcus dysgalactiae* subspecies *equisimilis.*

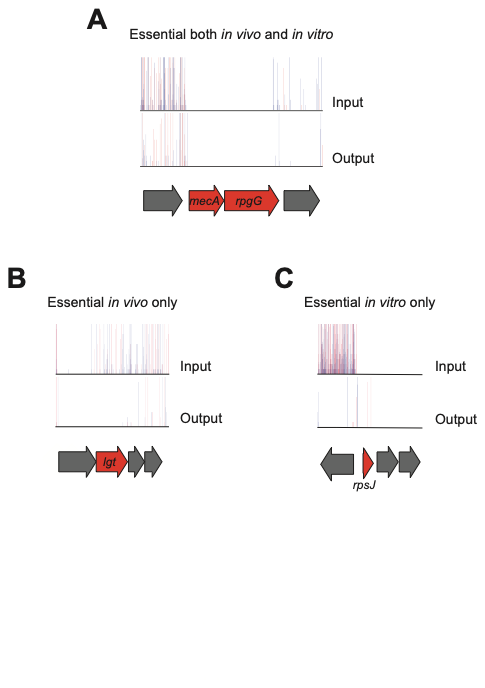

**Figure S6. Representative transposon insertion maps of conditionally essential genes in SDSE.** Insertion profiles are shown for genes that are essential (**A**) both *in vitro* and *in vivo*, (**B**) *in vivo* only, and (**C**) *in vitro* only, comparing the input library (*in vitro*), and the output library recovered after NHP infection (*in vivo*). Genes of interest are highlighted in red. All insertion maps are shown with respect to the SDSE strain MGCS36044. Each vertical bar represents a unique insertion site, and the height of the bar reflects the number of sequencing reads associated with that insertion. Blue and red bars indicate transposon insertions on the minus and plus DNA strands, respectively. NHP: non-human primate; SDSE: *Streptococcus dysgalactiae* subspecies *equisimilis.*

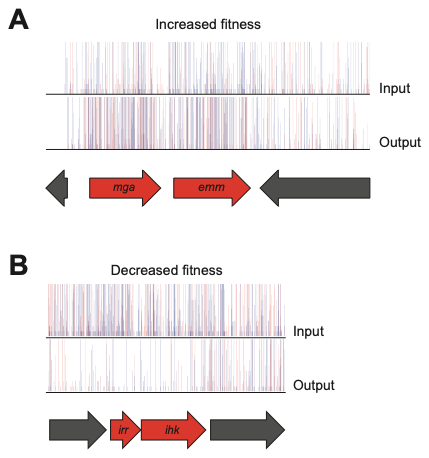

**Figure S7. Representative transposon insertion maps of fitness genes in SDSE mutant libraries before and after NHP infection.** Insertion profiles of genes for which a transposon insertion resulted in (**A**) increased or (**B**) decreased fitness during necrotizing myositis. For each gene, transposon insertion maps are shown for the input and output libraries. Genes of interest are highlighted in red. All insertion maps are shown with respect to the SDSE isolate MGCS36044. Each vertical bar represents a unique insertion site, and the height of the bar reflects the number of sequencing reads associated with that insertion. Blue and red bars indicate transposon insertions on the minus and plus DNA strands, respectively. The results shown here are consistent with the fold changes listed in Table S5. NHP: non-human primate; SDSE: *Streptococcus dysgalactiae* subspecies *equisimilis.*
