## Supplementary material for "Gene contribution of *Streptococcus dysgalactiae* subspecies *equisimilis*, an emerging pathogen, to experimental primate necrotizing myositis": Table S7B

**Table S7B. List of genes shared between MGCS36044 and MGCS36089 where transposon insertion conferred decreased fitness.**

| **No. ^†^** | **Locus tag 044^∫^** | **Locus tag 089^∫∫^** | **Gene** | **Function** | **logFC 044^∫^** | **logFC 089^∫∫^** | **Transport** | **Core/ Acc^††^** | **COG class** |
| --- | --- | --- | --- | --- | --- | --- | --- | --- | --- |
| 1 | MGCS36044_00966 | MGCS36089_00952 | ***upp*** | uracil phosphoribosyltransferase Upp | -1.06 | -2.25 |  | Core | F |
| 2 | MGCS36044_04004 | MGCS36089_04018 | ***pbp2A*** | multimodular transpeptidase-transglycosylase penicillin-binding protein Pbp2A | -1.19 | -1.24 |  | Core | M |
| 3 | MGCS36044_03032 | MGCS36089_03044 | ***-*** | amino acid ABC transporter permease | -1.27 | -1.54 | Yes | Core | E |
| 4 | MGCS36044_03480 | MGCS36089_03492 | ***ecsB*** | ABC exoprotein transporter permease EcsB | -1.32 | -1.89 | Yes | Core | U |
| 5 | MGCS36044_04274 | MGCS36089_04288 | ***htrA*** | trypsin-like serine protease HtrA | -1.32 | -1.76 |  | Core | O |
| 6 | MGCS36044_01938 | MGCS36089_01926 | ***potD*** | spermidine putrescine ABC transport system substrate-binding protein PotD | -1.35 | -1.31 | Yes | Core | P |
| 7 | MGCS36044_00802 | MGCS36089_00798 | ***oppC_1*** | oligopeptide ABC transporter permease protein OppC | -1.36 | -1.74 | Yes | Core | P |
| 8 | MGCS36044_02044 | MGCS36089_02036 | ***-*** | LolD superfamily ABC transporter ATP-binding component | -1.39 | -1.91 | Yes | Core | V |
| 9 | MGCS36044_01630 | MGCS36089_01618 | ***-*** | LTA synthase family protein | -1.42 | -1.49 |  | Core | M |
| 10 | MGCS36044_02360 | MGCS36089_02352 | ***-*** | amino acid ABC transporter permease | -1.45 | -1.23 | Yes | Core | P |
| 11 | MGCS36044_01522 | MGCS36089_01510 | ***sagF*** | streptolysin S biosynthesis protein SagF | -1.58 | -1.86 |  | Core | K |
| 12 | MGCS36044_01346 | MGCS36089_01334 | ***ppc*** | phosphoenolpyruvate carboxylase Ppc | -1.63 | -1.27 |  | Core | G |
| 13 | MGCS36044_02826 | MGCS36089_02838 | ***-*** | LCP family anionic cell polymer synthesis enzyme | -1.66 | -2.28 |  | Core | M |
| 14 | MGCS36044_01520 | MGCS36089_01508 | ***sagE*** | streptolysin S self-immunity protein SagE | -1.72 | -1.56 |  | Core | V |
| 15 | MGCS36044_00778 | MGCS36089_00774 | ***uppP*** | undecaprenyl pyrophosphate phosphatase UppP | -1.76 | -1.72 |  | Core | M |
| 16 | MGCS36044_02352 | MGCS36089_02344 | ***fieF*** | FieF family cation diffusion facilitator family transporter | -1.85 | -3.16 | Yes | Core | P |
| 17 | MGCS36044_00800 | MGCS36089_00796 | ***oppB_1*** | oligopeptide ABC transporter permease protein OppB | -1.94 | -1.60 | Yes | Core | P |
| 18 | MGCS36044_03682 | MGCS36089_03694 | ***pflB*** | formate C-acetyltransferase | -2.09 | -2.40 |  | Core | C |
| 19 | MGCS36044_00858 | MGCS36089_00844 | ***metN_1*** | methionine ABC transporter ATP-binding protein MetN | -2.14 | -2.65 | Yes | Core | E |
| 20 | MGCS36044_01568 | MGCS36089_01556 | ***murA_1*** | UDP-N-acetylglucosamine 1-carboxyvinyltransferase protein MurA | -2.20 | -3.76 |  | Core | M |
| 21 | MGCS36044_02042 | MGCS36089_02034 | ***acrA*** | AcrA superfamily multidrug efflux pump | -2.37 | -2.36 | Yes | Core | P |
| 22 | MGCS36044_04226 | MGCS36089_04240 | ***sdhB*** | L-serinedehydratase beta subunit SdhB | -2.42 | -1.13 |  | Core | E |
| 23 | MGCS36044_02238 | MGCS36089_02230 | ***cls*** | cardiolipin synthase | -2.69 | -2.44 |  | Core | I |
| 24 | MGCS36044_01636 | MGCS36089_01624 | ***pepT*** | peptidase (T) PepT | -2.73 | -2.67 |  | Core | E |
| 25 | MGCS36044_02046 | MGCS36089_02038 | ***-*** | SalY superfamily ABC transporter permease component | -2.81 | -2.94 | Yes | Core | V |
| 26 | MGCS36044_04090 | MGCS36089_04104 | ***nrdD_2*** | anaerobic ribonucleoside-triphosphate reductase NrdD | -2.92 | -3.66 |  | Core | L |
| 27 | MGCS36044_04082 | MGCS36089_04096 | ***nrdG*** | anaerobic ribonucleoside-triphosphate reductase activating protein NrdG | -2.94 | -3.24 |  | Core | L |
| 28 | MGCS36044_02050 | MGCS36089_02042 | ***ihk*** | TCS signal transduction histidine kinase sensor Ihk | -3.05 | -1.62 |  | Core | T |
| 29 | MGCS36044_01638 | MGCS36089_01626 | ***ebsA*** | EbsA family pore-forming protein | -3.14 | -2.69 |  | Core | M |
| 30 | MGCS36044_02048 | MGCS36089_02040 | ***irr*** | TCS signal transduction DNA-binding response regulator Irr | -3.14 | -2.03 |  | Core | T |
| 31 | MGCS36044_00032 | MGCS36089_00032 | ***plaP*** | amino acid permease PlaP | -3.52 | -3.20 | Yes | Core | E |
| 32 | MGCS36044_01686 | MGCS36089_01674 | ***lspA*** | lipoprotein signal peptidase II LspA | -3.87 | -2.46 |  | Core | M |
| 33 | MGCS36044_02248 | MGCS36089_02240 | ***fhs_1*** | formate--tetrahydrofolate ligase Fhs | -4.03 | -3.35 |  | Core | H |

† Genes are ordered ascendingly based on the locus tag numbers.

∫ Refers to MGCS36044

∫∫ Refers to MGCS36089

†† Genomic location. Core: the core genome, Acc: the accessory genome, ROD: regions of difference.
