## Supplementary material for "Gene contribution of *Streptococcus dysgalactiae* subspecies *equisimilis*, an emerging pathogen, to experimental primate necrotizing myositis": Table S7A

**Table S7A. List of genes shared between MGCS36044 and MGCS36089 where transposon insertion conferred increased fitness.**

| **No.^†^** | **Locus tag 044^∫^** | **Locus tag 089^∫∫^** | | **Gene** | | **Function** | | | **logFC 044^∫^** | | **logFC 089^∫^** | | **Transport** | | **Core/ Acc^††^** | | **COG class** |
| --- | --- | --- | --- | --- | --- | --- | --- | --- | --- | --- | --- | --- | --- | --- | --- | --- | --- |
| 1 | MGCS36044_00516 | | MGCS36089_00516 | | ***emm*** | | cell surface M protein Emm | 4.52 | | 1.88 | |  | | Core | | M | |
| 2 | MGCS36044_00514 | | MGCS36089_00514 | | ***mga*** | | M protein trans-acting positive regulator Mga | 4.50 | | 2.34 | |  | | Core | | K | |
| 3 | MGCS36044_00378 | | MGCS36089_00378 | | ***fbp*** | | secreted fibronectin-binding protein | 2.21 | | 1.68 | |  | | Core | | M | |
| 4 | MGCS36044_01234 | | MGCS36089_01220 | | ***-*** | | glutamine ABC transporter substrate-binding secreted protein | 1.97 | | 1.20 | | Yes | | Core | | E | |
| 5 | MGCS36044_00368 | | MGCS36089_00368 | | ***srtC_1*** | | class C sortase SrtC | 1.54 | | 2.64 | |  | | Core | | M | |
| 6 | MGCS36044_00362 | | MGCS36089_00362 | | ***rofA*** | | pilus transcriptional regulator RofA | 1.51 | | 2.10 | |  | | Core | | K | |
| 7 | MGCS36044_00374 | | MGCS36089_00374 | | ***-*** | | 3' end fragment of disrupted class C sortase | 1.50 | | 2.14 | |  | | Core | | - | |
| 8 | MGCS36044_00364 | | MGCS36089_00364 | | ***-*** | | BP, secreted pilin backbone/major protein | 1.47 | | 2.49 | |  | | Core | | M | |
| 9 | MGCS36044_00370 | | MGCS36089_00370 | | ***srtC_2*** | | truncated class C sortase SrtC | 1.45 | | 1.53 | |  | | Core | | M | |
| 10 | MGCS36044_03076 | | MGCS36089_03088 | | ***-*** | | DUF910 domain-containing protein | 1.41 | | 2.85 | |  | | Core | | S | |
| 11 | MGCS36044_00430 | | MGCS36089_00430 | | ***-*** | | ATP-binding cassette domain-containing protein | 1.11 | | 1.01 | |  | | Core | | L | |
| 12 | MGCS36044_04052 | | MGCS36089_04066 | | ***potE*** | | PotE family amino acid transporter | 1.04 | | 1.11 | | Yes | | Core | | E | |
| 13 | MGCS36044_00380 | | MGCS36089_00380 | | ***-*** | | PAS domain-containing protein | 1.03 | | 1.09 | |  | | Core | | K | |

† Genes are ordered ascendingly based on the locus tag numbers.

∫ Refers to MGCS36044

∫∫ Refers to MGCS36089

†† Genomic location. Core: the core genome, Acc: the accessory genome, ROD: regions of difference.
