## Supplementary material for "Gene contribution of *Streptococcus dysgalactiae* subspecies *equisimilis*, an emerging pathogen, to experimental primate necrotizing myositis": Table S6B

**Table S6B. List of MGCS36089 genes where transposon insertion conferred decreased fitness.**

| **No. ^†^** | **Locus tag** | **Gene** | **Function** | **logFC** | **Transporter** | **Core/ Acc^††^** | **COG class** |
| --- | --- | --- | --- | --- | --- | --- | --- |
| 1 | MGCS36089_02532 | ***-*** | TIGR01906 family membrane protein | -1.02 | Yes | Core | R |
| 2 | MGCS36089_02268 | ***-*** | putative nucleoside ABC transporter | -1.03 | Yes | Core | R |
| 3 | MGCS36089_01648 | ***-*** | YlbF/YmcA family competence regulator | -1.08 |  | Core | T |
| 4 | MGCS36089_02654 | ***glgP*** | maltodextrin phosphorylase protein GlgP | -1.09 |  | Core | G |
| 5 | MGCS36089_04240 | ***sdhB*** | L-serine dehydratase beta subunit SdhB | -1.13 |  | Core | E |
| 6 | MGCS36089_02662 | ***malE*** | maltose/maltodextrin ABC transport system | -1.16 | Yes | Core | G |
| 7 | MGCS36089_00588 | ***rmlB*** | 23S rRNA (guanosine(2251)-2'-O)-methyltransferase RlmB | -1.20 |  | Core | J |
| 8 | MGCS36089_02322 | ***tetR*** | TetR family transcriptional regulator | -1.21 |  | Core | K |
| 9 | MGCS36089_02352 | ***-*** | amino acid ABC transporter permease | -1.23 | Yes | Core | P |
| 10 | MGCS36089_04018 | ***pbp2A*** | multimodular transpeptidase-transglycosylase | -1.24 |  | Core | M |
| 11 | MGCS36089_02134 | ***-*** | helix-turn-helix transcriptional regulator | -1.25 |  | Core | K |
| 12 | MGCS36089_02272 | ***deoC*** | deoxyribose-phosphate aldolase DeoC | -1.25 |  | Core | G |
| 13 | MGCS36089_01334 | ***ppc*** | phosphoenolpyruvate carboxylase Ppc | -1.27 |  | Core | G |
| 14 | MGCS36089_01926 | ***potD*** | spermidine putrescine ABC transport system | -1.31 | Yes | Core | P |
| 15 | MGCS36089_03890 | ***-*** | hypothetical protein | -1.33 |  | Core | S |
| 16 | MGCS36089_02350 | ***-*** | amino acid ABC transporter ATP-binding protein | -1.38 | Yes | Core | E |
| 17 | MGCS36089_02282 | ***ciaH*** | TCS sensor histidine kinase protein CiaH | -1.39 |  | Core | T |
| 18 | MGCS36089_00794 | ***oppA_1*** | oligopeptide ABC transporter substrate-binding | -1.41 | Yes | Core | E |
| 19 | MGCS36089_02348 | ***-*** | amino acid ABC transporter substrate-binding | -1.42 | Yes | Core | E |
| 20 | MGCS36089_00306 | ***adcB*** | metal ABC transporter permease AdcB | -1.44 | Yes | Core | P |
| 21 | MGCS36089_00914 | ***rluB*** | ribosomal large subunit pseudouridine synthase | -1.47 |  | Core | J |
| 22 | MGCS36089_02270 | ***cdd*** | cytidine deaminase Cdd | -1.48 |  | Core | F |
| 23 | MGCS36089_01618 | ***-*** | LTA synthase family protein | -1.49 |  | Core | M |
| 24 | MGCS36089_02304 | ***-*** | UPF0223 family protein | -1.53 |  | Core | S |
| 25 | MGCS36089_03044 | ***-*** | amino acid ABC transporter permease | -1.54 | Yes | Core | E |
| 26 | MGCS36089_01508 | ***sagE*** | streptolysin S self-immunity protein SagE | -1.56 |  | Core | V |
| 27 | MGCS36089_00796 | ***oppB_1*** | oligopeptide ABC transporter permease protein | -1.60 | Yes | Core | P |
| 28 | MGCS36089_02042 | ***ihk*** | TCS signal transduction histidine kinase sensor | -1.62 |  | Core | T |
| 29 | MGCS36089_02316 | ***hsdS*** | type I restriction endonuclease subunit S | -1.65 |  | Core | L |
| 30 | MGCS36089_00774 | ***uppP*** | undecaprenyl pyrophosphate phosphatase UppP | -1.72 |  | Core | M |
| 31 | MGCS36089_03012 | ***-*** | DegV family EDD domain-containing protein | -1.73 |  | Core | I |
| 32 | MGCS36089_00800 | ***oppD_1*** | oligopeptide ABC transporter permease protein | -1.73 | Yes | Core | P |
| 33 | MGCS36089_00798 | ***oppC_1*** | oligopeptide ABC transporter permease protein | -1.74 | Yes | Core | P |
| 34 | MGCS36089_04288 | ***htrA*** | trypsin-like serine protease HtrA | -1.76 |  | Core | O |
| 35 | MGCS36089_00452 | ***-*** | MdlB family ABC transporter ATP-binding/permease | -1.77 | Yes | Core | V |
| 36 | MGCS36089_02310 | ***truB*** | tRNA pseudouridine(55) synthase TruB | -1.78 |  | Core | J |
| 37 | MGCS36089_02974 | ***prfC*** | peptide chain release factor 3 PrfC | -1.79 |  | Core | J |
| 38 | MGCS36089_01510 | ***sagF*** | streptolysin S biosynthesis protein SagF | -1.86 |  | Core | V |
| 39 | MGCS36089_03492 | ***ecsB*** | ABC exoprotein transporter permease EcsB | -1.89 | Yes | Core | U |
| 40 | MGCS36089_02036 | ***-*** | LolD superfamily ABC transporter ATP-binding | -1.91 | Yes | Core | V |
| 41 | MGCS36089_03080 | ***-*** | DUF3165 family protein | -1.92 |  | Core | S |
| 42 | MGCS36089_02040 | ***irr*** | TCS signal transduction DNA-binding response | -2.03 |  | Core | T |
| 43 | MGCS36089_00952 | ***upp*** | uracil phosphoribosyltransferase Upp | -2.25 |  | Core | F |
| 44 | MGCS36089_02208 | ***-*** | apolipoprotein A1/A4/E family protein | -2.28 |  | Core | I |
| 45 | MGCS36089_02838 | ***-*** | LCP family anionic cell polymer synthesis | -2.28 |  | Core | M |
| 46 | MGCS36089_02034 | ***acrA*** | AcrA superfamily multidrug efflux pump | -2.36 | Yes | Core | P |
| 47 | MGCS36089_01712 | ***trxB_1*** | NAD(P)/FAD-dependent oxidoreductase | -2.40 |  | Core | C |
| 48 | MGCS36089_03694 | ***pflB*** | formate C-acetyltransferase | -2.40 |  | Core | C |
| 49 | MGCS36089_03724 | ***-*** | Xre family helix-turn-helix transcriptional | -2.42 |  | ROD.8 | K |
| 50 | MGCS36089_02230 | ***cls*** | cardiolipin synthase | -2.44 |  | Core | I |
| 51 | MGCS36089_01674 | ***lspA*** | lipoprotein signal peptidase II LspA | -2.46 |  | Core | M |
| 52 | MGCS36089_00854 | ***ktrB*** | potassium uptake transporter channel subunit | -2.56 | Yes | Core | P |
| 53 | MGCS36089_00852 | ***ktrA*** | potassium uptake transporter gating subunit | -2.57 | Yes | Core | P |
| 54 | MGCS36089_01730 | ***-*** | DUF1149 domain-containing protein | -2.58 |  | Core | S |
| 55 | MGCS36089_00844 | ***metN_1*** | methionine ABC transporter ATP-binding protein | -2.65 | Yes | Core | E |
| 56 | MGCS36089_01624 | ***pepT*** | peptidase (T) PepT | -2.67 |  | Core | E |
| 57 | MGCS36089_01626 | ***ebsA*** | EbsA family pore-forming protein | -2.69 |  | Core | M |
| 58 | MGCS36089_03830 | ***purR*** | pur operon repressor PurR | -2.74 |  | Core | F |
| 59 | MGCS36089_02038 | ***-*** | SalY superfamily ABC transporter permease | -2.94 | Yes | Core | V |
| 60 | MGCS36089_00846 | ***metP_1*** | methionine ABC transporter permease MetP | -3.01 | Yes | Core | E |
| 61 | MGCS36089_02344 | ***fieF*** | FieF family cation diffusion facilitator family | -3.16 |  | Core | P |
| 62 | MGCS36089_00032 | ***plaP*** | amino acid permease PlaP | -3.20 | Yes | Core | E |
| 63 | MGCS36089_04096 | ***nrdG*** | anaerobic ribonucleoside-triphosphate reductase | -3.24 |  | Core | L |
| 64 | MGCS36089_02240 | ***fhs_1*** | formate--tetrahydrofolate ligase Fhs | -3.35 |  | Core | H |
| 65 | MGCS36089_02680 | ***dltD*** | D-alanyl-lipoteichoic acid biosynthesis protein | -3.65 |  | Core | M |
| 66 | MGCS36089_04104 | ***nrdD_2*** | anaerobic ribonucleoside-triphosphate reductase | -3.66 |  | Core | L |
| 67 | MGCS36089_01556 | ***murA_1*** | UDP-N-acetylglucosamine 1-carboxyvinyltransferase protein MurA | -3.76 |  | Core | M |
| 68 | MGCS36089_02032 | ***-*** | putative lipoprotein | -3.80 |  | Core | R |
| 69 | MGCS36089_00782 | ***sufD*** | Fe-S cluster assembly protein SufD | -3.98 |  | Core | H |
| 70 | MGCS36089_01616 | ***-*** | glycosyltransferase family 1 protein | -4.69 |  | Core | M |
| 71 | MGCS36089_00310 | ***pbp1b*** | bifunctional PG transglycosylase-transpeptidase | -5.47 |  | Core | M |
| 72 | MGCS36089_03330 | ***atoB*** | 3-ketoacyl-CoA thiolase/acetyl-CoA | -7.79 |  | Core | I |

† Genes are ordered ascendingly based on the locus tag numbers.

†† Genomic location. Core: the core genome, Acc: the accessory genome, ROD: regions of difference.
