## Supplementary material for "Gene contribution of *Streptococcus dysgalactiae* subspecies *equisimilis*, an emerging pathogen, to experimental primate necrotizing myositis": Table S6A

**Table S6A. List of MGCS36089 genes where transposon insertion conferred increased fitness**

| **No.^†^** | **Locus tag** | **Gene** | **Function** | **logFC** | **Transporter** | **Core/ Acc^††^** | **COG class** |
| --- | --- | --- | --- | --- | --- | --- | --- |
| 1 | MGCS36089_03516 | ***manM*** | PTS transporter mannose-specific IIC component | 4.15 | Yes | Core | G |
| 2 | MGCS36089_03514 | ***manL*** | PTS transporter mannose-specific IIB & IIA | 3.71 | Yes | Core | G |
| 3 | MGCS36089_03518 | ***manN*** | PTS transporter mannose-specific IID component | 3.55 | Yes | Core | G |
| 4 | MGCS36089_03086 | ***glcK*** | glucokinase GlcK | 3.10 |  | Core | G |
| 5 | MGCS36089_03088 | ***-*** | DUF910 domain-containing protein | 2.85 |  | Core | S |
| 6 | MGCS36089_00368 | ***srtC_1*** | class C sortase SrtC | 2.64 |  | Core | M |
| 7 | MGCS36089_00364 | ***-*** | secreted pilin backbone/major protein. | 2.49 |  | Core | M |
| 8 | MGCS36089_01026 | ***-*** | hypothetical protein. | 2.46 |  | Core | R |
| 9 | MGCS36089_00514 | ***mga*** | M protein trans-acting positive regulator Mga | 2.34 |  | Core | K |
| 10 | MGCS36089_00332 | ***comYG*** | competence system protein ComYG | 2.23 |  | Core | U |
| 11 | MGCS36089_00374 | ***-*** | 3' end fragment of disrupted class C sortase | 2.14 |  | Core | - |
| 12 | MGCS36089_00362 | ***rofA*** | pilus transcriptional regulator RofA | 2.10 |  | Core | K |
| 13 | MGCS36089_02888 | ***-*** | DUF1294 domain-containing protein | 1.94 |  | Core | S |
| 14 | MGCS36089_00504 | ***-*** | M42 family metallopeptidase | 1.94 |  | Core | E |
| 15 | MGCS36089_00156 | ***adhP*** | alcohol dehydrogenase AdhP | 1.92 |  | Core | G |
| 16 | MGCS36089_00516 | ***emm*** | cell surface M protein Emm. | 1.88 |  | Core | M |
| 17 | MGCS36089_00378 | ***fbp*** | secreted fibronectin-binding protein. | 1.68 |  | Core | M |
| 18 | MGCS36089_01218 | ***-*** | glutamine ABC transporter permease | 1.64 | Yes | Core | E |
| 19 | MGCS36089_00372 | ***-*** | secreted pilin minor/ancillary protein. | 1.58 |  | Core | M |
| 20 | MGCS36089_01222 | ***glnQ_1*** | glutamine ABC transporter ATPase GlnQ | 1.56 | Yes | Core | E |
| 21 | MGCS36089_00370 | ***srtC_2*** | class C sortase SrtC | 1.53 |  | Core | M |
| 22 | MGCS36089_00444 | ***flaR*** | DNA topology modulation protein | 1.40 |  | Core | L |
| 23 | MGCS36089_00416 | ***-*** | hypothetical protein | 1.34 |  | Core | S |
| 24 | MGCS36089_03604 | ***-*** | aldo/keto reductase | 1.31 |  | Core | S |
| 25 | MGCS36089_00638 | ***-*** | conjugal transfer protein | 1.26 |  | ROD.3 | R |
| 26 | MGCS36089_04206 | ***-*** | DUF1700 domain-containing protein | 1.25 |  | Core | S |
| 27 | MGCS36089_03568 | ***-*** | GlsB/YeaQ/YmgE family stress response membrane | 1.23 |  | Core | D |
| 28 | MGCS36089_01220 | ***-*** | glutamine ABC transporter substrate-binding. | 1.20 | Yes | Core | E |
| 29 | MGCS36089_00456 | ***-*** | aminoglycoside 6-adenylyltransferase | 1.20 |  | Core | V |
| 30 | MGCS36089_04166 | ***-*** | sigma-70 family RNA polymerase sigma factor | 1.18 |  | ROD.9 | K |
| 31 | MGCS36089_04070 | ***hutG*** | formiminoglutamase HutG | 1.13 |  | Core | E |
| 32 | MGCS36089_04066 | ***potE*** | PotE family amino acid transporter | 1.11 | Yes | Core | E |
| 33 | MGCS36089_01692 | ***-*** | RND family transporter membrane fusion protein | 1.09 | Yes | Core | M |
| 34 | MGCS36089_00380 | ***-*** | PAS domain-containing protein | 1.09 |  | Core | K |
| 35 | MGCS36089_01104 | ***-*** | hypothetical protein | 1.03 |  | Core | S |
| 36 | MGCS36089_01158 | ***-*** | multidrug efflux MFS transporter | 1.03 | Yes | Core | R |
| 37 | MGCS36089_03670 | ***-*** | hypothetical protein | 1.03 |  | Core | S |
| 38 | MGCS36089_00430 | ***-*** | ATP-binding cassette domain-containing protein | 1.01 |  | Core | L |

† Genes are ordered ascendingly based on the locus tag numbers.

††Genomic location. Core: the core genome, Acc: the accessory genome, ROD: regions of difference.
