## Supplementary material for "Gene contribution of *Streptococcus dysgalactiae* subspecies *equisimilis*, an emerging pathogen, to experimental primate necrotizing myositis": Table S5B

**Table S5B. List of MGCS36044 genes where transposon insertion conferred decreased fitness**

| **No.^†^** | **Locus tag** | | **Gene** | **Function** | **logFC** | **Transporter** | **Core/**  **Acc^††^** | **COG**  **class** |
| --- | --- | --- | --- | --- | --- | --- | --- | --- |
| 1 | MGCS36044_02082 | ***femX*** | | FemABX-like family peptidoglycan interpeptide bridge formation enzyme | -1.05 |  | Core | M |
| 2 | MGCS36044_02480 | ***-*** | | AgaB family mannose/fructose/N-acetylgalactosamine-specific component IIB | -1.06 |  | Core | G |
| 3 | MGCS36044_00966 | ***upp*** | | uracil phosphoribosyltransferase Upp | -1.06 |  | Core | F |
| 4 | MGCS36044_02328 | ***-*** | | ABC transporter ATP-binding protein LolD-like | -1.15 | Yes | Core | M |
| 5 | MGCS36044_03634 | ***scrR*** | | sucrose operon repressor ScrR | -1.19 |  | Core | K |
| 6 | MGCS36044_04004 | ***pbp2A*** | | multimodular transpeptidase-transglycosylase penicillin-binding protein Pbp2A | -1.19 |  | Core | M |
| 7 | MGCS36044_03222 | ***-*** | | hypothetical protein | -1.24 |  | Core | S |
| 8 | MGCS36044_03032 | ***-*** | | amino acid ABC transporter permease | -1.27 | Yes | Core | E |
| 9 | MGCS36044_03816 | ***prgA*** | | surface exclusion domain-containing secreted protein PrgA | -1.29 |  | Core | M |
| 10 | MGCS36044_03480 | ***ecsB*** | | ABC exoprotein transporter permease EcsB | -1.32 | Yes | Core | U |
| 11 | MGCS36044_04274 | ***htrA*** | | trypsin-like serine protease HtrA | -1.32 |  | Core | O |
| 12 | MGCS36044_04252 | ***glcU*** | | glucose uptake permease GlcU | -1.33 | Yes | Core | G |
| 13 | MGCS36044_00612 | ***-*** | | DNA-binding protein | -1.34 |  | ROD.3 | R |
| 14 | MGCS36044_01938 | ***potD*** | | spermidine putrescine ABC transport system substrate-binding protein PotD | -1.35 | Yes | Core | P |
| 15 | MGCS36044_00802 | ***oppC_1*** | | oligopeptide ABC transporter permease protein OppC | -1.36 | Yes | Core | P |
| 16 | MGCS36044_02186 | ***gid*** | | tRNA (uracil-5-)-methyltransferase/glucose inhibited division protein (A) GidA | -1.37 |  | Core | J |
| 17 | MGCS36044_02044 | ***-*** | | LolD superfamily ABC transporter ATP-binding component | -1.39 | Yes | Core | V |
| 18 | MGCS36044_01630 | ***-*** | | LTA synthase family protein | -1.42 |  | Core | M |
| 19 | MGCS36044_02360 | ***-*** | | amino acid ABC transporter permease | -1.45 | Yes | Core | P |
| 20 | MGCS36044_02432 | ***-*** | | LysR family transcriptional regulator | -1.50 |  | Core | K |
| 21 | MGCS36044_02862 | ***murA_2*** | | UDP-N-acetylglucosamine 1-carboxyvinyltransferase MurA | -1.52 |  | Core | M |
| 22 | MGCS36044_02562 | ***phnK*** | | PhnK family ABC transporter ATPase component | -1.53 | Yes | Core | P |
| 23 | MGCS36044_01522 | ***sagF*** | | streptolysin S biosynthesis protein SagF | -1.58 |  | Core | K |
| 24 | MGCS36044_01346 | ***ppc*** | | phosphoenolpyruvate carboxylase Ppc | -1.63 |  | Core | G |
| 25 | MGCS36044_01932 | ***potA*** | | spermidine putrescine ABC transport system ATP-binding protein PotA | -1.64 | Yes | Core | P |
| 26 | MGCS36044_02826 | ***-*** | | LCP family anionic cell polymer synthesis enzyme | -1.66 |  | Core | M |
| 27 | MGCS36044_00834 | ***-*** | | cysteine hydrolase | -1.67 |  | Core | E |
| 28 | MGCS36044_04172 | ***-*** | | MFS transporter | -1.67 | Yes | ROD.9 | R |
| 29 | MGCS36044_02588 | ***perM*** | | PerM family predicted purR regulated permease | -1.70 | Yes | Core | D |
| 30 | MGCS36044_01520 | ***sagE*** | | streptolysin S self-immunity protein SagE | -1.72 |  | Core | V |
| 31 | MGCS36044_02294 | ***pepN*** | | lysyl aminopeptidase/alanine aminopeptidase PepN | -1.74 |  | Core | E |
| 32 | MGCS36044_00778 | ***uppP*** | | undecaprenyl pyrophosphate phosphatase UppP | -1.76 |  | Core | M |
| 33 | MGCS36044_00304 | ***adcC*** | | metal ABC transporter ATP-binding protein AdcC | -1.76 | Yes | Core | P |
| 34 | MGCS36044_02352 | ***fieF*** | | FieF family cation diffusion facilitator family transporter | -1.85 | Yes | Core | P |
| 35 | MGCS36044_01254 | ***-*** | | Cof-type HAD-IIB family hydrolase | -1.89 |  | Core | Q |
| 36 | MGCS36044_01088 | ***cvfB*** | | S1 RNA-binding domain-containing protein CvfB | -1.91 |  | Core | K |
| 37 | MGCS36044_00800 | ***oppB_1*** | | oligopeptide ABC transporter permease protein OppB | -1.94 | Yes | Core | P |
| 38 | MGCS36044_02430 | ***-*** | | hypothetical protein | -1.96 |  | Core | S |
| 39 | MGCS36044_03482 | ***ecsA*** | | ABC exoprotein transporter ATPase EcsA | -2.00 | Yes | Core | V |
| 40 | MGCS36044_01688 | ***rluD*** | | ribosomal large subunit pseudouridine synthase RluD | -2.01 |  | Core | J |
| 41 | MGCS36044_00360 | ***hslO*** | | Hsp33 family molecular chaperone HslO | -2.04 |  | Core | O |
| 42 | MGCS36044_00836 | ***rsfS*** | | ribosome silencing factor RsfS | -2.05 |  | Core | H |
| 43 | MGCS36044_03682 | ***pflB*** | | formate C-acetyltransferase | -2.09 |  | Core | C |
| 44 | MGCS36044_04084 | ***-*** | | putative acetyltransferase | -2.12 |  | Core | K |
| 45 | MGCS36044_00858 | ***metN_1*** | | methionine ABC transporter ATP-binding protein MetN | -2.14 | Yes | Core | E |
| 46 | MGCS36044_02380 | ***-*** | | ABC transporter permease | -2.16 | Yes | Core | R |
| 47 | MGCS36044_01568 | ***murA_1*** | | UDP-N-acetylglucosamine 1-carboxyvinyltransferase protein MurA | -2.20 |  | Core | M |
| 48 | MGCS36044_00874 | ***htpX*** | | zinc metalloprotease HtpX | -2.20 |  | Core | O |
| 49 | MGCS36044_02854 | ***fadR*** | | FadR family DNA-binding transcriptional regulator | -2.31 |  | Core | K |
| 50 | MGCS36044_02042 | ***acrA*** | | AcrA superfamily multidrug efflux pump | -2.37 | Yes | Core | P |
| 51 | MGCS36044_01762 | ***sptR*** | | SptR-like TCS DNA-binding response regulator | -2.38 |  | Core | T |
| 52 | MGCS36044_02366 | ***lepB_1*** | | signal peptidase I | -2.38 |  | Core | U |
| 53 | MGCS36044_02566 | ***-*** | | ABC transporter substrate binding component | -2.39 | Yes | Core | E |
| 54 | MGCS36044_04226 | ***sdhB*** | | L-serinedehydratase beta subunit SdhB | -2.42 |  | Core | E |
| 55 | MGCS36044_04254 | ***guaB*** | | IMP dehydrogenase GuaB | -2.44 |  | Core | F |
| 56 | MGCS36044_01912 | ***dacA_3*** | | secreted D,D-carboxypeptidase penicillin-binding protein DacA | -2.47 |  | Core | M |
| 57 | MGCS36044_01760 | ***-*** | | GTP pyrophosphokinase family protein | -2.48 |  | Core | M |
| 58 | MGCS36044_00772 | ***-*** | | ABC amino acid transporter ATP-binding protein | -2.56 |  | Core | E |
| 59 | MGCS36044_02304 | ***ptsC*** | | phosphate ABC transporter permease PstC | -2.68 | Yes | Core | P |
| 60 | MGCS36044_02238 | ***cls*** | | cardiolipin synthase | -2.69 | Yes | Core | I |
| 61 | MGCS36044_01636 | ***pepT*** | | peptidase (T) PepT | -2.73 |  | Core | E |
| 62 | MGCS36044_02046 | ***-*** | | SalY superfamily ABC transporter permease component | -2.81 | Yes | Core | V |
| 63 | MGCS36044_00274 | ***-*** | | helix-turn-helix transcriptional regulator | -2.89 |  | ROD.1 | K |
| 64 | MGCS36044_04090 | ***nrdD_2*** | | anaerobic ribonucleoside-triphosphate reductase NrdD | -2.92 |  | Core | L |
| 65 | MGCS36044_04082 | ***nrdG*** | | anaerobic ribonucleoside-triphosphate reductase activating protein NrdG | -2.94 |  | Core | L |
| 66 | MGCS36044_02908 | ***prsA*** | | peptidylprolyl isomerase lipoprotein PrsA | -3.00 |  | Core | O |
| 67 | MGCS36044_00384 | ***-*** | | PTS sugar transporter subunit IIC | -3.01 | Yes | Core | G |
| 68 | MGCS36044_02050 | ***ihk*** | | TCS signal transduction histidine kinase sensor Ihk | -3.05 |  | Core | T |
| 69 | MGCS36044_01638 | ***ebsA*** | | EbsA family pore-forming protein | -3.14 |  | Core | M |
| 70 | MGCS36044_02048 | ***irr*** | | TCS signal transduction DNA-binding response regulator Irr | -3.14 |  | Core | T |
| 71 | MGCS36044_01692 | ***-*** | |  | -3.15 |  | Core | S |
| 72 | MGCS36044_03944 | ***perR*** | | peroxide-responsive transcriptional repressor PerR | -3.41 |  | Core | K |
| 73 | MGCS36044_04276 | ***parB*** | | chromosome partitioning protein ParB | -3.42 |  | Core | D |
| 74 | MGCS36044_00032 | ***plaP*** | | amino acid permease PlaP | -3.52 | Yes | Core | E |
| 75 | MGCS36044_02128 | ***guaA*** | | glutamine-hydrolyzing GMP synthase | -3.53 |  | ROD.5 | F |
| 76 | MGCS36044_01302 | ***lgt*** | | prolipoprotein diacylglyceryl transferase Lgt | -3.58 |  | Core | M |
| 77 | MGCS36044_04092 | ***-*** | | DUF2079 domain-containing protein | -3.66 |  | Core | M |
| 78 | MGCS36044_03542 | ***-*** | | PfpI family predicted protease/amidase | -3.82 |  | Core | R |
| 79 | MGCS36044_00740 | ***tig*** | | trigger factor molecular chaperone Tig | -3.84 |  | Core | O |
| 80 | MGCS36044_01686 | ***lspA*** | | lipoprotein signal peptidase II LspA | -3.87 |  | Core | M |
| 81 | MGCS36044_01632 | ***galE*** | | UDP-glucose 4-epimerase GalE | -3.97 |  | Core | M |
| 82 | MGCS36044_02248 | ***fhs_1*** | | formate--tetrahydrofolate ligase Fhs | -4.03 |  | Core | H |
| 83 | MGCS36044_00792 | ***sufB*** | | Fe-S cluster assembly protein SufB | -5.32 |  | Core | H |

† Genes are ordered ascendingly based on the locus tag numbers.

††Genomic location. Core: the core genome, Acc: the accessory genome, ROD: regions of difference.
