## Supplementary material for "Gene contribution of *Streptococcus dysgalactiae* subspecies *equisimilis*, an emerging pathogen, to experimental primate necrotizing myositis": Table S5A

**Table S5A. List of MGCS36044 genes where transposon insertion conferred increased fitness**

| **No.^†^** | **Locus tag** | **Gene** | **Function** | **logFC** | **Transporter** | **Core/ Acc^††^** | **COG class** |
| --- | --- | --- | --- | --- | --- | --- | --- |
| 1 | MGCS36044_00516 | ***emm*** | cell surface M protein Emm | 4.52 |  | Core | M |
| 2 | MGCS36044_00514 | ***mga*** | M protein trans-acting positive regulator Mga | 4.50 |  | Core | K |
| 3 | MGCS36044_00378 | ***fbp*** | secreted fibronectin-binding protein | 2.21 |  | Core | M |
| 4 | MGCS36044_00322 | ***comYB*** | competence system type II secretion system protein ComYB | 2.19 |  | Core | U |
| 5 | MGCS36044_01234 | **-** | glutamine ABC transporter substrate-binding secreted protein | 1.97 | Yes | Core | E |
| 6 | MGCS36044_00366 | **-** | secreted pilin AP2, minor/ancillary protein 2 | 1.73 |  | Core | M |
| 7 | MGCS36044_03392 | ***glpO*** | type 1 glycerol-3-phosphate oxidase GlpO | 1.72 | Yes | Core | C |
| 8 | MGCS36044_00368 | ***srtC_1*** | class C sortase SrtC | 1.54 |  | Core | M |
| 9 | MGCS36044_00362 | ***rofA*** | pilus transcriptional regulator RofA | 1.51 |  | Core | K |
| 10 | MGCS36044_00438 | **-** | bacteriocin immunity protein | 1.50 |  | Core | V |
| 11 | MGCS36044_00374 | **-** | Disrupted class C sortase | 1.50 |  | Core | - |
| 12 | MGCS36044_00476 | **-** | DUF771 domain-containing protein | 1.48 |  | ROD.2 | S |
| 13 | MGCS36044_00364 | **-** | BP, secreted pilin backbone/major protein | 1.47 |  | Core | M |
| 14 | MGCS36044_03452 | ***sipA*** | signal peptidase I SipA | 1.45 |  | Core | U |
| 15 | MGCS36044_00370 | ***srtC_2*** | truncated class C sortase SrtC | 1.45 |  | Core | - |
| 16 | MGCS36044_00376 | ***srtC_3*** | class C sortase SrtC | 1.42 |  | Core | M |
| 17 | MGCS36044_00532 | ***dexB*** | glucan 1,6-alpha-glucosidase DexB | 1.41 |  | Core | C |
| 18 | MGCS36044_03076 | **-** | DUF910 domain-containing protein | 1.41 |  | Core | S |
| 19 | MGCS36044_00280 | **-** | XRE family ImmR-like transcriptional regulator | 1.40 |  | ROD.1 | K |
| 20 | MGCS36044_03948 | **-** | hypothetical protein | 1.28 |  | Core | S |
| 21 | MGCS36044_01016 | ***sdpI*** | SdpI family immunity protein | 1.25 |  | Core | V |
| 22 | MGCS36044_00386 | **-** | toxic anion resistance protein, tellurite resistance protein | 1.23 |  | Core | P |
| 23 | MGCS36044_00666 | **-** | hypothetical protein | 1.20 |  | ROD.3 | S |
| 24 | MGCS36044_02858 | **-** | sugar kinase | 1.14 |  | Core | R |
| 25 | MGCS36044_00430 | **-** | ATP-binding cassette domain-containing protein | 1.11 |  | Core | L |
| 26 | MGCS36044_03976 | ***metC*** | disrupted cystathionine beta-lyase encoding gene | 1.10 |  | Core | - |
| 27 | MGCS36044_00432 | **-** | disrupted cyclic nucleotide-binding domain-containing protein | 1.09 |  | Core | - |
| 28 | MGCS36044_04052 | ***potE*** | PotE family amino acid transporter | 1.04 | Yes | Core | E |
| 29 | MGCS36044_01516 | ***sagC*** | streptolysin S biosynthesis protein SagC | 1.03 |  | Core | K |
| 30 | MGCS36044_00380 | **-** | PAS domain-containing protein | 1.03 |  | Core | K |
| 31 | MGCS36044_00450 | **-** | hypothetical protein | 1.02 |  | Core | S |
| 32 | MGCS36044_03344 | ***pepC*** | aminopeptidase (A) PepC | 1.01 | Yes | Core | E |
| 33 | MGCS36044_03972 | ***ulaA*** | ascorbate-specific PTS transporter EIIC protein UlaA | 1.01 | Yes | Core | G |
| 33 | MGCS36044_03426 | **lacB** | galactose-6-phosphate isomerase subunit LacB | 1.00 |  | Core | G |

† Genes are ordered ascendingly based on the locus tag numbers.

†† Genomic location. Core: the core genome, Acc: the accessory genome, ROD: regions of difference.
