## Supplementary material for "Gene contribution of *Streptococcus dysgalactiae* subspecies *equisimilis*, an emerging pathogen, to experimental primate necrotizing myositis": Table S4D

**Table S4D. Conditionally essential genes in MGCS36044 and MGCS36089 *in vivo*, but not *in vitro*.**

| **No. ^†^** | **Locus tag 044^∫^** | **Locus tag 089^∫∫^** | **Gene** | **Function** | **COG class** | **COG description** | **Core/ Acc^††^** | **Pro/ RNA^‡‡^** |
| --- | --- | --- | --- | --- | --- | --- | --- | --- |
| 1 | MGCS36044_00784 | MGCS36089_00780 | ***sufC*** | Fe-S cluster assembly ATPase SufC | H | METABOLISM; coenzyme transport and metabolism | Core | Pro |
| 2 | MGCS36044_00792 | MGCS36089_00788 | ***sufB*** | Fe-S cluster assembly protein SufB | H | METABOLISM; coenzyme transport and metabolism | Core | Pro |
| 3 | MGCS36044_01302 | MGCS36089_01290 | ***lgt*** | prolipoprotein diacylglyceryl transferase Lgt | M | CELLULAR PROCESSES AND SIGNALING; cell wall/membrane/envelope biogenesis | Core | Pro |
| 4 | MGCS36044_01446 | MGCS36089_01434 | ***whiA*** | cell division involved DNA-binding protein WhiA | D | CELLULAR PROCESSES AND SIGNALING; cell cycle control, cell division, chromosome partitioning | Core | Pro |
| 5 | MGCS36044_01568 | MGCS36089_01556 | ***murA_1*** | UDP-N-acetylglucosamine 1-carboxyvinyltransferase protein MurA | M | CELLULAR PROCESSES AND SIGNALING; cell wall/membrane/envelope biogenesis | Core | Pro |
| 6 | MGCS36044_01632 | MGCS36089_01620 | ***galE*** | UDP-glucose 4-epimerase GalE | M | CELLULAR PROCESSES AND SIGNALING; cell wall/membrane/envelope biogenesis | Core | Pro |
| 7 | MGCS36044_01724 | MGCS36089_01712 | ***trxB_1*** | NAD(P)/FAD-dependent oxidoreductase | H | METABOLISM; coenzyme transport and metabolism | Core | Pro |

† Genes are ordered ascendingly based on the locus tag numbers.

**^∫^** Refers to MGCS36044

**^∫∫^** Refers to MGCS36089

††Genomic location. Core: the core genome, Acc: the accessory genome, ROD: regions of difference.

‡‡ Pro: protein.
