## Supplementary material for "Gene contribution of *Streptococcus dysgalactiae* subspecies *equisimilis*, an emerging pathogen, to experimental primate necrotizing myositis": Table S4C

**Table S4C. Conditional essential genes in MGCS36089**

| **No. ^†^** | **Locus tag** | **Gene** | **Virulence^∫^** | **Function** | **COG class** | **COG description** | **Core/ Acc^††^** | | **Pro/ RNA^‡‡^** |  |  |
| --- | --- | --- | --- | --- | --- | --- | --- | --- | --- | --- | --- |
| 1 | MGCS36089_00112 | ***acpP_1*** |  | acyl carrier protein AcpP | I | METABOLISM; lipid metabolism | Core | | Pro |  |  |
| 2 | MGCS36089_00194 | ***rpsZ*** |  | type Z 30S ribosomal S14 protein RpsZ | J | INFORMATION STORAGE AND PROCESSING; translation, ribosomal structure and biogenesis | Core | | Pro |  |  |
| 3 | MGCS36089_00214 | ***rpmJ*** |  | 50S ribosomal L36 protein RpmJ | J | INFORMATION STORAGE AND PROCESSING; translation, ribosomal structure and biogenesis | Core | | Pro |  |  |
| 4 | MGCS36089_00726 | ***thiD*** |  | bifunctional hydroxymethylpyrimidine kinase/phosphomethylpyrimidine kinase PdxK | H | METABOLISM; coenzyme transport and metabolism | Core | | Pro |  |  |
| 5 | MGCS36089_00782 | ***sufD*** |  | Fe-S cluster assembly protein SufD | O | CELLULAR PROCESSES AND SIGNALING; post-translational modification, protein turnover, and chaperones | Core | | Pro |  |  |
| 6 | MGCS36089_00866 | ***covS*** | Virulence | TCS sensor kinase CovS | T | CELLULAR PROCESSES AND SIGNALING; signal transduction mechanisms | Core | | Pro |  |  |
| 7 | MGCS36089_00910 | ***scp1*** |  | segregation/condensation complex subunit (A) ScpA/Scp1 | D | CELLULAR PROCESSES AND SIGNALING; cell cycle control, cell division, chromosome partitioning | Core | | Pro |  |  |
| 8 | MGCS36089_01054 | ***mtsB*** |  | metal ABC transporter ATP-binding protein MtsB | P | METABOLISM; inorganic ion transport and metabolism | Core | | Pro |  |  |
| 9 | MGCS36089_01160 | ***secG*** |  | preprotein translocase subunit SecG | U | CELLULAR PROCESSES AND SIGNALING; Intracellular trafficking, secretion, and vesicular transport | Core | | Pro |  |  |
| 10 | MGCS36089_01404 | ***-*** |  | IS3 family transposase | |  | | Core | | | Pro |
| 11 | MGCS36089_01538 | ***atpE*** |  | ATP synthase C subunit AtpE | C | METABOLISM; energy production and conversion | Core | | Pro |  |  |
| 12 | MGCS36089_01586 | ***rpsU*** |  | 30S ribosomal S21 protein RpsU | J | INFORMATION STORAGE AND PROCESSING; translation, ribosomal structure and biogenesis | Core | | Pro |  |  |
| 13 | MGCS36089_01676 | ***rluD*** |  | ribosomal large subunit pseudouridine synthase RluD | J | INFORMATION STORAGE AND PROCESSING; translation, ribosomal structure and biogenesis | Core | | Pro |  |  |
| 14 | MGCS36089_01704 | ***-*** |  | KH domain-containing protein | L | INFORMATION STORAGE AND PROCESSING; replication, recombination and repair | Core | | Pro |  |  |
| 15 | MGCS36089_02202 | ***-*** |  | DUF1836 domain-containing protein | S | POORLY CHARACTERIZED; function unknown | Core | | Pro |  |  |
| 16 | MGCS36089_02560 | ***-*** |  | hypothetical protein | S | POORLY CHARACTERIZED; function unknown | Core | | Pro |  |  |
| 17 | MGCS36089_02682 | ***dltC*** |  | D-alanine--poly(phosphoribitol) ligase subunit DltC | M | CELLULAR PROCESSES AND SIGNALING; cell wall/membrane/envelope biogenesis | Core | | Pro |  |  |
| 18 | MGCS36089_02718 | ***-*** |  | DUF4044 domain-containing protein | S | POORLY CHARACTERIZED; function unknown | Core | | Pro |  |  |
| 19 | MGCS36089_02810 | ***-*** |  | MdlB family multidrug ABC transporter ATPase and permease | V | CELLULAR PROCESSES AND SIGNALING; defense mechanisms | ROD.7 | | Pro |  |  |
| 20 | MGCS36089_02812 | ***-*** |  | MdlB family multidrug ABC transporter ATPase and permease | V | CELLULAR PROCESSES AND SIGNALING; defense mechanisms | ROD.7 | | Pro |  |  |
| 21 | MGCS36089_02814 | ***ecfA2*** |  | EcfA2 family ECF transporter ATPase | G | METABOLISM; carbohydrate transport and metabolism | ROD.7 | | Pro |  |  |
| 22 | MGCS36089_02816 | ***ecfT*** |  | ECF transporter transmembrane protein EcfT | P | METABOLISM; inorganic ion transport and metabolism | ROD.7 | | Pro |  |  |
| 23 | MGCS36089_02818 | ***-*** |  | ECF transporter S component | P | METABOLISM; inorganic ion transport and metabolism | ROD.7 | | Pro |  |  |
| 24 | MGCS36089_02820 | ***-*** |  | TetR/AcrR family transcriptional regulator | K | INFORMATION STORAGE AND PROCESSING; transcription | ROD.7 | | Pro |  |  |
| 25 | MGCS36089_03380 | ***ftsL*** |  | cell division protein FtsL | D | CELLULAR PROCESSES AND SIGNALING; cell cycle control, cell division, chromosome partitioning | Core | | Pro |  |  |
| 26 | MGCS36089_03474 | ***rbfA*** |  | 30S ribosome-binding factor RbfA | J | INFORMATION STORAGE AND PROCESSING; translation, ribosomal structure and biogenesis | Core | | Pro |  |  |
| 27 | MGCS36089_03504 | ***tsaE*** |  | tRNA (adenosine(37)-N6)-threonylcarbamoyltransferase TsaE | J | INFORMATION STORAGE AND PROCESSING; translation, ribosomal structure and biogenesis | Core | | Pro |  |  |
| 28 | MGCS36089_03538 | ***fabG_2*** |  | 3-ketoacyl-(acyl-carrier-protein) reductase protein FabG | I | METABOLISM; lipid metabolism | Core | | Pro |  |  |
| 29 | MGCS36089_03544 | ***acpP_2*** |  | acyl carrier protein AcpP | I | METABOLISM; lipid metabolism | Core | | Pro |  |  |
| 30 | MGCS36089_03546 | ***fabH*** |  | 3-oxoacyl-[acyl-carrier-protein] synthase protein FabH | I | METABOLISM; lipid metabolism | Core | | Pro |  |  |
| 31 | MGCS36089_03596 | ***alaT*** |  | AlaT family aminotransferase | E | METABOLISM; amino acid transport and metabolism | Core | | Pro |  |  |
| 32 | MGCS36089_03660 | ***-*** |  | CorA family divalent cation transport protein | P | METABOLISM; inorganic ion transport and metabolism | Core | | Pro |  |  |
| 33 | MGCS36089_03844 | ***rsmA*** |  | 16S rRNA (adenine(1518)-N(6)/adenine(1519)-N(6))- dimethyltransferase RsmA | J | INFORMATION STORAGE AND PROCESSING; translation, ribosomal structure and biogenesis | Core | | Pro |  |  |
| 34 | MGCS36089_03872 | ***jag*** |  | RNA-binding protein Jag | L | INFORMATION STORAGE AND PROCESSING; replication, recombination and repair | Core | | Pro |  |  |
| 35 | MGCS36089_04106 | ***-*** |  | DUF2079 domain-containing protein | S | POORLY CHARACTERIZED; function unknown | Core | | Pro |  |  |
| 36 | MGCS36089_04198 | ***-*** |  | DUF368 domain-containing protein | S | POORLY CHARACTERIZED; function unknown | Core | | Pro |  |  |
| 37 | MGCS36089_04256 | ***rodZ*** |  | cytoskeltal protein RodZ | O | CELLULAR PROCESSES AND SIGNALING; post-translational modification, protein turnover, and chaperones | Core | | Pro |  |  |
| 38 | MGCS36089_04264 | ***recF*** |  | DNA replication/repair protein RecF | L | INFORMATION STORAGE AND PROCESSING; DNA replication, recombination and repair | Core | | Pro |  |  |
| 39 | MGCS36089_04274 | ***uup*** |  | Uup family ATPase components of ABC transporters with duplicated ATPase domains | L | INFORMATION STORAGE AND PROCESSING; DNA replication, recombination and repair | Core | | Pro |  |  |

† Genes are ordered ascendingly based on the locus tag numbers.

‡ Putative virulence genes (Eraso et al., 2024, *mBio*).

††Genomic location. Core: the core genome, Acc: the accessory genome, ROD: regions of difference.

‡‡ Pro: protein.
