## Supplementary material for "Gene contribution of *Streptococcus dysgalactiae* subspecies *equisimilis*, an emerging pathogen, to experimental primate necrotizing myositis": Table S4B

**Table S4B. Conditional essential genes in MGCS36044**

| **No. ^†^** | **Locus tag** | **Gene** | **Virulence^‡^** | **Function** | **COG class** | **COG description** | | | | **Core/ ACC^††^** | **Pro/ RNA^‡‡^** |
| --- | --- | --- | --- | --- | --- | --- | --- | --- | --- | --- | --- |
| 1 | MGCS36044_00140 | ***purB*** |  | adenylosuccinate lyase PurB | F | METABOLISM; nucleotide transport and metabolism | | | | core | Pro |
| 2 | MGCS36044_00146 | ***ruvB*** |  | Holliday junction branch migration DNA helicase RvuB | L | INFORMATION STORAGE AND PROCESSING; DNA replication, recombination and repair | | | | core | Pro |
| 3 | MGCS36044_00164 | ***-*** |  | IS30 family transposase |  |  | | | | core | Pro |
| 4 | MGCS36044_00202 | ***rpsE*** |  | 30S ribosomal S5 protein RpsE | J | INFORMATION STORAGE AND PROCESSING; translation, ribosomal structure and biogenesis | | | | core | Pro |
| 5 | MGCS36044_00206 | ***rplO*** |  | 50S ribosomal L15 protein RplO | J | INFORMATION STORAGE AND PROCESSING; translation, ribosomal structure and biogenesis | | | | core | Pro |
| 6 | MGCS36044_00210 | ***adk*** |  | adenylate kinase protein Adk | F | METABOLISM; nucleotide transport and metabolism | | | | core | Pro |
| 7 | MGCS36044_00544 | ***cdsA*** |  | phosphatidate cytidylyltransferase CdsA | I | METABOLISM; lipid metabolism | | | | core | Pro |
| 8 | MGCS36044_00740 | ***tig*** |  | trigger factor molecular chaperone Tig | O | CELLULAR PROCESSES AND SIGNALING; post-translational modification, protein turnover, and chaperones | | | | core | Pro |
| 9 | MGCS36044_00760 | ***rpmB*** |  | 50S ribosomal L28 protein RpmB | J | INFORMATION STORAGE AND PROCESSING; translation, ribosomal structure and biogenesis | | | | core | Pro |
| 10 | MGCS36044_00762 | ***-*** |  | IS1548 family transposase | L | INFORMATION STORAGE AND PROCESSING; DNA replication, recombination and repair | | | | core | Pro |
| 11 | MGCS36044_00788 | ***sufS*** |  | cysteine desulfurase SufS | E | METABOLISM; amino acid transport and metabolism | | | | core | Pro |
| 12 | MGCS36044_00824 | ***yqeG*** |  | HAD IIIA-type phosphatase YqeG | F | METABOLISM; nucleotide transport and metabolism | | | | core | Pro |
| 13 | MGCS36044_00830 | ***nadD*** |  | nicotinate-nucleotide adenylyltransferase NadD | H | METABOLISM; coenzyme transport and metabolism | | | | core | Pro |
| 14 | MGCS36044_00876 | ***yceD*** |  | large ribosomal RNA subunit accumulation protein YceD | M | CELLULAR PROCESSES AND SIGNALING; cell wall/membrane/envelope biogenesis | | | | core | Pro |
| 15 | MGCS36044_00878 | ***covR*** | Virulence | TCS DNA-binding response regulator CovR | T | CELLULAR PROCESSES AND SIGNALING; signal transduction mechanisms | | | | core | Pro |
| 16 | MGCS36044_00900 | ***greA*** |  | transcription elongation factor GreA | K | INFORMATION STORAGE AND PROCESSING; transcription | | | | core | Pro |
| 17 | MGCS36044_00938 | ***-*** |  | ECF transporter S component | I | METABOLISM; lipid metabolism | | | | core | Pro |
| 18 | MGCS36044_00968 | ***clpP*** |  | ATP-dependent Clp protease proteolytic subunit ClpP | O | CELLULAR PROCESSES AND SIGNALING; post-translational modification, protein turnover, and chaperones | | | | core | Pro |
| 19 | MGCS36044_01060 | ***mtnN*** |  | 5'-methylthioadenosine/adenosylhomocysteine nucleosidase MtnN | F | METABOLISM; nucleotide transport and metabolism | | | | core | Pro |
| 20 | MGCS36044_01070 | ***mtsC*** |  | metal ABC transporter permease MtsC | P | METABOLISM; inorganic ion transport and metabolism | | | | core | Pro |
| 21 | MGCS36044_01080 | ***rplA*** |  | 50S ribosomal L1 protein RplA | J | INFORMATION STORAGE AND PROCESSING; translation, ribosomal structure and biogenesis | | | | core | Pro |
| 22 | MGCS36044_01130 | ***-*** |  | IS1548 family transposase | L | INFORMATION STORAGE AND PROCESSING; DNA replication, recombination and repair | | | | core | Pro |
| 23 | MGCS36044_01240 | ***vicK*** | Virulence | TCS signal transduction sensor kinase VicK | T | CELLULAR PROCESSES AND SIGNALING; signal transduction mechanisms | | | | core | Pro |
| 24 | MGCS36044_01244 | ***rnc*** |  | ribonuclease III Rnc | A | RNA processing and modification | | | | core | Pro |
| 25 | MGCS36044_01250 | ***-*** |  | IS1548 family transposase | L | INFORMATION STORAGE AND PROCESSING; DNA replication, recombination and repair | | | | core | Pro |
| 26 | MGCS36044_01330 | ***thiT*** |  | energy-coupled thiamine transporter ThiT | H | METABOLISM; coenzyme transport and metabolism | | | | core | Pro |
| 27 | MGCS36044_01336 | ***-*** |  | transposase IS116/IS110/IS902 family protein | core | | Pro |  |  |  |  |
| 28 | MGCS36044_01348 | ***ftsW*** |  | cell division protein FtsW | D | CELLULAR PROCESSES AND SIGNALING; cell cycle control, cell division, chromosome partitioning | | | | core | Pro |
| 29 | MGCS36044_01436 | ***aspC*** |  | aspartate aminotransferase protein AspC | E | METABOLISM; amino acid transport and metabolism | | | | core | Pro |
| 30 | MGCS36044_01438 | ***asnC*** |  | asparaginyl-tRNA synthetase protein AsnC | J | INFORMATION STORAGE AND PROCESSING; translation, ribosomal structure and biogenesis | | | | core | Pro |
| 31 | MGCS36044_01470 | ***rpmE*** |  | 50S ribosomal L31 type B protein RpmE | J | INFORMATION STORAGE AND PROCESSING; translation, ribosomal structure and biogenesis | | | | core | Pro |
| 32 | MGCS36044_01564 | ***atpC*** |  | ATP synthase epsilon subunit AtpC | C | METABOLISM; energy production and conversion | | | | core | Pro |
| 33 | MGCS36044_01588 | ***rexB*** |  | ATP-dependent nuclease B subunit RexB | L | INFORMATION STORAGE AND PROCESSING; DNA replication, recombination and repair | | | | core | Pro |
| 34 | MGCS36044_01596 | ***-*** |  | IS1548 family transposase | L | INFORMATION STORAGE AND PROCESSING; DNA replication, recombination and repair | | | | core | Pro |
| 35 | MGCS36044_01634 | ***-*** |  | RfbX superfamily lipopolysaccharide biosynthesis protein | M | CELLULAR PROCESSES AND SIGNALING; cell wall/membrane/envelope biogenesis | | | | core | Pro |
| 36 | MGCS36044_01654 | ***rlmK*** |  | 23S rRNA methyltransferase RmlK | J | INFORMATION STORAGE AND PROCESSING; translation, ribosomal structure and biogenesis | | | | core | Pro |
| 37 | MGCS36044_01656 | ***aroD*** |  | type I 3-dehydroquinate dehydratase AroD | E | METABOLISM; amino acid transport and metabolism | | | | core | Pro |
| 38 | MGCS36044_01802 | ***rplJ*** |  | 50S ribosomal L10 protein RplJ | J | INFORMATION STORAGE AND PROCESSING; translation, ribosomal structure and biogenesis | | | | core | Pro |
| 39 | MGCS36044_01806 | ***-*** |  | rli38 |  |  | | | | core | RNA |
| 40 | MGCS36044_01982 | ***-*** |  | Na+ driven multidrug efflux pump | P | METABOLISM; inorganic ion transport and metabolism | | | | core | Pro |
| 41 | MGCS36044_02078 | ***-*** |  | disrupted IS3 family transposase encoding gene | core | | Pro |  |  |  |  |
| 42 | MGCS36044_02080 | ***-*** |  | IS30 family transposase |  |  | | | | core | Pro |
| 43 | MGCS36044_02242 | ***-*** |  | disrupted IS3 family transposase encoding gene | core | | Pro |  |  |  |  |
| 44 | MGCS36044_02298 | ***ptsB1*** |  | phosphate ABC transporter ATP-binding protein PstB1 | P | METABOLISM; Inorganic ion transport and metabolism | | | | core | Pro |
| 45 | MGCS36044_02300 | ***ptsB2*** |  | phosphate ABC transporter ATP-binding protein PstB2 | P | METABOLISM; Inorganic ion transport and metabolism | | | | core | Pro |
| 46 | MGCS36044_02348 | ***-*** |  | glycine |  |  | | | | core | RNA |
| 47 | MGCS36044_02388 | ***-*** |  | IS1182 family transposase |  | | | | core | | Pro |
| 48 | MGCS36044_02444 | ***rpiA*** |  | ribose-5-phosphate isomerase RpiA | G | METABOLISM; carbohydrate transport and metabolism | | | | core | Pro |
| 49 | MGCS36044_02448 | ***pepV*** |  | dipeptidase PepV | E | METABOLISM; amino acid transport and metabolism | | | | core | Pro |
| 50 | MGCS36044_02608 | ***trmK*** |  | tRNA (adenine(22)-N(1))-methyltransferase TrmK | L | INFORMATION STORAGE AND PROCESSING; DNA replication, recombination and repair | | | | core | Pro |
| 51 | MGCS36044_02612 | ***dnaD*** |  | DNA replication protein DnaD | L | INFORMATION STORAGE AND PROCESSING; DNA replication, recombination and repair | | | | core | Pro |
| 52 | MGCS36044_02668 | ***dltD*** |  | D-alanyl-lipoteichoic acid biosynthesis protein DltD | M | CELLULAR PROCESSES AND SIGNALING; cell wall/membrane/envelope biogenesis | | | | core | Pro |
| 53 | MGCS36044_02730 | ***-*** |  | disrupted IS30 family transposase encoding gene | core | | Pro |  |  |  |  |
| 54 | MGCS36044_02886 | ***ptsI*** |  | phosphoenolpyruvate--protein phosphotransferase PtsI | G | METABOLISM; carbohydrate transport and metabolism | | | | core | Pro |
| 55 | MGCS36044_02938 | ***holA*** |  | DNA polymerase III delta subunit HolA | L | INFORMATION STORAGE AND PROCESSING; DNA replication, recombination and repair | | | | core | Pro |
| 56 | MGCS36044_02952 | ***deaD*** |  | DEAD/DEAH box helicase | L | INFORMATION STORAGE AND PROCESSING; DNA replication, recombination and repair | | | | core | Pro |
| 57 | MGCS36044_03018 | ***folD*** |  | bifunctional methylenetetrahydrofolate dehydrogenase/methenyltetrahydrofolate cyclohydrolase FolD | E | METABOLISM; amino acid transport and metabolism | | | | core | Pro |
| 58 | MGCS36044_03064 | ***murD*** |  | UDP-N-acetylmuramoyl-L-alanine--D-glutamate ligase MurD | core | | Pro |  |  |  |  |
| 59 | MGCS36044_03112 | ***coaD*** |  | pantetheine-phosphate adenylyltransferase CoaD | H | METABOLISM; coenzyme metabolism | | | | core | Pro |
| 60 | MGCS36044_03288 | ***cysK*** |  | cysteine synthase A CysK | E | METABOLISM; amino acid transport and metabolism | | | | core | Pro |
| 61 | MGCS36044_03294 | ***liaR*** | Virulence | three component system signal transduction response regulator protein LiaR | K | INFORMATION STORAGE AND PROCESSING; transcription | | | | core | Pro |
| 62 | MGCS36044_03302 | ***pppL*** |  | Stp1/IreP family PP2C-type Ser/Thr phosphatase | T | CELLULAR PROCESSES AND SIGNALING; signal transduction mechanisms | | | | core | Pro |
| 63 | MGCS36044_03472 | ***rimP*** |  | ribosome maturation factor RimP | J | INFORMATION STORAGE AND PROCESSING; translation, ribosomal structure and biogenesis | | | | core | Pro |
| 64 | MGCS36044_03522 | ***accB*** |  | acetyl-CoA carboxylase biotin carboxyl carrier protein AccB | I | METABOLISM; lipid metabolism | | | | core | Pro |
| 65 | MGCS36044_03618 | ***alr*** |  | alanine racemase Alr | E | METABOLISM; amino acid transport and metabolism | | | | core | Pro |
| 66 | MGCS36044_03636 | ***nusB*** |  | transcription termination protein NusB | K | INFORMATION STORAGE AND PROCESSING; transcription | | | | core | Pro |
| 67 | MGCS36044_03638 | ***-*** |  | Asp23/Gls24 family envelope stress response protein | R | POORLY CHARACTERIZED; general function prediction only | | | | core | Pro |
| 68 | MGCS36044_03738 | ***-*** |  | disrupted IS3 family transposase encoding gene | RD.8 | | Pro |  |  |  |  |
| 69 | MGCS36044_03824 | ***thiN*** |  | thiamine diphosphokinase ThiN | H | METABOLISM; coenzyme metabolism | | | | core | Pro |
| 70 | MGCS36044_03826 | ***rpe*** |  | ribulose-phosphate 3-epimerase Rpe | G | METABOLISM; carbohydrate transport and metabolism | | | | core | Pro |
| 71 | MGCS36044_03978 | ***-*** |  | IS30 family transposase |  |  | | | | core | Pro |
| 72 | MGCS36044_03980 | ***-*** |  | disrupted IS3 family transposase encoding gene | core | | Pro |  |  |  |  |
| 73 | MGCS36044_03990 | ***-*** |  | IS982 family transposase |  |  | | | | core | Pro |
| 74 | MGCS36044_04000 | ***nusG*** |  | transcription antitermination protein NusG | K | INFORMATION STORAGE AND PROCESSING; transcription | | | | core | Pro |
| 75 | MGCS36044_04108 | ***recA*** |  | recombinase RecA | L | INFORMATION STORAGE AND PROCESSING; DNA replication, recombination and repair | | | | core | Pro |
| 76 | MGCS36044_04202 | ***rpsD*** |  | 30S ribosomal S4 protein RpsD | J | INFORMATION STORAGE AND PROCESSING; translation, ribosomal structure and biogenesis | | | | core | Pro |
| 77 | MGCS36044_04220 | ***-*** |  | Spd-sr37 |  |  | | | | core | RNA |

† Genes are ordered ascendingly based on the locus tag numbers.

‡ Putative virulence genes (Eraso et al., 2024, *mBio*).

††Genomic location. Core: the core genome, Acc: the accessory genome, ROD: regions of difference.

‡‡ Pro: protein.
