## Supplementary material for "Gene contribution of *Streptococcus dysgalactiae* subspecies *equisimilis*, an emerging pathogen, to experimental primate necrotizing myositis": Table S4A

**Table S4A. Essential genes shared by MGCS36044 and MGCS36089**

| **No.^†^** | **Locus tag 044^∫^** | **Locus tag 089^∫∫^** | **Gene** | **Virulence^‡^** | **Exported^¶^** | **Function** | **COG class** | **COG description** | **Core/ Acc^††^** | **Pro/ RNA^‡‡^** |
| --- | --- | --- | --- | --- | --- | --- | --- | --- | --- | --- |
| 1 | MGCS36044_00002 | MGCS36089_00002 | ***dnaA*** |  |  | chromosomal replication initiator protein DnaA | L | INFORMATION STORAGE AND PROCESSING; replication, recombination and repair | Core | Pro |
| 2 | MGCS36044_00004 | MGCS36089_00004 | ***dnaN*** |  |  | DNA polymerase III subunit beta protein DnaN | L | INFORMATION STORAGE AND PROCESSING; replication, recombination and repair | Core | Pro |
| 3 | MGCS36044_00010 | MGCS36089_00010 | ***engD*** |  |  | redox-regulated ATPase EngD | J | INFORMATION STORAGE AND PROCESSING; translation, ribosomal structure and biogenesis | Core | Pro |
| 4 | MGCS36044_00012 | MGCS36089_00012 | ***pth*** |  |  | aminoacyl-tRNA hydrolase Pth | J | INFORMATION STORAGE AND PROCESSING; translation, ribosomal structure and biogenesis | Core | Pro |
| 5 | MGCS36044_00018 | MGCS36089_00018 | ***-*** |  |  | RNA-binding S4 domain-containing protein | J | INFORMATION STORAGE AND PROCESSING; translation, ribosomal structure and biogenesis | Core | Pro |
| 6 | MGCS36044_00020 | MGCS36089_00020 | ***divIC*** |  |  | septum formation initiator family protein FtsB/DivIC | D | CELLULAR PROCESSES AND SIGNALING; cell division and chromosome partitioning | Core | Pro |
| 7 | MGCS36044_00024 | MGCS36089_00024 | ***-*** |  | Secreted | class A beta-lactamase-related serine hydrolase | M | CELLULAR PROCESSES AND SIGNALING; cell envelope biogenesis, outermembrane | Core | Pro |
| 8 | MGCS36044_00026 | MGCS36089_00026 | ***tilS*** |  |  | tRNA lysidine(34) synthetase TilS | J | INFORMATION STORAGE AND PROCESSING; translation, ribosomal structure and biogenesis | Core | Pro |
| 9 | MGCS36044_00030 | MGCS36089_00030 | ***ftsH*** |  |  | ATP-dependent zinc metalloprotease FtsH | O | CELLULAR PROCESSES AND SIGNALING; post-translational modification, protein turnover, and chaperones | Core | Pro |
| 10 | MGCS36044_00104 | MGCS36089_00104 | ***sibA*** |  | Secreted | CHAP domain-containing protein/secreted immunoglobulin-binding protein SibA | U | CELLULAR PROCESSES AND SIGNALING; intracellular trafficking, secretion, and vesicular transport | Core | Pro |
| 11 | MGCS36044_00106 | MGCS36089_00106 | ***prs*** |  |  | ribose-phosphate pyrophosphokinase PrsA | G | METABOLISM; carbohydrate transport and metabolism | Core | Pro |
| 12 | MGCS36044_00110 | MGCS36089_00110 | ***plsX*** |  |  | phosphate acyltransferase PlsX | I | METABOLISM; lipid metabolism | Core | Pro |
| 13 | MGCS36044_00152 | MGCS36089_00152 | ***oatA*** |  |  | acetyltransferase OatA | M | CELLULAR PROCESSES AND SIGNALING; cell envelope biogenesis, outermembrane | Core | Pro |
| 14 | MGCS36044_00168 | MGCS36089_00168 | ***rplC*** |  |  | 50S ribosomal L3 protein RplC | J | INFORMATION STORAGE AND PROCESSING; translation, ribosomal structure and biogenesis | Core | Pro |
| 15 | MGCS36044_00170 | MGCS36089_00170 | ***rplD*** |  |  | 50S ribosomal L4 protein RplD | J | INFORMATION STORAGE AND PROCESSING; translation, ribosomal structure and biogenesis | Core | Pro |
| 16 | MGCS36044_00172 | MGCS36089_00172 | ***rplW*** |  |  | 50S ribosomal L23 protein RplW | J | INFORMATION STORAGE AND PROCESSING; translation, ribosomal structure and biogenesis | Core | Pro |
| 17 | MGCS36044_00174 | MGCS36089_00174 | ***rplB*** |  |  | 50S ribosomal L2 protein RplB | J | INFORMATION STORAGE AND PROCESSING; translation, ribosomal structure and biogenesis | Core | Pro |
| 18 | MGCS36044_00176 | MGCS36089_00176 | ***rpsS*** |  |  | 30S ribosomal S19 protein RpsS | J | INFORMATION STORAGE AND PROCESSING; translation, ribosomal structure and biogenesis | Core | Pro |
| 19 | MGCS36044_00178 | MGCS36089_00178 | ***rplV*** |  |  | 50S ribosomal L22 protein RplV | J | INFORMATION STORAGE AND PROCESSING; translation, ribosomal structure and biogenesis | Core | Pro |
| 20 | MGCS36044_00180 | MGCS36089_00180 | ***rpsC*** |  |  | 30S ribosomal S3 protein RpsC | J | INFORMATION STORAGE AND PROCESSING; translation, ribosomal structure and biogenesis | Core | Pro |
| 21 | MGCS36044_00184 | MGCS36089_00184 | ***rpmC*** |  |  | 50S ribosomal L16 protein RpmC | J | INFORMATION STORAGE AND PROCESSING; translation, ribosomal structure and biogenesis | Core | Pro |
| 22 | MGCS36044_00186 | MGCS36089_00186 | ***rpsQ*** |  |  | 30S ribosomal S17 protein RpsQ | J | INFORMATION STORAGE AND PROCESSING; translation, ribosomal structure and biogenesis | Core | Pro |
| 23 | MGCS36044_00188 | MGCS36089_00188 | ***rplN*** |  |  | 50S ribosomal L14 protein RplN | J | INFORMATION STORAGE AND PROCESSING; translation, ribosomal structure and biogenesis | Core | Pro |
| 24 | MGCS36044_00190 | MGCS36089_00190 | ***rplX*** |  |  | 50S ribosomal L24 protein RplX | J | INFORMATION STORAGE AND PROCESSING; translation, ribosomal structure and biogenesis | Core | Pro |
| 25 | MGCS36044_00192 | MGCS36089_00192 | ***rplE*** |  |  | 50S ribosomal L5 protein RplE | J | INFORMATION STORAGE AND PROCESSING; translation, ribosomal structure and biogenesis | Core | Pro |
| 26 | MGCS36044_00196 | MGCS36089_00196 | ***rpsH*** |  |  | 30S ribosomal S8 protein RpsH | J | INFORMATION STORAGE AND PROCESSING; translation, ribosomal structure and biogenesis | Core | Pro |
| 27 | MGCS36044_00198 | MGCS36089_00198 | ***rplF*** |  |  | 50S ribosomal L6 protein RplF | J | INFORMATION STORAGE AND PROCESSING; translation, ribosomal structure and biogenesis | Core | Pro |
| 28 | MGCS36044_00200 | MGCS36089_00200 | ***rplR*** |  |  | 50S ribosomal L18 protein RplR | J | INFORMATION STORAGE AND PROCESSING; translation, ribosomal structure and biogenesis | Core | Pro |
| 29 | MGCS36044_00204 | MGCS36089_00204 | ***rpmD*** |  |  | 50S ribosomal L30 protein RpmD | J | INFORMATION STORAGE AND PROCESSING; translation, ribosomal structure and biogenesis | Core | Pro |
| 30 | MGCS36044_00208 | MGCS36089_00208 | ***secY*** |  |  | preprotein translocase subunit SecY | U | CELLULAR PROCESSES AND SIGNALING; intracellular trafficking, secretion, and vesicular transport | Core | Pro |
| 31 | MGCS36044_00212 | MGCS36089_00212 | ***infA*** |  |  | translation initiation factor IF-1 protein InfA | J | INFORMATION STORAGE AND PROCESSING; translation, ribosomal structure and biogenesis | Core | Pro |
| 32 | MGCS36044_00216 | MGCS36089_00216 | ***rpsM*** |  |  | 30S ribosomal S13 protein RpsM | J | INFORMATION STORAGE AND PROCESSING; translation, ribosomal structure and biogenesis | Core | Pro |
| 33 | MGCS36044_00218 | MGCS36089_00218 | ***rpsK*** |  |  | 30S ribosomal S11 protein RpsK | J | INFORMATION STORAGE AND PROCESSING; translation, ribosomal structure and biogenesis | Core | Pro |
| 34 | MGCS36044_00220 | MGCS36089_00220 | ***rpoA*** |  |  | DNA-directed RNA polymerase subunit alpha RpoA | K | INFORMATION STORAGE AND PROCESSING; transcription | Core | Pro |
| 35 | MGCS36044_00222 | MGCS36089_00222 | ***rplQ*** |  |  | 50S ribosomal L17 protein RplQ | J | INFORMATION STORAGE AND PROCESSING; translation, ribosomal structure and biogenesis | Core | Pro |
| 36 | MGCS36044_00226 | MGCS36089_00226 | ***-*** |  |  | disrupted IS1239 transposase encoding gene |  |  | Core | Pro |
| 37 | MGCS36044_00228 | MGCS36089_00228 | ***-*** |  |  | IS30 family transposase |  |  | Core | Pro |
| 38 | MGCS36044_00308 | MGCS36089_00308 | ***tyrS*** |  |  | tyrosyl-tRNA synthetase TyrS | J | INFORMATION STORAGE AND PROCESSING; translation, ribosomal structure and biogenesis | Core | Pro |
| 39 | MGCS36044_00310 | MGCS36089_00310 | ***pbp1B*** |  |  | bifunctional PG transglycosylase-transpeptidase, penicillin-binding protein PBP1B | M | CELLULAR PROCESSES AND SIGNALING; cell envelope biogenesis, outermembrane | Core | Pro |
| 40 | MGCS36044_00312 | MGCS36089_00312 | ***-*** |  |  | Lacto-rpoB |  |  | Core | RNA |
| 41 | MGCS36044_00314 | MGCS36089_00314 | ***rpoB*** |  |  | DNA-directed RNA polymerase subunit beta RpoB | K | INFORMATION STORAGE AND PROCESSING; transcription | Core | Pro |
| 42 | MGCS36044_00316 | MGCS36089_00316 | ***rpoC*** |  |  | DNA-directed RNA polymerase subunit beta' RpoC | K | INFORMATION STORAGE AND PROCESSING; transcription | Core | Pro |
| 43 | MGCS36044_00336 | MGCS36089_00336 | ***ackA*** |  |  | acetate kinase AckA | C | METABOLISM; energy production and conversion | Core | Pro |
| 44 | MGCS36044_00542 | MGCS36089_00540 | ***uppS*** |  |  | UDP pyrophosphate synthase UppS | I | METABOLISM; lipid metabolism | Core | Pro |
| 45 | MGCS36044_00548 | MGCS36089_00546 | ***proS*** |  |  | prolyl-tRNA synthetase ProS | J | INFORMATION STORAGE AND PROCESSING; translation, ribosomal structure and biogenesis | Core | Pro |
| 46 | MGCS36044_00552 | MGCS36089_00550 | ***polC*** |  |  | DNA polymerase III PolC | L | INFORMATION STORAGE AND PROCESSING; replication, recombination and repair | Core | Pro |
| 47 | MGCS36044_00558 | MGCS36089_00556 | ***def*** |  |  | peptide deformylase Def | J | INFORMATION STORAGE AND PROCESSING; translation, ribosomal structure and biogenesis | Core | Pro |
| 48 | MGCS36044_00564 | MGCS36089_00562 | ***rpsO*** |  |  | 30S ribosomal S15 protein RpsO | J | INFORMATION STORAGE AND PROCESSING; translation, ribosomal structure and biogenesis | Core | Pro |
| 49 | MGCS36044_00574 | MGCS36089_00572 | ***pnp*** |  |  | polyribonucleotide nucleotidyltransferase Pnp | J | INFORMATION STORAGE AND PROCESSING; translation, ribosomal structure and biogenesis | Core | Pro |
| 50 | MGCS36044_00576 | MGCS36089_00574 | ***-*** |  |  | polynucleotide phosphorylase/polyadenylase | L | INFORMATION STORAGE AND PROCESSING; replication, recombination and repair | Core | Pro |
| 51 | MGCS36044_00578 | MGCS36089_00576 | ***cysE*** |  |  | serine O-acetyltransferase CysE | E | METABOLISM; amino acid transport and metabolism | Core | Pro |
| 52 | MGCS36044_00582 | MGCS36089_00580 | ***cysS*** |  |  | cysteine--tRNA synthetase CysS | J | INFORMATION STORAGE AND PROCESSING; translation, ribosomal structure and biogenesis | Core | Pro |
| 53 | MGCS36044_00600 | MGCS36089_00598 | ***-*** |  |  | IS30 family transposase |  |  | Core | Pro |
| 54 | MGCS36044_00602 | MGCS36089_00600 | ***-*** |  |  | L13_leader |  |  | Core | RNA |
| 55 | MGCS36044_00604 | MGCS36089_00602 | ***rplM*** |  |  | 50S ribosomal L13 protein RplM | J | INFORMATION STORAGE AND PROCESSING; translation, ribosomal structure and biogenesis | Core | Pro |
| 56 | MGCS36044_00606 | MGCS36089_00604 | ***rpsI*** |  |  | 30S ribosomal S9 protein RpsI | J | INFORMATION STORAGE AND PROCESSING; translation, ribosomal structure and biogenesis | Core | Pro |
| 57 | MGCS36044_00610 | MGCS36089_00608 | ***-*** |  |  | helix-turn-helix transcriptional regulator | K | INFORMATION STORAGE AND PROCESSING; transcription | RD.3 | Pro |
| 58 | MGCS36044_00622 | MGCS36089_00620 | ***-*** |  |  | DNA cytosine methyltransferase | L | INFORMATION STORAGE AND PROCESSING; replication, recombination and repair | RD.3 | Pro |
| 59 | MGCS36044_00668 | MGCS36089_00666 | ***-*** |  |  | Cro/CI family transcriptional regulator | K | INFORMATION STORAGE AND PROCESSING; transcription | RD.3 | Pro |
| 60 | MGCS36044_00708 | MGCS36089_00706 | ***-*** |  |  | IS30 family transposase |  |  | Core | Pro |
| 61 | MGCS36044_00722 | MGCS36089_00720 | ***comX_1*** |  |  | competence protein ComX | K | INFORMATION STORAGE AND PROCESSING; transcription | Core | Pro |
| 62 | MGCS36044_00742 | MGCS36089_00738 | ***rpoE*** |  |  | DNA-directed RNA polymerase subunit delta RpoE | K | INFORMATION STORAGE AND PROCESSING; transcription | Core | Pro |
| 63 | MGCS36044_00758 | MGCS36089_00754 | ***fba_2*** |  |  | fructose-bisphosphate aldolase | G | METABOLISM; carbohydrate transport and metabolism | Core | Pro |
| 64 | MGCS36044_00764 | MGCS36089_00760 | ***-*** |  |  | Asp23/Gls24 family envelope stress response protein | M | CELLULAR PROCESSES AND SIGNALING; cell envelope biogenesis, outermembrane | Core | Pro |
| 65 | MGCS36044_00766 | MGCS36089_00762 | ***-*** |  |  | DAK2 domain-containing protein | S | POORLY CHARACTERIZED; function unknown | Core | Pro |
| 66 | MGCS36044_00780 | MGCS36089_00776 | ***mecA*** |  |  | negative regulator of genetic competence, adaptor protein MecA | O | CELLULAR PROCESSES AND SIGNALING; post-translational modification, protein turnover, and chaperones | Core | Pro |
| 67 | MGCS36044_00782 | MGCS36089_00778 | ***rgpG*** |  |  | undecaprenyl/decaprenyl-phosphate alpha-N-acetylglucosaminyl 1-phosphate transferase RgpG | M | CELLULAR PROCESSES AND SIGNALING; cell envelope biogenesis, outermembrane | Core | Pro |
| 68 | MGCS36044_00822 | MGCS36089_00808 | ***comX_2*** |  |  | competence protein ComX | K | INFORMATION STORAGE AND PROCESSING; transcription | Core | Pro |
| 69 | MGCS36044_00826 | MGCS36089_00812 | ***yqeH*** |  |  | ribosome biogenesis GTPase YqeH | P | METABOLISM; inorganic ion transport and metabolism | Core | Pro |
| 70 | MGCS36044_00864 | MGCS36089_00850 | ***sstT*** |  |  | serine/threonine transporter SstT | E | METABOLISM; amino acid transport and metabolism | Core | Pro |
| 71 | MGCS36044_00884 | MGCS36089_00870 | ***dnaB*** |  |  | replication initiation and membrane attachment protein DnaB | L | INFORMATION STORAGE AND PROCESSING; replication, recombination and repair | Core | Pro |
| 72 | MGCS36044_00886 | MGCS36089_00872 | ***dnaI*** |  |  | primosomal protein DnaI | L | INFORMATION STORAGE AND PROCESSING; replication, recombination and repair | Core | Pro |
| 73 | MGCS36044_00888 | MGCS36089_00874 | ***der*** |  |  | ribosome biogenesis GTPase Der | J | INFORMATION STORAGE AND PROCESSING; translation, ribosomal structure and biogenesis | Core | Pro |
| 74 | MGCS36044_00894 | MGCS36089_00880 | ***murC*** |  |  | UDP-N-acetylmuramate--L-alanine ligase MurC | M | CELLULAR PROCESSES AND SIGNALING; cell envelope biogenesis, outermembrane | Core | Pro |
| 75 | MGCS36044_00898 | MGCS36089_00884 | ***mltG*** |  |  | endolytic transglycosylase MltG | F | METABOLISM; nucleotide transport and metabolism | Core | Pro |
| 76 | MGCS36044_00902 | MGCS36089_00888 | ***yidC_1*** |  | Lipo | membrane protein insertase lipoprotein YidC | M | CELLULAR PROCESSES AND SIGNALING; cell envelope biogenesis, outermembrane | Core | Pro |
| 77 | MGCS36044_00912 | MGCS36089_00898 | ***yneF*** |  |  | YneF family protein | S | POORLY CHARACTERIZED; function unknown | Core | Pro |
| 78 | MGCS36044_00914 | MGCS36089_00900 | ***murI*** |  |  | glutamate racemase MurI | M | CELLULAR PROCESSES AND SIGNALING; cell envelope biogenesis, outermembrane | Core | Pro |
| 79 | MGCS36044_00950 | MGCS36089_00936 | ***ppaC*** |  |  | manganese-dependent inorganic pyrophosphatase PpaC | R | General function prediction | Core | Pro |
| 80 | MGCS36044_00962 | MGCS36089_00948 | ***murE_1*** |  |  | UDP-N-acetylmuramoyl-L-alanyl-D-glutamate--L- lysine ligase MurE | M | CELLULAR PROCESSES AND SIGNALING; cell envelope biogenesis, outermembrane | Core | Pro |
| 81 | MGCS36044_00964 | MGCS36089_00950 | ***murJ*** |  |  | peptidoglycan lipid-II intermdiate flippase MurJ | O | CELLULAR PROCESSES AND SIGNALING; post-translational modification, protein turnover, and chaperones | Core | Pro |
| 82 | MGCS36044_00988 | MGCS36089_00974 | ***tmk*** |  |  | thymidylate kinase Tmk | F | METABOLISM; Nucleotide transport and metabolism | Core | Pro |
| 83 | MGCS36044_00990 | MGCS36089_00976 | ***holB*** |  |  | DNA polymerase III subunit delta' HolB | L | INFORMATION STORAGE AND PROCESSING; replication, recombination and repair | Core | Pro |
| 84 | MGCS36044_00994 | MGCS36089_00980 | ***yabA*** |  |  | DNA replication initiation control protein YabA | L | INFORMATION STORAGE AND PROCESSING; replication, recombination and repair | Core | Pro |
| 85 | MGCS36044_01028 | MGCS36089_01014 | ***metS*** |  |  | methionine--tRNA synthase MetS | J | INFORMATION STORAGE AND PROCESSING; translation, ribosomal structure and biogenesis | Core | Pro |
| 86 | MGCS36044_01054 | MGCS36089_01040 | ***glmU*** |  |  | bifunctional UDP-N-acetylglucosamine diphosphorylase/glucosamine-1-phosphate N-acetyltransferase GlmU | G | METABOLISM; carbohydrate transport and metabolism | Core | Pro |
| 87 | MGCS36044_01078 | MGCS36089_01064 | ***rplK*** |  |  | 50S ribosomal L11P protein RplK | J | INFORMATION STORAGE AND PROCESSING; translation, ribosomal structure and biogenesis | Core | Pro |
| 88 | MGCS36044_01084 | MGCS36089_01070 | ***pyrH*** |  |  | UMP kinase PyrH | F | METABOLISM; nucleotide transport and metabolism | Core | Pro |
| 89 | MGCS36044_01086 | MGCS36089_01072 | ***frr*** |  |  | ribosome recycling factor Frr | J | INFORMATION STORAGE AND PROCESSING; translation, ribosomal structure and biogenesis | Core | Pro |
| 90 | MGCS36044_01102 | MGCS36089_01088 | ***ybeY*** |  |  | rRNA maturation RNase YbeY | J | INFORMATION STORAGE AND PROCESSING; translation, ribosomal structure and biogenesis | Core | Pro |
| 91 | MGCS36044_01104 | MGCS36089_01090 | ***dgkA*** |  |  | diacylglycerol kinase DgkA | M | CELLULAR PROCESSES AND SIGNALING; cell envelope biogenesis, outermembrane | Core | Pro |
| 92 | MGCS36044_01106 | MGCS36089_01092 | ***era*** |  |  | GTPase Era | J | INFORMATION STORAGE AND PROCESSING; translation, ribosomal structure and biogenesis | Core | Pro |
| 93 | MGCS36044_01122 | MGCS36089_01108 | ***-*** |  |  | IS1182 family transposase |  |  | Core | Pro |
| 94 | MGCS36044_01124 | MGCS36089_01110 | ***-*** |  |  | disrupted IS3 family transposase encoding gene |  |  | Core | Pro |
| 95 | MGCS36044_01164 | MGCS36089_01150 | ***coaE*** |  |  | dephospho-CoA kinase CoaE | H | METABOLISM; coenzyme transport and metabolism | Core | Pro |
| 96 | MGCS36044_01178 | MGCS36089_01164 | ***smpB*** |  |  | SsrA(tmRNA)-binding protein SmpB | O | CELLULAR PROCESSES AND SIGNALING; post-translational modification, protein turnover, and chaperones | Core | Pro |
| 97 | MGCS36044_01182 | MGCS36089_01168 | ***-*** |  |  | disrupted IS3 family transposase encoding gene |  |  | Core | Pro |
| 98 | MGCS36044_01194 | MGCS36089_01180 | ***ccpA*** | Virulence |  | catabolite control protein CcpA | K | INFORMATION STORAGE AND PROCESSING; transcription | Core | Pro |
| 99 | MGCS36044_01198 | MGCS36089_01184 | ***-*** |  |  | glycosyltransferase | R | General function prediction only | Core | Pro |
| 100 | MGCS36044_01200 | MGCS36089_01186 | ***-*** |  |  | glycosyltransferase | M | CELLULAR PROCESSES AND SIGNALING; cell envelope biogenesis, outermembrane | Core | Pro |
| 101 | MGCS36044_01202 | MGCS36089_01188 | ***thrS*** |  |  | threonyl-tRNA synthetase ThrS | J | INFORMATION STORAGE AND PROCESSING; translation, ribosomal structure and biogenesis | Core | Pro |
| 102 | MGCS36044_01238 | MGCS36089_01224 | ***vicR*** | Virulence |  | TCS DNA-binding response regulator VicR | T | CELLULAR PROCESSES AND SIGNALING; signal transduction mechanisms | Core | Pro |
| 103 | MGCS36044_01246 | MGCS36089_01232 | ***smc*** |  |  | chromosome segregation protein Smc | D | CELLULAR PROCESSES AND SIGNALING; cell division and chromosome partitioning | Core | Pro |
| 104 | MGCS36044_01256 | MGCS36089_01244 | ***ftsY*** |  |  | signal recognition particle-docking protein FtsY | U | CELLULAR PROCESSES AND SIGNALING; intracellular trafficking, secretion, and vesicular transport | Core | Pro |
| 105 | MGCS36044_01300 | MGCS36089_01288 | ***hprK*** |  |  | HPr(Ser) kinase/phosphatase HprK | T | CELLULAR PROCESSES AND SIGNALING; signal transduction mechanisms | Core | Pro |
| 106 | MGCS36044_01308 | MGCS36089_01296 | ***-*** |  |  | DUF3270 domain-containing protein | S | POORLY CHARACTERIZED; function unknown | Core | Pro |
| 107 | MGCS36044_01320 | MGCS36089_01308 | ***lysS*** |  |  | lysyl-tRNA synthetase LysS | J | INFORMATION STORAGE AND PROCESSING; translation, ribosomal structure and biogenesis | Core | Pro |
| 108 | MGCS36044_01354 | MGCS36089_01342 | ***tufA*** |  |  | translation elongation factor Tu protein TufA | J | INFORMATION STORAGE AND PROCESSING; translation, ribosomal structure and biogenesis | Core | Pro |
| 109 | MGCS36044_01364 | MGCS36089_01352 | ***tpiA*** |  |  | triose-phosphate isomerase TpiA | G | METABOLISM; carbohydrate transport and metabolism | Core | Pro |
| 110 | MGCS36044_01366 | MGCS36089_01354 | ***-*** |  |  | disrupted IS3 family transposase encoding gene |  |  | Core | Pro |
| 111 | MGCS36044_01368 | MGCS36089_01356 | ***-*** |  |  | disrupted IS3 family transposase encoding gene |  |  | Core | Pro |
| 112 | MGCS36044_01372 | MGCS36089_01360 | ***murN*** |  |  | peptidoglycan lipid II-Ala--L-alanine ligase protein MurN | M | CELLULAR PROCESSES AND SIGNALING; cell envelope biogenesis, outermembrane | Core | Pro |
| 113 | MGCS36044_01374 | MGCS36089_01362 | ***murM*** |  |  | peptidoglycan lipid II--L-alanine ligase protein MurM | M | CELLULAR PROCESSES AND SIGNALING; cell envelope biogenesis, outermembrane | Core | Pro |
| 114 | MGCS36044_01376 | MGCS36089_01364 | ***-*** |  |  | sugar-phosphatase | R | General function prediction only | Core | Pro |
| 115 | MGCS36044_01384 | MGCS36089_01372 | ***mgtA*** |  |  | MgtA superfmily cation-translocating P-type ATPase | P | METABOLISM; inorganic ion transport and metabolism | Core | Pro |
| 116 | MGCS36044_01426 | MGCS36089_01414 | ***prfB*** |  |  | peptide chain release factor 2 PrfB | J | INFORMATION STORAGE AND PROCESSING; translation, ribosomal structure and biogenesis | Core | Pro |
| 117 | MGCS36044_01428 | MGCS36089_01416 | ***ftsE*** |  |  | cell division ATP-binding protein FtsE | D | CELLULAR PROCESSES AND SIGNALING; cell division and chromosome partitioning | Core | Pro |
| 118 | MGCS36044_01430 | MGCS36089_01418 | ***ftsX*** |  |  | cell division permease-like protein FtsX | D | CELLULAR PROCESSES AND SIGNALING; cell division and chromosome partitioning | Core | Pro |
| 119 | MGCS36044_01456 | MGCS36089_01444 | ***lacD_1*** |  |  | tagatose-bisphosphate aldolase LacD-like | G | METABOLISM; carbohydrate transport and metabolism | Core | Pro |
| 120 | MGCS36044_01472 | MGCS36089_01460 | ***-*** |  |  | IS30 family transposase |  |  | Core | Pro |
| 121 | MGCS36044_01474 | MGCS36089_01462 | ***nrnA*** |  |  | bifunctional oligoribonuclease/PAP phosphatase NrnA | J | INFORMATION STORAGE AND PROCESSING; translation, ribosomal structure and biogenesis | Core | Pro |
| 122 | MGCS36044_01482 | MGCS36089_01470 | ***-*** |  |  | IS1548 family transposase | L | INFORMATION STORAGE AND PROCESSING; replication, recombination and repair | Core | Pro |
| 123 | MGCS36044_01484 | MGCS36089_01472 | ***fldA*** |  |  | flavodoxin FldA | C | METABOLISM; energy production and conversion | Core | Pro |
| 124 | MGCS36044_01490 | MGCS36089_01478 | ***rplS*** |  |  | 50S ribosomal L19 protein RpsL | J | INFORMATION STORAGE AND PROCESSING; translation, ribosomal structure and biogenesis | Core | Pro |
| 125 | MGCS36044_01496 | MGCS36089_01484 | ***-*** |  |  | HAD-IA family hydrolase | R | General function prediction | Core | Pro |
| 126 | MGCS36044_01498 | MGCS36089_01486 | ***gyrB*** |  |  | DNA topoisomerase ATP-hydrolyzing B subunit GyrB | L | INFORMATION STORAGE AND PROCESSING; replication, recombination and repair | Core | Pro |
| 127 | MGCS36044_01500 | MGCS36089_01488 | ***ezrA*** |  |  | cell division septation ring formation regulator EzrA | D | CELLULAR PROCESSES AND SIGNALING; cell division and chromosome partitioning | Core | Pro |
| 128 | MGCS36044_01506 | MGCS36089_01494 | ***eno*** |  |  | phosphopyruvate hydratase -- enolase protein Eno | G | METABOLISM; carbohydrate transport and metabolism | Core | Pro |
| 129 | MGCS36044_01524 | MGCS36089_01512 | ***sagG*** | Virulence |  | streptolysin S export protein SagG | U | CELLULAR PROCESSES AND SIGNALING; intracellular trafficking, secretion, and vesicular transport | Core | Pro |
| 130 | MGCS36044_01526 | MGCS36089_01514 | ***sagH*** | Virulence |  | streptolysin S export permease protein SagH | U | CELLULAR PROCESSES AND SIGNALING; intracellular trafficking, secretion, and vesicular transport | Core | Pro |
| 131 | MGCS36044_01528 | MGCS36089_01516 | ***sagI*** | Virulence |  | streptolysin S export permease protein SagI | U | CELLULAR PROCESSES AND SIGNALING; intracellular trafficking, secretion, and vesicular transport | Core | Pro |
| 132 | MGCS36044_01536 | MGCS36089_01524 | ***ligA*** |  |  | NAD-dependent DNA ligase LigA | L | INFORMATION STORAGE AND PROCESSING; replication, recombination and repair | Core | Pro |
| 133 | MGCS36044_01538 | MGCS36089_01526 | ***dagK*** |  |  | diacylglycerol kinase family lipid kinase | I | METABOLISM; lipid metabolism | Core | Pro |
| 134 | MGCS36044_01552 | MGCS36089_01540 | ***atpB*** |  |  | ATP synthase A subunit AtpB | C | METABOLISM; energy production and conversion | Core | Pro |
| 135 | MGCS36044_01554 | MGCS36089_01542 | ***atpF*** |  |  | ATP synthase B subunit AtpF | C | METABOLISM; energy production and conversion | Core | Pro |
| 136 | MGCS36044_01556 | MGCS36089_01544 | ***atpH*** |  |  | ATP synthase delta subunit AtpH | C | METABOLISM; energy production and conversion | Core | Pro |
| 137 | MGCS36044_01558 | MGCS36089_01546 | ***atpA*** |  |  | ATP synthase alpha chain, AtpA | C | METABOLISM; energy production and conversion | Core | Pro |
| 138 | MGCS36044_01560 | MGCS36089_01548 | ***atpG*** |  |  | ATP synthase gamma subunit AtpG | C | METABOLISM; energy production and conversion | Core | Pro |
| 139 | MGCS36044_01562 | MGCS36089_01550 | ***atpD*** |  |  | ATP synthase beta subunit AtpD | C | METABOLISM; energy production and conversion | Core | Pro |
| 140 | MGCS36044_01574 | MGCS36089_01562 | ***pheS*** |  |  | phenylalanyl-tRNA synthetase alpha subunit PheS | J | INFORMATION STORAGE AND PROCESSING; translation, ribosomal structure and biogenesis | Core | Pro |
| 141 | MGCS36044_01576 | MGCS36089_01564 | ***pheT*** |  |  | phenylalanyl-tRNA synthetase beta subunit PheT | J | INFORMATION STORAGE AND PROCESSING; translation, ribosomal structure and biogenesis | Core | Pro |
| 142 | MGCS36044_01590 | MGCS36089_01578 | ***rexA*** |  |  | ATP-dependent nuclease A subunit RexA | L | INFORMATION STORAGE AND PROCESSING; replication, recombination and repair | Core | Pro |
| 143 | MGCS36044_01602 | MGCS36089_01590 | ***dnaG*** |  |  | DNA primase protein DnaG | L | INFORMATION STORAGE AND PROCESSING; replication, recombination and repair | Core | Pro |
| 144 | MGCS36044_01604 | MGCS36089_01592 | ***rpoD*** |  |  | RNA polymerase sigma factor RpoD | K | INFORMATION STORAGE AND PROCESSING; transcription | Core | Pro |
| 145 | MGCS36044_01608 | MGCS36089_01596 | ***rmlD*** |  |  | dTDP-4-dehydrorhamnose reductase protein RmlD | G | METABOLISM; carbohydrate transport and metabolism | Core | Pro |
| 146 | MGCS36044_01610 | MGCS36089_01598 | ***rgpA*** |  |  | glycosyltransferase family 1 protein RgpA | G | METABOLISM; carbohydrate transport and metabolism | Core | Pro |
| 147 | MGCS36044_01612 | MGCS36089_01600 | ***rgpB*** |  |  | glycosyltransferase family GT2 protein RgpB | G | METABOLISM; carbohydrate transport and metabolism | Core | Pro |
| 148 | MGCS36044_01614 | MGCS36089_01602 | ***rgpC*** |  |  | ABC transporter polysaccharide/polyol phosphate export permease RgpC | G | METABOLISM; carbohydrate transport and metabolism | Core | Pro |
| 149 | MGCS36044_01616 | MGCS36089_01604 | ***rgpD*** |  |  | ABC transporter polysaccharide/polyol phosphate ATPase component RgpD | G | METABOLISM; carbohydrate transport and metabolism | Core | Pro |
| 150 | MGCS36044_01618 | MGCS36089_01606 | ***rgpE*** |  |  | glycosyltransferase family GT2 protein RgpE | G | METABOLISM; carbohydrate transport and metabolism | Core | Pro |
| 151 | MGCS36044_01620 | MGCS36089_01608 | ***rgpF*** |  |  | alpha-L-Rha alpha-1,3-L-rhamnosyltransferase RgpF | G | METABOLISM; carbohydrate transport and metabolism | Core | Pro |
| 152 | MGCS36044_01626 | MGCS36089_01614 | ***-*** |  |  | DUF2142 domain-containing protein | S | POORLY CHARACTERIZED; function unknown | Core | Pro |
| 153 | MGCS36044_01642 | MGCS36089_01630 | ***cmk*** |  |  | CMP kinase Cmk | F | METABOLISM; nucleotide transport and metabolism | Core | Pro |
| 154 | MGCS36044_01644 | MGCS36089_01632 | ***-*** |  |  | L20_leader |  |  | Core | RNA |
| 155 | MGCS36044_01646 | MGCS36089_01634 | ***infC*** |  |  | translation initiation factor InfC | J | INFORMATION STORAGE AND PROCESSING; translation, ribosomal structure and biogenesis | Core | Pro |
| 156 | MGCS36044_01648 | MGCS36089_01636 | ***rpmI*** |  |  | 50S ribosomal L35 protein RpmL | J | INFORMATION STORAGE AND PROCESSING; translation, ribosomal structure and biogenesis | Core | Pro |
| 157 | MGCS36044_01650 | MGCS36089_01638 | ***rplT*** |  |  | 50S ribosomal L20 protein RplT | J | INFORMATION STORAGE AND PROCESSING; translation, ribosomal structure and biogenesis | Core | Pro |
| 158 | MGCS36044_01652 | MGCS36089_01640 | ***ltaS*** |  |  | LTA synthase LtaS | M | CELLULAR PROCESSES AND SIGNALING; cell envelope biogenesis, outermembrane | Core | Pro |
| 159 | MGCS36044_01674 | MGCS36089_01662 | ***-*** |  |  | IS30 family transposase |  |  | Core | Pro |
| 160 | MGCS36044_01676 | MGCS36089_01664 | ***-*** |  |  | L21_leader |  |  | Core | RNA |
| 161 | MGCS36044_01678 | MGCS36089_01666 | ***rplU*** |  |  | 50S ribosomal L21 protein RplU | J | INFORMATION STORAGE AND PROCESSING; translation, ribosomal structure and biogenesis | Core | Pro |
| 162 | MGCS36044_01680 | MGCS36089_01668 | ***prp*** |  |  | ribosomal-processing cysteine protease Prp | J | INFORMATION STORAGE AND PROCESSING; translation, ribosomal structure and biogenesis | Core | Pro |
| 163 | MGCS36044_01682 | MGCS36089_01670 | ***rpmA*** |  |  | 50S ribosomal L27 protein RpmA | J | INFORMATION STORAGE AND PROCESSING; translation, ribosomal structure and biogenesis | Core | Pro |
| 164 | MGCS36044_01684 | MGCS36089_01672 | ***lysR*** |  |  | LysR family transcriptional regulator | K | INFORMATION STORAGE AND PROCESSING; transcription | Core | Pro |
| 165 | MGCS36044_01714 | MGCS36089_01702 | ***rpsP*** |  |  | 30S ribosomal S16 protein RpsP | J | INFORMATION STORAGE AND PROCESSING; translation, ribosomal structure and biogenesis | Core | Pro |
| 166 | MGCS36044_01720 | MGCS36089_01708 | ***rimM*** |  |  | ribosome maturation factor RimM | J | INFORMATION STORAGE AND PROCESSING; translation, ribosomal structure and biogenesis | Core | Pro |
| 167 | MGCS36044_01722 | MGCS36089_01710 | ***trmD*** |  |  | tRNA (guanosine(37)-N1)-methyltransferase TrmD | J | INFORMATION STORAGE AND PROCESSING; translation, ribosomal structure and biogenesis | Core | Pro |
| 168 | MGCS36044_01746 | MGCS36089_01734 | ***-*** |  |  | tRNA CCA-pyrophosphorylase | J | INFORMATION STORAGE AND PROCESSING; translation, ribosomal structure and biogenesis | Core | Pro |
| 169 | MGCS36044_01766 | MGCS36089_01754 | ***mvaK1*** |  |  | mevalonate kinase MvaK1 | I | METABOLISM; lipid metabolism | Core | Pro |
| 170 | MGCS36044_01768 | MGCS36089_01756 | ***mvaD*** |  |  | diphosphomevalonate decarboxylase MvaD | I | METABOLISM; lipid metabolism | Core | Pro |
| 171 | MGCS36044_01770 | MGCS36089_01758 | ***mvaK2*** |  |  | mevalonate kinase MvaK2 | I | METABOLISM; lipid metabolism | Core | Pro |
| 172 | MGCS36044_01772 | MGCS36089_01760 | ***-*** |  |  | isopentenyl-diphosphate delta-isomerase | I | METABOLISM; lipid metabolism | Core | Pro |
| 173 | MGCS36044_01782 | MGCS36089_01770 | ***mvaS1*** |  |  | hydroxymethylglutaryl-CoA reductase protein (1) MvaS1 | I | METABOLISM; lipid metabolism | Core | Pro |
| 174 | MGCS36044_01784 | MGCS36089_01772 | ***mvaS2*** |  |  | hydroxymethylglutaryl-CoA synthase protein (2) MvaS2 | I | METABOLISM; lipid metabolism | Core | Pro |
| 175 | MGCS36044_01786 | MGCS36089_01774 | ***thyA*** |  |  | thymidylate synthase ThyA | F | METABOLISM; nucleotide transport and metabolism | Core | Pro |
| 176 | MGCS36044_01788 | MGCS36089_01776 | ***dyr*** |  |  | dihydrofolate reductase Dyr | H | METABOLISM; coenzyme transport and metabolism | Core | Pro |
| 177 | MGCS36044_01792 | MGCS36089_01780 | ***clpX*** |  |  | ATP-dependent Clp protease, ATP-binding subunit ClpX | O | CELLULAR PROCESSES AND SIGNALING; post-translational modification, protein turnover, and chaperones | Core | Pro |
| 178 | MGCS36044_01794 | MGCS36089_01782 | ***engB*** |  |  | ribosome biogenesis GTP-binding protein EngB | J | INFORMATION STORAGE AND PROCESSING; translation, ribosomal structure and biogenesis | Core | Pro |
| 179 | MGCS36044_01800 | MGCS36089_01790 | ***-*** |  |  | L10_leader |  |  | Core | RNA |
| 180 | MGCS36044_01804 | MGCS36089_01794 | ***rplL*** |  |  | 50S ribosomal L7/L12 protein RplL | J | INFORMATION STORAGE AND PROCESSING; translation, ribosomal structure and biogenesis | Core | Pro |
| 181 | MGCS36044_01838 | MGCS36089_01828 | ***-*** |  |  | type IV toxin-antitoxin system AbiEi family antitoxin | V | CELLULAR PROCESSES AND SIGNALING; defense mechanisms | RD.4 | Pro |
| 182 | MGCS36044_01920 | MGCS36089_01908 | ***folC*** |  |  | dihydrofolate synthase FolC | H | METABOLISM; coenzyme transport and metabolism | Core | Pro |
| 183 | MGCS36044_01922 | MGCS36089_01910 | ***folE*** |  |  | GTP cyclohydrolase I protein FolE | H | METABOLISM; coenzyme transport and metabolism | Core | Pro |
| 184 | MGCS36044_01924 | MGCS36089_01912 | ***folP*** |  |  | dihydropteroate synthase protein FolP | H | METABOLISM; coenzyme transport and metabolism | Core | Pro |
| 185 | MGCS36044_01926 | MGCS36089_01914 | ***folQ*** |  |  | dihydroneopterin aldolase protein FolB | H | METABOLISM; coenzyme transport and metabolism | Core | Pro |
| 186 | MGCS36044_01928 | MGCS36089_01916 | ***folK*** |  |  | 2-amino-4-hydroxy-6- hydroxymethyldihydropteridine pyrophosphokinase protein FolK | H | METABOLISM; coenzyme transport and metabolism | Core | Pro |
| 187 | MGCS36044_01930 | MGCS36089_01918 | ***murB*** |  |  | UDP-N-acetylmuramate dehydrogenase MurB | M | CELLULAR PROCESSES AND SIGNALING; cell envelope biogenesis, outermembrane | Core | Pro |
| 188 | MGCS36044_01960 | MGCS36089_01948 | ***rex*** |  |  | redox-sensing transcriptional repressor Rex | K | INFORMATION STORAGE AND PROCESSING; transcription | Core | Pro |
| 189 | MGCS36044_01966 | MGCS36089_01954 | ***nifS_2*** |  |  | NifS superfamily cysteine desulfurase | E | METABOLISM; amino acid transport and metabolism | Core | Pro |
| 190 | MGCS36044_01968 | MGCS36089_01956 | ***ribP*** |  |  | ribose-phosphate pyrophosphokinase RibP | F | METABOLISM; Nucleotide transport and metabolism | Core | Pro |
| 191 | MGCS36044_01974 | MGCS36089_01962 | ***nadK*** |  |  | NAD kinase NadK | H | METABOLISM; coenzyme transport and metabolism | Core | Pro |
| 192 | MGCS36044_01978 | MGCS36089_01966 | ***eutD*** |  |  | phosphate acetyltransferase EutD | C | METABOLISM; energy production and conversion | Core | Pro |
| 193 | MGCS36044_02002 | MGCS36089_01990 | ***prfA*** |  |  | peptide chain release factor 1 PrfA | J | INFORMATION STORAGE AND PROCESSING; translation, ribosomal structure and biogenesis | Core | Pro |
| 194 | MGCS36044_02004 | MGCS36089_01992 | ***prmC*** |  |  | peptide chain release factor N(5)-glutamine methyltransferase PrmC | J | INFORMATION STORAGE AND PROCESSING; translation, ribosomal structure and biogenesis | Core | Pro |
| 195 | MGCS36044_02006 | MGCS36089_01994 | ***-*** |  |  | Sua5/YciO/YrdC/YwlC family protein ribosome maturation factor | J | INFORMATION STORAGE AND PROCESSING; translation, ribosomal structure and biogenesis | Core | Pro |
| 196 | MGCS36044_02024 | MGCS36089_02012 | ***ldh*** |  |  | L-lactate dehydrogenase Ldh | C | METABOLISM; energy production and conversion | Core | Pro |
| 197 | MGCS36044_02026 | MGCS36089_02014 | ***gyrA*** |  |  | DNA gyrase subunit A GyrA | L | INFORMATION STORAGE AND PROCESSING; replication, recombination and repair | Core | Pro |
| 198 | MGCS36044_02028 | MGCS36089_02016 | ***srtA*** |  |  | class A sortase SrtA | M | CELLULAR PROCESSES AND SIGNALING; cell envelope biogenesis, outermembrane | Core | Pro |
| 199 | MGCS36044_02038 | MGCS36089_02030 | ***-*** |  |  | disrupted IS3 family transposase encoding gene |  |  | Core | Pro |
| 200 | MGCS36044_02086 | MGCS36089_02078 | ***-*** |  |  | disrupted IS1182 family transposase encoding gene |  |  | Core | Pro |
| 201 | MGCS36044_02136 | MGCS36089_02128 | ***ffh*** |  |  | signal recognition particle protein | U | CELLULAR PROCESSES AND SIGNALING; intracellular trafficking, secretion, and vesicular transport | Core | Pro |
| 202 | MGCS36044_02154 | MGCS36089_02146 | ***xerS*** |  |  | site-specific tyrosine recombinase XerS | L | INFORMATION STORAGE AND PROCESSING; replication, recombination and repair | ROD.6 | Pro |
| 203 | MGCS36044_02200 | MGCS36089_02192 | ***topA*** |  |  | type I DNA topoisomerase TopA | L | INFORMATION STORAGE AND PROCESSING; replication, recombination and repair | Core | Pro |
| 204 | MGCS36044_02206 | MGCS36089_02198 | ***ylqF*** |  |  | ribosome biogenesis GTPase YlqF | J | INFORMATION STORAGE AND PROCESSING; translation, ribosomal structure and biogenesis | Core | Pro |
| 205 | MGCS36044_02212 | MGCS36089_02204 | ***yqfA*** |  |  | membrane channel forming/hemolysin III protein YqfA | K | INFORMATION STORAGE AND PROCESSING; transcription | Core | Pro |
| 206 | MGCS36044_02262 | MGCS36089_02254 | ***coaB*** |  |  | phosphopantothenate--cysteine ligase CoaB | H | METABOLISM; coenzyme transport and metabolism | Core | Pro |
| 207 | MGCS36044_02264 | MGCS36089_02256 | ***coaC*** |  |  | phosphopantothenoylcysteine decarboxylase CoaC | H | METABOLISM; coenzyme transport and metabolism | Core | Pro |
| 208 | MGCS36044_02266 | MGCS36089_02258 | ***panT*** |  |  | pantothenic acid transporter PanT | T | CELLULAR PROCESSES AND SIGNALING; signal transduction mechanisms | Core | Pro |
| 209 | MGCS36044_02268 | MGCS36089_02260 | ***pgmA*** |  |  | phospho-sugar mutase PgmA | G | METABOLISM; carbohydrate transport and metabolism | Core | Pro |
| 210 | MGCS36044_02286 | MGCS36089_02278 | ***coaA*** |  |  | type I pantothenate kinase | H | METABOLISM; coenzyme transport and metabolism | Core | Pro |
| 211 | MGCS36044_02296 | MGCS36089_02288 | ***phoU_2*** |  |  | phosphate signaling complex protein PhoU | P | METABOLISM; inorganic ion transport and metabolism | Core | Pro |
| 212 | MGCS36044_02314 | MGCS36089_02306 | ***spxA_1*** |  |  | transcriptional regulator SpxA | K | INFORMATION STORAGE AND PROCESSING; transcription | Core | Pro |
| 213 | MGCS36044_02316 | MGCS36089_02308 | ***ribF*** |  |  | bifunctional riboflavin kinase/FAD synthetase RibF | H | METABOLISM; coenzyme transport and metabolism | Core | Pro |
| 214 | MGCS36044_02332 | MGCS36089_02324 | ***-*** |  |  | GntR family transcriptional regulator | K | INFORMATION STORAGE AND PROCESSING; transcription | Core | Pro |
| 215 | MGCS36044_02346 | MGCS36089_02338 | ***pcrA*** |  |  | DNA helicase PcrA | L | INFORMATION STORAGE AND PROCESSING; replication, recombination and repair | Core | Pro |
| 216 | MGCS36044_02354 | MGCS36089_02346 | ***-*** |  |  | IS1182 family transposase |  |  | Core | Pro |
| 217 | MGCS36044_02364 | MGCS36089_02356 | ***glmS*** |  |  | glutamine--fructose-6-phosphate transaminase (isomerizing) | E | METABOLISM; amino acid transport and metabolism | Core | Pro |
| 218 | MGCS36044_02370 | MGCS36089_02360 | ***pyk*** |  |  | pyruvate kinase Pyk | G | METABOLISM; carbohydrate transport and metabolism | Core | Pro |
| 219 | MGCS36044_02372 | MGCS36089_02362 | ***pfkA*** |  |  | 6-phosphofructokinase PfkA | G | METABOLISM; carbohydrate transport and metabolism | Core | Pro |
| 220 | MGCS36044_02374 | MGCS36089_02364 | ***dnaE*** |  |  | DNA polymerase III subunit alpha DnaE | L | INFORMATION STORAGE AND PROCESSING; replication, recombination and repair | Core | Pro |
| 221 | MGCS36044_02376 | MGCS36089_02366 | ***yhcF*** |  |  | YhcF family transcriptional regulator | K | INFORMATION STORAGE AND PROCESSING; transcription | Core | Pro |
| 222 | MGCS36044_02384 | MGCS36089_02374 | ***ssrA*** |  |  | transfer-messenger RNA |  |  | Core | RNA |
| 223 | MGCS36044_02392 | MGCS36089_02382 | ***rpsA*** |  |  | 30S ribosomal S1 protein RpsA | J | INFORMATION STORAGE AND PROCESSING; translation, ribosomal structure and biogenesis | Core | Pro |
| 224 | MGCS36044_02402 | MGCS36089_02392 | ***parC*** |  |  | DNA topoisomerase IV subunit A ParC | L | INFORMATION STORAGE AND PROCESSING; replication, recombination and repair | Core | Pro |
| 225 | MGCS36044_02406 | MGCS36089_02396 | ***parE*** |  |  | DNA topoisomerase IV subunit B ParE | L | INFORMATION STORAGE AND PROCESSING; replication, recombination and repair | Core | Pro |
| 226 | MGCS36044_02408 | MGCS36089_02398 | ***plsY*** |  |  | glycerol-3-phosphate 1-O-acyltransferase PlsY | L | INFORMATION STORAGE AND PROCESSING; replication, recombination and repair | Core | Pro |
| 227 | MGCS36044_02446 | MGCS36089_02436 | ***mnmE*** |  |  | MnmE family tRNA uridine-5-carboxymethylaminomethyl(34) synthesis GTPase | J | INFORMATION STORAGE AND PROCESSING; translation, ribosomal structure and biogenesis | Core | Pro |
| 228 | MGCS36044_02528 | MGCS36089_02542 | ***glmM*** |  |  | phosphoglucosamine mutase GlmM | G | METABOLISM; carbohydrate transport and metabolism | Core | Pro |
| 229 | MGCS36044_02532 | MGCS36089_02546 | ***-*** |  |  | DisA N domain-containing diadenylate cyclase | G | METABOLISM; carbohydrate transport and metabolism | Core | Pro |
| 230 | MGCS36044_02534 | MGCS36089_02548 | ***murE_2*** |  |  | UDP-N-acetylmuramoylalanyl-D-glutamate-2, 6-diaminopimelate ligase MurE | M | CELLULAR PROCESSES AND SIGNALING; cell envelope biogenesis, outermembrane | Core | Pro |
| 231 | MGCS36044_02536 | MGCS36089_02550 | ***-*** |  |  | CobQ-like type 1 glutamine amidotransferase | M | CELLULAR PROCESSES AND SIGNALING; cell envelope biogenesis, outermembrane | Core | Pro |
| 232 | MGCS36044_02538 | MGCS36089_02552 | ***lplA_2*** |  |  | lipoate--protein ligase LplA | H | METABOLISM; coenzyme transport and metabolism | Core | Pro |
| 233 | MGCS36044_02544 | MGCS36089_02558 | ***acoL*** |  |  | dihydrolipoyl dehydrogenase AcoL | C | METABOLISM; energy production and conversion | Core | Pro |
| 234 | MGCS36044_02548 | MGCS36089_02562 | ***acoC*** |  |  | dihydrolipoamide acetyltransferase AcoC | C | METABOLISM; energy production and conversion | Core | Pro |
| 235 | MGCS36044_02550 | MGCS36089_02564 | ***acoB*** |  |  | pyruvate dehydrogenase E1 component beta subunit AcoB | C | METABOLISM; energy production and conversion | Core | Pro |
| 236 | MGCS36044_02552 | MGCS36089_02566 | ***acoA*** |  |  | Pyruvate dehydrogenase E1 component alpha subunit AcoA | C | METABOLISM; energy production and conversion | Core | Pro |
| 237 | MGCS36044_02560 | MGCS36089_02574 | ***rnjA_1*** |  |  | mRNA degradation ribonuclease RnjA | A | INFORMATION STORAGE AND PROCESSING; RNA processing and modification | Core | Pro |
| 238 | MGCS36044_02578 | MGCS36089_02592 | ***-*** |  |  | disrupted IS3 family transposase encoding gene |  |  | Core | Pro |
| 239 | MGCS36044_02596 | MGCS36089_02608 | ***rfbB*** |  |  | dTDP-glucose 4,6-dehydratase RfbB | M | CELLULAR PROCESSES AND SIGNALING; cell envelope biogenesis, outermembrane | Core | Pro |
| 240 | MGCS36044_02598 | MGCS36089_02610 | ***rfbC*** |  |  | dTDP-4-dehydrorhamnose 3,5-epimerase RfbC | G | METABOLISM; carbohydrate transport and metabolism | Core | Pro |
| 241 | MGCS36044_02600 | MGCS36089_02612 | ***rfbA*** |  |  | glucose-1-phosphate thymidylyltransferase RfbA | M | CELLULAR PROCESSES AND SIGNALING; cell envelope biogenesis, outermembrane | Core | Pro |
| 242 | MGCS36044_02620 | MGCS36089_02632 | ***rnz*** |  |  | ribonuclease Rnz | J | INFORMATION STORAGE AND PROCESSING; translation, ribosomal structure and biogenesis | Core | Pro |
| 243 | MGCS36044_02644 | MGCS36089_02656 | ***malQ*** |  |  | 4-alpha-glucanotransferase (amylomaltase) protein MalQ | G | METABOLISM; carbohydrate transport and metabolism | Core | Pro |
| 244 | MGCS36044_02648 | MGCS36089_02660 | ***-*** |  |  | IS30 family transposase |  |  | Core | Pro |
| 245 | MGCS36044_02672 | MGCS36089_02684 | ***dltB*** |  |  | D-alanyl-lipoteichoic acid biosynthesis protein DltB | M | CELLULAR PROCESSES AND SIGNALING; cell envelope biogenesis, outermembrane | Core | Pro |
| 246 | MGCS36044_02674 | MGCS36089_02686 | ***dltA*** |  |  | D-alanine--poly(phosphoribitol) ligase subunit DltA | M | CELLULAR PROCESSES AND SIGNALING; cell envelope biogenesis, outermembrane | Core | Pro |
| 247 | MGCS36044_02704 | MGCS36089_02716 | ***obgE*** |  |  | GTPase ObgE | L | INFORMATION STORAGE AND PROCESSING; replication, recombination and repair | Core | Pro |
| 248 | MGCS36044_02834 | MGCS36089_02846 | ***map*** |  |  | methionyl aminopeptidase Map | J | INFORMATION STORAGE AND PROCESSING; translation, ribosomal structure and biogenesis | Core | Pro |
| 249 | MGCS36044_02864 | MGCS36089_02876 | ***metK*** |  |  | methionine adenosyltransferase MetK | J | INFORMATION STORAGE AND PROCESSING; translation, ribosomal structure and biogenesis | Core | Pro |
| 250 | MGCS36044_02870 | MGCS36089_02882 | ***dnaX*** |  |  | DNA polymerase III gamma/tau subunit DnaX | L | INFORMATION STORAGE AND PROCESSING; replication, recombination and repair | Core | Pro |
| 251 | MGCS36044_02880 | MGCS36089_02892 | ***srmB*** |  |  | SrmB superfailily II DNA and RNA helicase | L | INFORMATION STORAGE AND PROCESSING; replication, recombination and repair | Core | Pro |
| 252 | MGCS36044_02884 | MGCS36089_02896 | ***gapN*** |  |  | NADP-dependent glyceraldehyde-3-phosphate dehydrogenase GapN | G | METABOLISM; carbohydrate transport and metabolism | Core | Pro |
| 253 | MGCS36044_02888 | MGCS36089_02900 | ***ptsH*** |  |  | PTS transporter phosphocarrier protein PtsH | G | METABOLISM; carbohydrate transport and metabolism | Core | Pro |
| 254 | MGCS36044_02890 | MGCS36089_02902 | ***nrdH*** |  |  | glutaredoxin-like protein NrdH | O | CELLULAR PROCESSES AND SIGNALING; post-translational modification, protein turnover, and chaperones | Core | Pro |
| 255 | MGCS36044_02892 | MGCS36089_02904 | ***nrdE_2*** |  |  | class 1b ribonucleoside-diphosphate reductase alpha subunit NrdE | F | METABOLISM; nucleotide transport and metabolism | Core | Pro |
| 256 | MGCS36044_02894 | MGCS36089_02906 | ***nrdF_2*** |  |  | class 1b ribonucleoside-diphosphate reductase beta subunit NrdF | F | METABOLISM; nucleotide transport and metabolism | Core | Pro |
| 257 | MGCS36044_02904 | MGCS36089_02916 | ***alaS*** |  |  | alanine--tRNA synthetase AlaS | J | INFORMATION STORAGE AND PROCESSING; translation, ribosomal structure and biogenesis | Core | Pro |
| 258 | MGCS36044_02936 | MGCS36089_02948 | ***sodA*** |  |  | superoxide dismutase SodA | V | CELLULAR PROCESSES AND SIGNALING; defense mechanisms | Core | Pro |
| 259 | MGCS36044_02944 | MGCS36089_02956 | ***plsC*** |  | Secreted | secreted 1-acyl-sn-glycerol-3-phosphate acyltransferase PlsC | I | METABOLISM; lipid metabolism | Core | Pro |
| 260 | MGCS36044_02966 | MGCS36089_02978 | ***murF*** |  |  | UDP-N-acetylmuramoyl-tripeptide--D-alanyl-D- alanine ligase MurF | M | CELLULAR PROCESSES AND SIGNALING; cell envelope biogenesis, outermembrane | Core | Pro |
| 261 | MGCS36044_02968 | MGCS36089_02980 | ***ddl*** |  |  | D-alanine--D-alanine ligase Ddl | M | CELLULAR PROCESSES AND SIGNALING; cell envelope biogenesis, outermembrane | Core | Pro |
| 262 | MGCS36044_02970 | MGCS36089_02982 | ***recR*** |  |  | recombination mediator RecR | L | INFORMATION STORAGE AND PROCESSING; replication, recombination and repair | Core | Pro |
| 263 | MGCS36044_02980 | MGCS36089_02992 | ***gpmA*** |  |  | phosphoglycerate mutase GpmA | G | METABOLISM; carbohydrate transport and metabolism | Core | Pro |
| 264 | MGCS36044_02982 | MGCS36089_02994 | ***-*** |  |  | IS982 family transposase |  |  | Core | Pro |
| 265 | MGCS36044_02994 | MGCS36089_03006 | ***-*** |  |  | DNA-binding protein HU | L | INFORMATION STORAGE AND PROCESSING; replication, recombination and repair | Core | Pro |
| 266 | MGCS36044_03008 | MGCS36089_03020 | ***tlyA*** |  |  | TlyA family RNA methyltransferase | J | INFORMATION STORAGE AND PROCESSING; translation, ribosomal structure and biogenesis | Core | Pro |
| 267 | MGCS36044_03014 | MGCS36089_03026 | ***xseA*** |  |  | exodeoxyribonuclease VII large subunit XseA | L | INFORMATION STORAGE AND PROCESSING; replication, recombination and repair | Core | Pro |
| 268 | MGCS36044_03044 | MGCS36089_03056 | ***ileS*** |  |  | isoleucine--tRNA synthetase IleS | J | INFORMATION STORAGE AND PROCESSING; translation, ribosomal structure and biogenesis | Core | Pro |
| 269 | MGCS36044_03046 | MGCS36089_03058 | ***divIVA*** |  |  | cell division protein DivIVA | D | CELLULAR PROCESSES AND SIGNALING; cell division and chromosome partitioning | Core | Pro |
| 270 | MGCS36044_03048 | MGCS36089_03060 | ***-*** |  |  | RNA-binding protein | R | General function prediction only | Core | Pro |
| 271 | MGCS36044_03050 | MGCS36089_03062 | ***-*** |  |  | YggT family protein | S | POORLY CHARACTERIZED; function unknown | Core | Pro |
| 272 | MGCS36044_03054 | MGCS36089_03066 | ***yggS*** |  |  | YggS family pyridoxal phosphate-dependent enzyme | F | METABOLISM; nucleotide transport and metabolism | Core | Pro |
| 273 | MGCS36044_03056 | MGCS36089_03068 | ***ftsZ*** |  |  | cell division protein FtsZ | D | CELLULAR PROCESSES AND SIGNALING; cell division and chromosome partitioning | Core | Pro |
| 274 | MGCS36044_03058 | MGCS36089_03070 | ***ftsA*** |  |  | cell division protein FtsA | D | CELLULAR PROCESSES AND SIGNALING; cell division and chromosome partitioning | Core | Pro |
| 275 | MGCS36044_03060 | MGCS36089_03072 | ***ftsQ*** |  |  | cell division protein FtsQ/DivIB | D | CELLULAR PROCESSES AND SIGNALING; cell division and chromosome partitioning | Core | Pro |
| 276 | MGCS36044_03062 | MGCS36089_03074 | ***murG*** |  |  | UDP-N-acetylglucosamine--N-acetylmuramyl- (pentapeptide) pyrophosphoryl-undecaprenol N-acetylglucosamine transferase MurG | M | CELLULAR PROCESSES AND SIGNALING; cell envelope biogenesis, outermembrane | Core | Pro |
| 277 | MGCS36044_03070 | MGCS36089_03082 | ***typA*** |  |  | translational GTPase TypA | T | CELLULAR PROCESSES AND SIGNALING; signal transduction mechanisms | Core | Pro |
| 278 | MGCS36044_03092 | MGCS36089_03104 | ***-*** |  |  | IS1548 family transposase | L | INFORMATION STORAGE AND PROCESSING; replication, recombination and repair | Core | Pro |
| 279 | MGCS36044_03096 | MGCS36089_03108 | ***-*** |  |  | IS110 family transposase |  |  | Core | Pro |
| 280 | MGCS36044_03120 | MGCS36089_03132 | ***-*** |  |  | IS110 family transposase |  |  | Core | Pro |
| 281 | MGCS36044_03170 | MGCS36089_03180 | ***valS*** |  |  | valine--tRNA synthetase ValS | J | INFORMATION STORAGE AND PROCESSING; translation, ribosomal structure and biogenesis | Core | Pro |
| 282 | MGCS36044_03180 | MGCS36089_03190 | ***-*** |  |  | DUF1912 family protein | S | POORLY CHARACTERIZED; function unknown | Core | Pro |
| 283 | MGCS36044_03278 | MGCS36089_03290 | ***-*** |  |  | IS30 family transposase |  |  | Core | Pro |
| 284 | MGCS36044_03300 | MGCS36089_03312 | ***pknB*** |  |  | Stk1 family PASTA domain-containing Ser/Thr kinase | T | CELLULAR PROCESSES AND SIGNALING; signal transduction mechanisms | Core | Pro |
| 285 | MGCS36044_03306 | MGCS36089_03318 | ***fmt*** |  |  | methionyl-tRNA formyl transferase Fmt | J | INFORMATION STORAGE AND PROCESSING; translation, ribosomal structure and biogenesis | Core | Pro |
| 286 | MGCS36044_03308 | MGCS36089_03320 | ***priA*** |  |  | primosomal protein PriA | L | INFORMATION STORAGE AND PROCESSING; replication, recombination and repair | Core | Pro |
| 287 | MGCS36044_03310 | MGCS36089_03322 | ***rpoZ*** |  |  | DNA-directed RNA polymerase omega subunit RpoZ | K | INFORMATION STORAGE AND PROCESSING; transcription | Core | Pro |
| 288 | MGCS36044_03312 | MGCS36089_03324 | ***gmk*** |  |  | guanylate kinase Gmk | F | METABOLISM; nucleotide transport and metabolism | Core | Pro |
| 289 | MGCS36044_03314 | MGCS36089_03326 | ***rny*** |  |  | ribonuclease (Y) Rny | D | CELLULAR PROCESSES AND SIGNALING; cell division and chromosome partitioning | Core | Pro |
| 290 | MGCS36044_03330 | MGCS36089_03342 | ***mapZ*** |  |  | MapZ family cell division site-positioning protein | D | CELLULAR PROCESSES AND SIGNALING; cell division and chromosome partitioning | Core | Pro |
| 291 | MGCS36044_03334 | MGCS36089_03346 | ***-*** |  |  | RNaseP_bact_b |  |  | Core | RNA |
| 292 | MGCS36044_03336 | MGCS36089_03348 | ***gpsB*** |  |  | cell division regulator GpsB | D | CELLULAR PROCESSES AND SIGNALING; cell division and chromosome partitioning | Core | Pro |
| 293 | MGCS36044_03340 | MGCS36089_03352 | ***recU*** |  |  | Holliday junction resolvase RecU | L | INFORMATION STORAGE AND PROCESSING; DNA replication, recombination and repair | Core | Pro |
| 294 | MGCS36044_03342 | MGCS36089_03354 | ***pbp1A*** |  |  | bifunctional PG transglycosylase-transpeptidase classA penicillin-binding protein PBP1A | M | CELLULAR PROCESSES AND SIGNALING; cell envelope biogenesis, outermembrane | Core | Pro |
| 295 | MGCS36044_03346 | MGCS36089_03358 | ***nadE*** |  |  | ammonia-dependent NAD(+) synthetase NadE | H | METABOLISM; coenzyme transport and metabolism | Core | Pro |
| 296 | MGCS36044_03348 | MGCS36089_03360 | ***pncB*** |  |  | nicotinate phosphoribosyltransferase PncB | H | METABOLISM; coenzyme transport and metabolism | Core | Pro |
| 297 | MGCS36044_03352 | MGCS36089_03364 | ***trxB_2*** |  |  | thioredoxin-disulfide reductase TrxB | C | METABOLISM; energy production and conversion | Core | Pro |
| 298 | MGCS36044_03354 | MGCS36089_03366 | ***-*** |  |  | DUF4059 family protein | S | POORLY CHARACTERIZED; function unknown | Core | Pro |
| 299 | MGCS36044_03356 | MGCS36089_03368 | ***-*** |  |  | GlnQ family polar amino acid ABC transporter ATPase | E | METABOLISM; amino acid transport and metabolism | Core | Pro |
| 300 | MGCS36044_03358 | MGCS36089_03370 | ***hisM*** |  |  | HisM family amino acid ABC transporter permease | P | METABOLISM; inorganic ion transport and metabolism | Core | Pro |
| 301 | MGCS36044_03362 | MGCS36089_03374 | ***cshB*** |  |  | DEAD/DEAH box helicase | L | INFORMATION STORAGE AND PROCESSING; replication, recombination and repair | Core | Pro |
| 302 | MGCS36044_03364 | MGCS36089_03376 | ***mraY*** |  |  | phospho-N-acetylmuramoyl-pentapeptide- translocase MraY | M | CELLULAR PROCESSES AND SIGNALING; cell envelope biogenesis, outermembrane | Core | Pro |
| 303 | MGCS36044_03366 | MGCS36089_03378 | ***pbp2X*** |  |  | PG transpeptidase class B penicillin-binding protein PBP2X/FtsI | M | CELLULAR PROCESSES AND SIGNALING; cell envelope biogenesis, outermembrane | Core | Pro |
| 304 | MGCS36044_03370 | MGCS36089_03382 | ***mraW*** |  |  | S-adenosyl-methyltransferase MraW | D | CELLULAR PROCESSES AND SIGNALING; Cell cycle control, cell division, chromosome partitioning | Core | Pro |
| 305 | MGCS36044_03380 | MGCS36089_03392 | ***-*** |  |  | disrupted IS3 family transposase encoding gene |  |  | Core | Pro |
| 306 | MGCS36044_03382 | MGCS36089_03394 | ***tkt*** |  |  | transketolase Tkt | G | METABOLISM; carbohydrate transport and metabolism | Core | Pro |
| 307 | MGCS36044_03398 | MGCS36089_03410 | ***ynzC*** |  |  | DUF896 family protein | S | POORLY CHARACTERIZED; function unknown | Core | Pro |
| 308 | MGCS36044_03400 | MGCS36089_03412 | ***glyS*** |  |  | glycine--tRNA ligase beta subunit GlyS | J | INFORMATION STORAGE AND PROCESSING; translation, ribosomal structure and biogenesis | Core | Pro |
| 309 | MGCS36044_03402 | MGCS36089_03414 | ***glyQ*** |  |  | glycine--tRNA ligase alpha subunit GlyQ | J | INFORMATION STORAGE AND PROCESSING; translation, ribosomal structure and biogenesis | Core | Pro |
| 310 | MGCS36044_03414 | MGCS36089_03426 | ***degV_2*** |  |  | DegV family fatty acid-binding protein | I | METABOLISM; lipid metabolism | Core | Pro |
| 311 | MGCS36044_03464 | MGCS36089_03476 | ***infB*** |  |  | translation initiation factor IF-2 | J | INFORMATION STORAGE AND PROCESSING; translation, ribosomal structure and biogenesis | Core | Pro |
| 312 | MGCS36044_03466 | MGCS36089_03478 | ***-*** |  |  | YlxQ-related RNA-binding protein | J | INFORMATION STORAGE AND PROCESSING; translation, ribosomal structure and biogenesis | Core | Pro |
| 313 | MGCS36044_03468 | MGCS36089_03480 | ***-*** |  |  | YlxR family putative RNA-binding protein | K | INFORMATION STORAGE AND PROCESSING; transcription | Core | Pro |
| 314 | MGCS36044_03470 | MGCS36089_03482 | ***nusA*** |  |  | transcription termination factor NusA | K | INFORMATION STORAGE AND PROCESSING; transcription | Core | Pro |
| 315 | MGCS36044_03478 | MGCS36089_03490 | ***cotS*** |  |  | CotS family thaimine kinase | M | CELLULAR PROCESSES AND SIGNALING; cell envelope biogenesis, outermembrane | Core | Pro |
| 316 | MGCS36044_03512 | MGCS36089_03524 | ***serS*** |  |  | seryl-tRNA synthetase SerS | J | INFORMATION STORAGE AND PROCESSING; translation, ribosomal structure and biogenesis | Core | Pro |
| 317 | MGCS36044_03514 | MGCS36089_03526 | ***accD*** |  |  | acetyl-CoA carboxylase carboxyl transferase alpha subunit AccD | I | METABOLISM; lipid metabolism | Core | Pro |
| 318 | MGCS36044_03516 | MGCS36089_03528 | ***accA*** |  |  | acetyl-CoA carboxylase, carboxyltransferase beta subunit AccA | I | METABOLISM; lipid metabolism | Core | Pro |
| 319 | MGCS36044_03518 | MGCS36089_03530 | ***accC*** |  |  | acetyl-CoA carboxylase biotin carboxylase subunit AccC | I | METABOLISM; lipid metabolism | Core | Pro |
| 320 | MGCS36044_03520 | MGCS36089_03532 | ***fabZ*** |  |  | 3-hydroxyacyl-ACP dehydratase FabZ | I | METABOLISM; lipid metabolism | Core | Pro |
| 321 | MGCS36044_03524 | MGCS36089_03536 | ***fabF*** |  |  | 3-oxoacyl-[acyl-carrier-protein] synthase protein FabF | I | METABOLISM; lipid metabolism | Core | Pro |
| 322 | MGCS36044_03528 | MGCS36089_03540 | ***fabD*** |  |  | malonyl CoA-acyl carrier protein transacylase protein FabD | I | METABOLISM; lipid metabolism | Core | Pro |
| 323 | MGCS36044_03530 | MGCS36089_03542 | ***fabK*** |  |  | Enoyl-[acyl-carrier-protein] reductase protein FabK | I | METABOLISM; lipid metabolism | Core | Pro |
| 324 | MGCS36044_03536 | MGCS36089_03548 | ***fabT*** |  |  | transcriptional regulatory protein FabT | I | METABOLISM; lipid metabolism | Core | Pro |
| 325 | MGCS36044_03538 | MGCS36089_03550 | ***phaB*** |  |  | enoyl-CoA hydratase protein PhaB | I | METABOLISM; lipid metabolism | Core | Pro |
| 326 | MGCS36044_03540 | MGCS36089_03552 | ***dnaJ*** |  |  | chaperone protein DnaJ | O | CELLULAR PROCESSES AND SIGNALING; post-translational modification, protein turnover, and chaperones | Core | Pro |
| 327 | MGCS36044_03544 | MGCS36089_03556 | ***dnaK*** |  |  | molecular chaperone DnaK | O | CELLULAR PROCESSES AND SIGNALING; post-translational modification, protein turnover, and chaperones | Core | Pro |
| 328 | MGCS36044_03546 | MGCS36089_03558 | ***grpE*** |  |  | heat shock protein/nucleotide exchange factor GrpE | O | CELLULAR PROCESSES AND SIGNALING; post-translational modification, protein turnover, and chaperones | Core | Pro |
| 329 | MGCS36044_03548 | MGCS36089_03560 | ***hrcA*** |  |  | heat-inducible transcriptional repressor HrcA | K | INFORMATION STORAGE AND PROCESSING; transcription | Core | Pro |
| 330 | MGCS36044_03560 | MGCS36089_03572 | ***gatB_2*** |  |  | aspartyl-tRNA(Asn) or glutamyl-tRNA(Gln) amidotransferase B subunit GatB | J | INFORMATION STORAGE AND PROCESSING; translation, ribosomal structure and biogenesis | Core | Pro |
| 331 | MGCS36044_03562 | MGCS36089_03574 | ***gatA_2*** |  |  | aspartyl-tRNA(Asn) or glutamyl-tRNA(Gln) amidotransferase A subunit GatA | J | INFORMATION STORAGE AND PROCESSING; translation, ribosomal structure and biogenesis | Core | Pro |
| 332 | MGCS36044_03564 | MGCS36089_03576 | ***gatC_2*** |  |  | aspartyl-tRNA(Asn) or glutamyl-tRNA(Gln) amidotransferase C subunit GatC | J | INFORMATION STORAGE AND PROCESSING; translation, ribosomal structure and biogenesis | Core | Pro |
| 333 | MGCS36044_03582 | MGCS36089_03594 | ***codY*** |  |  | CodY family GTP-sensing pleiotropic transcriptional regulator | K | INFORMATION STORAGE AND PROCESSING; transcription | Core | Pro |
| 334 | MGCS36044_03594 | MGCS36089_03606 | ***recG*** |  |  | ATP-dependent DNA helicase RecG | L | INFORMATION STORAGE AND PROCESSING; replication, recombination and repair | Core | Pro |
| 335 | MGCS36044_03620 | MGCS36089_03632 | ***acpS*** |  |  | AcpS family provisional 4'-phosphopantetheinyl transferase | I | METABOLISM; lipid metabolism | Core | Pro |
| 336 | MGCS36044_03622 | MGCS36089_03634 | ***secA*** |  |  | preprotein translocase subunit SecA | U | CELLULAR PROCESSES AND SIGNALING; intracellular trafficking, secretion, and vesicular transport | Core | Pro |
| 337 | MGCS36044_03632 | MGCS36089_03644 | ***scrB*** |  |  | sucrose-6-phosphate hydrolase ScrB | G | METABOLISM; carbohydrate transport and metabolism | Core | Pro |
| 338 | MGCS36044_03640 | MGCS36089_03652 | ***efp*** |  |  | translation elongation factor (P) Efp | J | INFORMATION STORAGE AND PROCESSING; translation, ribosomal structure and biogenesis | Core | Pro |
| 339 | MGCS36044_03652 | MGCS36089_03664 | ***rpsR*** |  |  | 30S ribosomal S18 protein RpsR | J | INFORMATION STORAGE AND PROCESSING; translation, ribosomal structure and biogenesis | Core | Pro |
| 340 | MGCS36044_03654 | MGCS36089_03666 | ***ssb_2*** |  |  | single-stranded DNA-binding protein | L | INFORMATION STORAGE AND PROCESSING; replication, recombination and repair | Core | Pro |
| 341 | MGCS36044_03656 | MGCS36089_03668 | ***rpsF*** |  |  | 30S ribosomal S6 protein RpsF | J | INFORMATION STORAGE AND PROCESSING; translation, ribosomal structure and biogenesis | Core | Pro |
| 342 | MGCS36044_03662 | MGCS36089_03674 | ***trxA_2*** |  |  | thioredoxin TrxA | O | CELLULAR PROCESSES AND SIGNALING; post-translational modification, protein turnover, and chaperones | Core | Pro |
| 343 | MGCS36044_03668 | MGCS36089_03680 | ***-*** |  |  | colicin V production family protein | O | CELLULAR PROCESSES AND SIGNALING; post-translational modification, protein turnover, and chaperones | Core | Pro |
| 344 | MGCS36044_03672 | MGCS36089_03684 | ***rnhC*** |  |  | HIII ribonuclease RnhC | L | INFORMATION STORAGE AND PROCESSING; replication, recombination and repair | Core | Pro |
| 345 | MGCS36044_03674 | MGCS36089_03686 | ***lepB_2*** |  |  | signal peptidase I LepB | U | CELLULAR PROCESSES AND SIGNALING; intracellular trafficking, secretion, and vesicular transport | Core | Pro |
| 346 | MGCS36044_03704 | MGCS36089_03716 | ***glpF_2*** |  |  | glycerol uptake facilitator GlpF | G | METABOLISM; carbohydrate transport and metabolism | Core | Pro |
| 347 | MGCS36044_03770 | MGCS36089_03782 | ***-*** |  |  | DUF536 domain-containing protein | S | POORLY CHARACTERIZED; function unknown | Core | Pro |
| 348 | MGCS36044_03778 | MGCS36089_03790 | ***tsaD*** |  |  | tRNA (adenosine(37)-N6)-threonylcarbamoyltransferase complex transferase subunit TsaD | J | INFORMATION STORAGE AND PROCESSING; translation, ribosomal structure and biogenesis | Core | Pro |
| 349 | MGCS36044_03782 | MGCS36089_03794 | ***tsaB*** |  |  | tRNA (adenosine(37)-N6)-threonylcarbamoyltransferase complex dimerization subunit type 1 TsaB | O | CELLULAR PROCESSES AND SIGNALING; post-translational modification, protein turnover, and chaperones | Core | Pro |
| 350 | MGCS36044_03784 | MGCS36089_03796 | ***-*** |  |  | DUF1447 family protein | S | POORLY CHARACTERIZED; function unknown | Core | Pro |
| 351 | MGCS36044_03786 | MGCS36089_03798 | ***rnjA_2*** |  |  | mRNA degradation ribonuclease RnjA | A | INFORMATION STORAGE AND PROCESSING; RNA processing and modification | Core | Pro |
| 352 | MGCS36044_03804 | MGCS36089_03816 | ***pgk*** |  |  | phosphoglycerate kinase Pgk | G | METABOLISM; carbohydrate transport and metabolism | Core | Pro |
| 353 | MGCS36044_03808 | MGCS36089_03820 | ***gapA*** |  |  | glyceraldehyde-3-phosphate dehydrogenase GapA | G | METABOLISM; carbohydrate transport and metabolism | Core | Pro |
| 354 | MGCS36044_03810 | MGCS36089_03822 | ***fusA*** |  |  | FusA family elongation factor EF-G | J | INFORMATION STORAGE AND PROCESSING; translation, ribosomal structure and biogenesis | Core | Pro |
| 355 | MGCS36044_03812 | MGCS36089_03824 | ***rpsG*** |  |  | 30S ribosomal S7 protein RpsG | J | INFORMATION STORAGE AND PROCESSING; translation, ribosomal structure and biogenesis | Core | Pro |
| 356 | MGCS36044_03814 | MGCS36089_03826 | ***rpsL*** |  |  | 30S ribosomal S12 protein RpsL | J | INFORMATION STORAGE AND PROCESSING; translation, ribosomal structure and biogenesis | Core | Pro |
| 357 | MGCS36044_03828 | MGCS36089_03840 | ***rsgA*** |  |  | ribosome small subunit-dependent GTPase (A) RgsA | J | INFORMATION STORAGE AND PROCESSING; translation, ribosomal structure and biogenesis | Core | Pro |
| 358 | MGCS36044_03834 | MGCS36089_03846 | ***rrmV*** |  |  | 5S rRNA maturation endonuclease RnmV | J | INFORMATION STORAGE AND PROCESSING; translation, ribosomal structure and biogenesis | Core | Pro |
| 359 | MGCS36044_03858 | MGCS36089_03870 | ***rpmH*** |  |  | 50S ribosomal L34 protein RpmH | J | INFORMATION STORAGE AND PROCESSING; translation, ribosomal structure and biogenesis | Core | Pro |
| 360 | MGCS36044_03864 | MGCS36089_03876 | ***rnpA*** |  |  | ribonuclease P protein component RnpA | J | INFORMATION STORAGE AND PROCESSING; translation, ribosomal structure and biogenesis | Core | Pro |
| 361 | MGCS36044_03874 | MGCS36089_03886 | ***gltX*** |  |  | glutamate--tRNA ligase | J | INFORMATION STORAGE AND PROCESSING; translation, ribosomal structure and biogenesis | Core | Pro |
| 362 | MGCS36044_03884 | MGCS36089_03896 | ***-*** |  |  | carbonic anhydrase | P | METABOLISM; inorganic ion transport and metabolism | Core | Pro |
| 363 | MGCS36044_03898 | MGCS36089_03910 | ***gpsA*** |  |  | NAD(P)H-dependent glycerol-3-phosphate dehydrogenase GpsA | I | METABOLISM; lipid metabolism | Core | Pro |
| 364 | MGCS36044_03900 | MGCS36089_03912 | ***galU*** |  |  | UTP--glucose-1-phosphate uridylyltransferase GalU | M | CELLULAR PROCESSES AND SIGNALING; cell envelope biogenesis, outermembrane | Core | Pro |
| 365 | MGCS36044_03920 | MGCS36089_03932 | ***pgi*** |  |  | Pgi family glucose-6-phosphate isomerase | G | METABOLISM; carbohydrate transport and metabolism | Core | Pro |
| 366 | MGCS36044_03928 | MGCS36089_03940 | ***-*** |  |  | Bacteria_small_SRP |  |  | Core | RNA |
| 367 | MGCS36044_03974 | MGCS36089_03986 | ***leuS*** |  |  | leucine--tRNA synthase LeuS | J | INFORMATION STORAGE AND PROCESSING; translation, ribosomal structure and biogenesis | Core | Pro |
| 368 | MGCS36044_03996 | MGCS36089_04010 | ***ifs*** |  |  | nicotine adenine dinucleotide glycohydrolase inhibitor Ifs | R | General function prediction only | Core | Pro |
| 369 | MGCS36044_04002 | MGCS36089_04016 | ***secE*** |  |  | preprotein translocase subunit protein SecE | U | CELLULAR PROCESSES AND SIGNALING; intracellular trafficking, secretion, and vesicular transport | Core | Pro |
| 370 | MGCS36044_04018 | MGCS36089_04032 | ***groEL*** |  |  | chaperonin GroEL | O | CELLULAR PROCESSES AND SIGNALING; post-translational modification, protein turnover, and chaperones | Core | Pro |
| 371 | MGCS36044_04020 | MGCS36089_04034 | ***groES*** |  |  | co-chaperone GroES | O | CELLULAR PROCESSES AND SIGNALING; post-translational modification, protein turnover, and chaperones | Core | Pro |
| 372 | MGCS36044_04032 | MGCS36089_04046 | ***-*** |  |  | disrupted IS1182 family transposase encoding gene |  |  | Core | Pro |
| 373 | MGCS36044_04060 | MGCS36089_04074 | ***rpsB*** |  |  | 30S ribosomal S2 protein RpsB | J | INFORMATION STORAGE AND PROCESSING; translation, ribosomal structure and biogenesis | Core | Pro |
| 374 | MGCS36044_04062 | MGCS36089_04076 | ***tsf*** |  |  | translation elongation factor Tsf | J | INFORMATION STORAGE AND PROCESSING; translation, ribosomal structure and biogenesis | Core | Pro |
| 375 | MGCS36044_04066 | MGCS36089_04080 | ***treC*** |  |  | trehalose-6-phosphate hydrolase TreC | G | METABOLISM; carbohydrate transport and metabolism | Core | Pro |
| 376 | MGCS36044_04096 | MGCS36089_04110 | ***-*** |  |  | DUF1292 domain-containing protein | S | POORLY CHARACTERIZED; function unknown | Core | Pro |
| 377 | MGCS36044_04098 | MGCS36089_04112 | ***ruvX*** |  |  | Holliday junction resolvase RuvX | K | INFORMATION STORAGE AND PROCESSING; transcription | Core | Pro |
| 378 | MGCS36044_04100 | MGCS36089_04114 | ***-*** |  |  | IreB-related regulatory phosphoprotein | S | POORLY CHARACTERIZED; function unknown | Core | Pro |
| 379 | MGCS36044_04104 | MGCS36089_04118 | ***spxA_2*** |  |  | transcriptional regulator SpxA | P | METABOLISM; inorganic ion transport and metabolism | Core | Pro |
| 380 | MGCS36044_04116 | MGCS36089_04130 | ***ruvA*** |  |  | Holliday junction ATP-dependent DNA helicase RuvA | L | INFORMATION STORAGE AND PROCESSING; replication, recombination and repair | Core | Pro |
| 381 | MGCS36044_04128 | MGCS36089_04142 | ***argS*** |  |  | arginine--tRNA synthase ArgS | J | INFORMATION STORAGE AND PROCESSING; translation, ribosomal structure and biogenesis | Core | Pro |
| 382 | MGCS36044_04138 | MGCS36089_04152 | ***aspS*** |  |  | aspartyl-tRNA synthetase | J | INFORMATION STORAGE AND PROCESSING; translation, ribosomal structure and biogenesis | Core | Pro |
| 383 | MGCS36044_04140 | MGCS36089_04154 | ***hisS*** |  |  | histidine--tRNA synthase HisS | J | INFORMATION STORAGE AND PROCESSING; translation, ribosomal structure and biogenesis | Core | Pro |
| 384 | MGCS36044_04142 | MGCS36089_04156 | ***rpmF*** |  |  | 50S ribosomal L32 protein RpmF | J | INFORMATION STORAGE AND PROCESSING; translation, ribosomal structure and biogenesis | Core | Pro |
| 385 | MGCS36044_04200 | MGCS36089_04214 | ***-*** |  |  | IS1182 family transposase |  |  | Core | Pro |
| 386 | MGCS36044_04206 | MGCS36089_04220 | ***-*** |  |  | Veg family protein | R | General function prediction only | Core | Pro |
| 387 | MGCS36044_04208 | MGCS36089_04222 | ***dnaC*** |  |  | replicative DNA helicase DnaC | L | INFORMATION STORAGE AND PROCESSING; replication, recombination and repair | Core | Pro |
| 388 | MGCS36044_04210 | MGCS36089_04224 | ***rplI*** |  |  | 50S ribosomal L9 protein RplI | J | INFORMATION STORAGE AND PROCESSING; translation, ribosomal structure and biogenesis | Core | Pro |
| 389 | MGCS36044_04212 | MGCS36089_04226 | ***-*** |  |  | DHH family phosphoesterase | R | General function prediction only | Core | Pro |
| 390 | MGCS36044_04214 | MGCS36089_04228 | ***-*** |  |  | IS1548 family transposase | L | INFORMATION STORAGE AND PROCESSING; replication, recombination and repair | Core | Pro |
| 391 | MGCS36044_04216 | MGCS36089_04230 | ***mnmG*** |  |  | tRNA uridine-5-carboxymethylaminomethyl(34) synthesis enzyme MnmG | J | INFORMATION STORAGE AND PROCESSING; translation, ribosomal structure and biogenesis | Core | Pro |
| 392 | MGCS36044_04224 | MGCS36089_04238 | ***mnmA*** |  |  | tRNA 2-thiouridine(34) synthase MnmA | J | INFORMATION STORAGE AND PROCESSING; translation, ribosomal structure and biogenesis | Core | Pro |
| 393 | MGCS36044_04234 | MGCS36089_04248 | ***cbiQ_2*** |  |  | cobalt ABC transporter permease CbiQ | P | METABOLISM; inorganic ion transport and metabolism | Core | Pro |
| 394 | MGCS36044_04236 | MGCS36089_04250 | ***cbiO2*** |  |  | cobalt ABC transporter ATPase CbiO1 | P | METABOLISM; inorganic ion transport and metabolism | Core | Pro |
| 395 | MGCS36044_04238 | MGCS36089_04252 | ***cbiO1*** |  |  | cobalt ABC transporter ATPase CbiO2 | P | METABOLISM; inorganic ion transport and metabolism | Core | Pro |
| 396 | MGCS36044_04240 | MGCS36089_04254 | ***pgsA*** |  |  | CDP-diacylglycerol--glycerol-3-phosphate 3-phosphatidyltransferase PgsA | I | METABOLISM; lipid metabolism | Core | Pro |
| 397 | MGCS36044_04256 | MGCS36089_04270 | ***trpS*** |  |  | tryptophanyl-tRNA synthetase | J | INFORMATION STORAGE AND PROCESSING; translation, ribosomal structure and biogenesis | Core | Pro |
| 398 | MGCS36044_04268 | MGCS36089_04282 | ***-*** |  |  | disrupted IS30 family transposase encoding gene |  |  | Core | Pro |

† Genes are ordered ascendingly based on the locus tag numbers.

**^∫^** Refers to MGCS36044

**^∫∫^** Refers to MGCS36089

‡ Putative virulence genes (Eraso et al., 2024, *mBio*).

¶ As defined by the Prokka annotator (signal P.6); Secreted: secreted outside the cell, Lipo: membrane lipoprotein.

††Genomic location. Core: the core genome, Acc: the accessory genome, ROD: regions of difference.

‡‡ Pro: protein.
