## Supplementary material for "Gene contribution of *Streptococcus dysgalactiae* subspecies *equisimilis*, an emerging pathogen, to experimental primate necrotizing myositis": Table S3C

**Table S3C. Essential MGCS36089 genes during growth in *vivo* but not *in vitro***

| **No. ^†^** | **Locus tag** | **Gene** | **Exported^¶^** | **Function** | **COG class** | **COG description** | **Core/ Acc^††^** | **Pro/ RNA^‡‡^** |
| --- | --- | --- | --- | --- | --- | --- | --- | --- |
| 1 | MGCS36089_00768 | ***-*** |  | ABC amino acid transporter ATP-binding protein | E | METABOLISM; amino acid transport and metabolism | Core | Pro |
| 2 | MGCS36089_00780 | ***sufC*** |  | Fe-S cluster assembly ATPase SufC | H | METABOLISM; coenzyme transport and metabolism | Core | Pro |
| 3 | MGCS36089_00784 | ***sufS*** |  | cysteine desulfurase SufS | H | METABOLISM; coenzyme transport and metabolism | Core | Pro |
| 4 | MGCS36089_00788 | ***sufB*** |  | Fe-S cluster assembly protein SufB | H | METABOLISM; coenzyme transport and metabolism | Core | Pro |
| 5 | MGCS36089_00912 | ***scp2*** |  | segregation/condensation complex subunit (B) | D | CELLULAR PROCESSES AND SIGNALING; cell division and chromosome partitioning | Core | Pro |
| 6 | MGCS36089_01290 | ***lgt*** |  | prolipoprotein diacylglyceryl transferase Lgt | M | CELLULAR PROCESSES AND SIGNALING; cell envelope biogenesis, outermembrane | Core | Pro |
| 7 | MGCS36089_01370 | ***-*** |  | SSRC10. Infernal predicted noncoding RNA | | | Core | RNA |
| 8 | MGCS36089_01392 | ***rpiB*** |  | RpiB/LacA/LacB family sugar-phosphate isomerase | G | METABOLISM; carbohydrate transport and metabolism; Pentose-P pathway | Core | Pro |
| 9 | MGCS36089_01424 | ***aspC*** |  | aspartate aminotransferase protein AspC | E | METABOLISM; amino acid transport and metabolism | Core | Pro |
| 10 | MGCS36089_01434 | ***whiA*** |  | cell division involved DNA-binding protein WhiA | D | CELLULAR PROCESSES AND SIGNALING; cell division and chromosome partitioning | Core | Pro |
| 11 | MGCS36089_01556 | ***murA_1*** |  | UDP-N-acetylglucosamine 1-carboxyvinyltransferase protein MurA | M | CELLULAR PROCESSES AND SIGNALING; cell envelope biogenesis, outermembrane | Core | Pro |
| 12 | MGCS36089_01616 | ***-*** |  | glycosyltransferase family 1 protein | M | CELLULAR PROCESSES AND SIGNALING; cell envelope biogenesis, outermembrane | Core | Pro |
| 13 | MGCS36089_01620 | ***galE*** |  | UDP-glucose 4-epimerase GalE | M | CELLULAR PROCESSES AND SIGNALING; cell envelope biogenesis, outermembrane | Core | Pro |
| 14 | MGCS36089_01628 | ***-*** |  | LysM peptidoglycan-binding domain-containing | M | CELLULAR PROCESSES AND SIGNALING; cell envelope biogenesis, outermembrane | Core | Pro |
| 15 | MGCS36089_01642 | ***rlmK*** |  | 23S rRNA methyltransferase RmlK | J | INFORMATION STORAGE AND PROCESSING; translation, ribosomal structure and biogenesis | Core | Pro |
| 16 | MGCS36089_01712 | ***trxB_1*** |  | NAD(P)/FAD-dependent oxidoreductase | H | METABOLISM; coenzyme transport and metabolism | Core | Pro |
| 17 | MGCS36089_01900 | ***dacA_3*** | Secreted | secreted D,D-carboxypeptidase penicillin-binding. SignalP-6 predicted standard secretion signal | M | CELLULAR PROCESSES AND SIGNALING; cell envelope biogenesis, outermembrane | Core | Pro |
| 18 | MGCS36089_02002 | ***-*** |  | lysozyme family protein | M | CELLULAR PROCESSES AND SIGNALING; cell envelope biogenesis, outermembrane | Core | Pro |
| 19 | MGCS36089_02388 | ***-*** |  | DUF2969 domain-containing protein | S | POORLY CHARACTERIZED; function unknown | Core | Pro |
| 20 | MGCS36089_02838 | ***-*** |  | LCP family anionic cell polymer synthesis | M | CELLULAR PROCESSES AND SIGNALING; cell envelope biogenesis, outermembrane | Core | Pro |
| 21 | MGCS36089_02918 | ***-*** |  | LURP-one-related family protein | S | POORLY CHARACTERIZED; function unknown | Core | Pro |
| 22 | MGCS36089_02964 | ***deaD*** |  | DEAD/DEAH box helicase | L | INFORMATION STORAGE AND PROCESSING; Dna replication, recombination and repair | Core | Pro |
| 23 | MGCS36089_03024 | ***xseB*** |  | exodeoxyribonuclease VII small subunit XseB | L | INFORMATION STORAGE AND PROCESSING; Dna replication, recombination and repair | Core | Pro |
| 24 | MGCS36089_03330 | ***atoB*** |  | 3-ketoacyl-CoA thiolase/acetyl-CoA | I | METABOLISM; lipid metabolism | Core | Pro |

† Genes are ordered ascendingly based on the locus tag numbers.

¶ As defined by the Prokka annotator (signal P.6); Secreted: secreted outside the cell, Lipo: membrane lipoprotein.

††Genomic location. Core: the core genome, Acc: the accessory genome, ROD: regions of difference.

‡‡ Pro: protein.
