## Supplementary material for "Gene contribution of *Streptococcus dysgalactiae* subspecies *equisimilis*, an emerging pathogen, to experimental primate necrotizing myositis": Table S2C

**Table S2C. Essential MGCS36044 genes during growth in *vivo* but not *in vitro***

| **N0. ^†^** | **Locus tag** | **Gene** | **Exported^¶^** | **Function** | **COG class** | **COG description** | **Core/ Acc^††^** | **Pro/ RNA^‡‡^** |
| --- | --- | --- | --- | --- | --- | --- | --- | --- |
| 1 | MGCS36044_00360 | ***hslO*** |  | Hsp33 family molecular chaperone HslO | O | CELLULAR PROCESSES AND SIGNALING; post-translational modification, protein turnover, and chaperones | Core | Pro |
| 2 | MGCS36044_00390 | ***purA*** |  | adenylosuccinate synthase PurA | F | METABOLISM; nucleotide transport and metabolism | Core | Pro |
| 3 | MGCS36044_00728 | ***thiD*** |  | bifunctional hydroxymethylpyrimidine kinase/phosphomethylpyrimidine kinase PdxK | H | METABOLISM; coenzyme transport and metabolism | Core | Pro |
| 4 | MGCS36044_00774 | ***-*** | Secreted | ABC amino acid transporter substrate-binding secreted protein | P | METABOLISM; inorganic ion transport and metabolism | Core | Pro |
| 5 | MGCS36044_00784 | ***sufC*** |  | Fe-S cluster assembly ATPase SufC | H | METABOLISM; coenzyme transport and metabolism | Core | Pro |
| 6 | MGCS36044_00786 | ***sufD*** |  | Fe-S cluster assembly protein SufD | H | METABOLISM; coenzyme transport and metabolism | Core | Pro |
| 7 | MGCS36044_00790 | ***sufE*** |  | SUF system NifU family Fe-S cluster assembly protein SufE | H | METABOLISM; coenzyme transport and metabolism | Core | Pro |
| 8 | MGCS36044_00792 | ***sufB*** |  | Fe-S cluster assembly protein SufB | H | METABOLISM; coenzyme transport and metabolism | Core | Pro |
| 9 | MGCS36044_00868 | ***ktrB*** |  | potassium uptake transporter channel subunit KtrB | P | METABOLISM; inorganic ion transport and metabolism | Core | Pro |
| 10 | MGCS36044_00882 | ***nrdR*** |  | transcriptional regulator NrdR | K | INFORMATION STORAGE AND PROCESSING; transcription | Core | Pro |
| 11 | MGCS36044_01066 | ***mtsA*** | Lipo | metal ABC transporter substrate-binding lipoprotein MtsA | P | METABOLISM; inorganic ion transport and metabolism | Core | Pro |
| 12 | MGCS36044_01302 | ***lgt*** |  | prolipoprotein diacylglyceryl transferase Lgt | M | CELLULAR PROCESSES AND SIGNALING; cell wall/membrane/envelope biogenesis | Core | Pro |
| 13 | MGCS36044_01446 | ***whiA*** |  | cell division involved DNA-binding protein WhiA | D | CELLULAR PROCESSES AND SIGNALING; cell cycle control, cell division, chromosome partitioning | Core | Pro |
| 14 | MGCS36044_01568 | ***murA_1*** |  | UDP-N-acetylglucosamine 1-carboxyvinyltransferase protein MurA | M | CELLULAR PROCESSES AND SIGNALING; cell wall/membrane/envelope biogenesis | Core | Pro |
| 15 | MGCS36044_01622 | ***-*** |  | glycosyltransferase family 2 protein | M | CELLULAR PROCESSES AND SIGNALING; cell wall/membrane/envelope biogenesis | Core | Pro |
| 16 | MGCS36044_01632 | ***galE*** |  | UDP-glucose 4-epimerase GalE | M | CELLULAR PROCESSES AND SIGNALING; cell wall/membrane/envelope biogenesis | Core | Pro |
| 17 | MGCS36044_01724 | ***trxB_1*** |  | NAD(P)/FAD-dependent oxidoreductase | H | METABOLISM; coenzyme transport and metabolism | Core | Pro |
| 18 | MGCS36044_01742 | ***-*** |  | DUF1149 domain-containing protein | S | POORLY CHARACTERIZED; function unknown | Core | Pro |
| 19 | MGCS36044_01744 | ***degV_1*** |  | DegV family protein | S | POORLY CHARACTERIZED; function unknown | Core | Pro |
| 20 | MGCS36044_02020 | ***fadH2*** |  | FadH2 superfamily FAD-dependent oxidoreductase | E | METABOLISM; amino acid transport and metabolism | Core | Pro |
| 21 | MGCS36044_02128 | ***guaA*** |  | glutamine-hydrolyzing GMP synthase | F | METABOLISM; nucleotide transport and metabolism | ROD.5 | Pro |
| 22 | MGCS36044_02132 | ***mngR*** |  | MngR family DNA-binding transcriptional regulator | K | INFORMATION STORAGE AND PROCESSING; transcription | Core | Pro |
| 23 | MGCS36044_02382 | ***-*** |  | TVP38/TMEM64 family protein | M | CELLULAR PROCESSES AND SIGNALING; cell wall/membrane/envelope biogenesis | Core | Pro |
| 24 | MGCS36044_02670 | ***dltC*** |  | D-alanine--poly(phosphoribitol) ligase subunit DltC | M | CELLULAR PROCESSES AND SIGNALING; cell wall/membrane/envelope biogenesis | Core | Pro |
| 25 | MGCS36044_02908 | ***prsA*** | Lipo | peptidylprolyl isomerase lipoprotein PrsA | O | CELLULAR PROCESSES AND SIGNALING; post-translational modification, protein turnover, and chaperones | Core | Pro |
| 26 | MGCS36044_03860 | ***jag*** |  | RNA-binding protein Jag | L | INFORMATION STORAGE AND PROCESSING; replication, recombination and repair | Core | Pro |
| 27 | MGCS36044_04092 | ***-*** |  | DUF2079 domain-containing protein | S | POORLY CHARACTERIZED; function unknown | Core | Pro |
| 28 | MGCS36044_04184 | ***-*** |  | DUF368 domain-containing protein | S | POORLY CHARACTERIZED; function unknown | Core | Pro |

† Genes are ordered ascendingly based on the locus tag numbers.

¶ As defined by the Prokka annotator (signal P.6); Secreted: secreted outside the cell, Lipo: membrane lipoprotein.

††Genomic location. Core: the core genome, Acc: the accessory genome, ROD: regions of difference.

‡‡ Pro: protein.
