## Supplementary material for "Gene contribution of *Streptococcus dysgalactiae* subspecies *equisimilis*, an emerging pathogen, to experimental primate necrotizing myositis": Table S2B

**Table S2B. Essential MGCS36044 genes during growth *in vitro* but not in *vivo***

| **No. ^†^** | **Locus tag** | **Gene** | **Virulence^‡^** | **Function** | **COG class** | **COG description** | **Core/ Acc^††^** | **Pro/ RNA^‡‡^** |
| --- | --- | --- | --- | --- | --- | --- | --- | --- |
| 1 | MGCS36044_00006 | ***-*** |  | DUF951 doamin-containing protein | S | POORLY CHARACTERIZED; function unknown | Core | Pro |
| 2 | MGCS36044_00034 | ***-*** |  | IS30 family transposase |  |  | Core | Pro |
| 3 | MGCS36044_00112 | ***acpP_1*** |  | acyl carrier protein AcpP | I | METABOLISM; lipid metabolism | Core | Pro |
| 4 | MGCS36044_00150 | ***-*** |  | hypothetical protein | S | POORLY CHARACTERIZED; function unknown | Core | Pro |
| 5 | MGCS36044_00166 | ***rpsJ*** |  | 30S ribosomal S10 protein RpsJ | J | INFORMATION STORAGE AND PROCESSING; translation, ribosomal structure and biogenesis | Core | Pro |
| 6 | MGCS36044_00182 | ***rplP*** |  | 50S ribosomal L29 protein RplP | J | INFORMATION STORAGE AND PROCESSING; translation, ribosomal structure and biogenesis | Core | Pro |
| 7 | MGCS36044_00194 | ***rpsZ*** |  | type Z 30S ribosomal S14 protein RpsZ | J | INFORMATION STORAGE AND PROCESSING; translation, ribosomal structure and biogenesis | Core | Pro |
| 8 | MGCS36044_00214 | ***rpmJ*** |  | 50S ribosomal L36 protein RpmJ | J | INFORMATION STORAGE AND PROCESSING; translation, ribosomal structure and biogenesis | Core | Pro |
| 9 | MGCS36044_00468 | ***-*** |  | hypothetical protein | S | POORLY CHARACTERIZED; function unknown | ROD.2 | Pro |
| 10 | MGCS36044_00512 | ***nrdI_1*** |  | class Ib ribonucleoside-diphosphate reductase assembly flavoprotein NrdI | F | METABOLISM; nucleotide transport and metabolism | Core | Pro |
| 11 | MGCS36044_00658 | ***-*** |  | type II toxin-antitoxin system Phd/YefM family antitoxin | D | CELLULAR PROCESSES AND SIGNALING; cell division and chromosome partitioning | ROD.3 | Pro |
| 12 | MGCS36044_00880 | ***covS*** | Virulence | TCS sensor kinase CovS | T | CELLULAR PROCESSES AND SIGNALING; signal transduction mechanisms | Core | Pro |
| 13 | MGCS36044_01068 | ***mtsB*** |  | metal ABC transporter ATP-binding protein MtsB | P | METABOLISM; inorganic ion transport and metabolism | Core | Pro |
| 14 | MGCS36044_01074 | ***ftsK*** |  | cell division protein FtsK | D | CELLULAR PROCESSES AND SIGNALING; cell division and chromosome partitioning | Core | Pro |
| 15 | MGCS36044_01174 | ***secG*** |  | preprotein translocase subunit SecG | U | CELLULAR PROCESSES AND SIGNALING; Intracellular trafficking, secretion, and vesicular transport | Core | Pro |
| 16 | MGCS36044_01210 | ***dhaQ*** |  | DhaKLM operon coactivator DhaQ | G | METABOLISM; carbohydrate transport and metabolism | Core | Pro |
| 17 | MGCS36044_01334 | ***-*** |  | GH25 muramidase superfamily lysozyme | M | CELLULAR PROCESSES AND SIGNALING; cell wall/membrane/envelope biogenesis | Core | Pro |
| 18 | MGCS36044_01404 | ***rpiB*** |  | RpiB/LacA/LacB family sugar-phosphate isomerase | G | METABOLISM; carbohydrate transport and metabolism | Core | Pro |
| 19 | MGCS36044_01550 | ***atpE*** |  | ATP synthase C subunit AtpE | C | METABOLISM; energy production and conversion | Core | Pro |
| 20 | MGCS36044_01598 | ***rpsU*** |  | 30S ribosomal S21 protein RpsU | J | INFORMATION STORAGE AND PROCESSING; translation, ribosomal structure and biogenesis | Core | Pro |
| 21 | MGCS36044_01716 | ***-*** |  | KH domain-containing protein | L | INFORMATION STORAGE AND PROCESSING; replication, recombination and repair | Core | Pro |
| 22 | MGCS36044_02134 | ***ylxM*** |  | YlxM superfamily signal recognition particle associated DNA-binding protein | J | INFORMATION STORAGE AND PROCESSING; translation, ribosomal structure and biogenesis | Core | Pro |
| 23 | MGCS36044_02210 | ***-*** |  | DUF1836 domain-containing protein | S | POORLY CHARACTERIZED; function unknown | Core | Pro |
| 24 | MGCS36044_02318 | ***truB*** |  | tRNA pseudouridine(55) synthase TruB | J | INFORMATION STORAGE AND PROCESSING; translation, ribosomal structure and biogenesis | Core | Pro |
| 25 | MGCS36044_02336 | ***-*** |  | CsbD family protein | R | General function prediction only | Core | Pro |
| 26 | MGCS36044_02338 | ***-*** |  | Asp23/Gls24 family envelope stress response protein | M | CELLULAR PROCESSES AND SIGNALING; cell wall/membrane/envelope biogenesis | Core | Pro |
| 27 | MGCS36044_02350 | ***alsT*** |  | sodium:alanine symporter family protein | E | METABOLISM; amino acid transport and metabolism | Core | Pro |
| 28 | MGCS36044_02398 | ***-*** |  | DUF2969 domain-containing protein | S | POORLY CHARACTERIZED; function unknown | Core | Pro |
| 29 | MGCS36044_02438 | ***xapA*** |  | XapA family purine-nucleoside phosphorylase | F | METABOLISM; nucleotide transport and metabolism | Core | Pro |
| 30 | MGCS36044_02496 | ***lepA*** |  | translation elongation factor 4 LepA | J | INFORMATION STORAGE AND PROCESSING; translation, ribosomal structure and biogenesis | Core | Pro |
| 31 | MGCS36044_02546 | ***-*** |  | hypothetical protein | S | POORLY CHARACTERIZED; function unknown | Core | Pro |
| 32 | MGCS36044_02614 | ***apt*** |  | adenine phosphoribosyltransferase Apt | F | METABOLISM; nucleotide transport and metabolism | Core | Pro |
| 33 | MGCS36044_02626 | ***miaA*** |  | tRNA (adenosine(37)-N6)-dimethylallyltransferase MiaA | J | INFORMATION STORAGE AND PROCESSING; translation, ribosomal structure and biogenesis | Core | Pro |
| 34 | MGCS36044_02676 | ***dltX*** |  | teichoic acid D-Ala incorporation-associated protein DltX | M | CELLULAR PROCESSES AND SIGNALING; cell wall/membrane/envelope biogenesis | Core | Pro |
| 35 | MGCS36044_02706 | ***-*** |  | DUF4044 domain-containing protein | S | POORLY CHARACTERIZED; function unknown | Core | Pro |
| 36 | MGCS36044_02868 | ***birA*** |  | bifunctional biotin--[acetyl-CoA-carboxylase] ligase/biotin operon repressor BirA | K | INFORMATION STORAGE AND PROCESSING; transcription | Core | Pro |
| 37 | MGCS36044_03012 | ***xseB*** |  | exodeoxyribonuclease VII small subunit XseB | L | INFORMATION STORAGE AND PROCESSING; replication, recombination and repair | Core | Pro |
| 38 | MGCS36044_03052 | ***sepF*** |  | cell division protein SepF | D | CELLULAR PROCESSES AND SIGNALING; cell division and chromosome partitioning | Core | Pro |
| 39 | MGCS36044_03368 | ***ftsL*** |  | cell division protein FtsL | D | CELLULAR PROCESSES AND SIGNALING; cell division and chromosome partitioning | Core | Pro |
| 40 | MGCS36044_03422 | ***lacD_2*** |  | tagatose-bisphosphate aldolase LacD | G | METABOLISM; carbohydrate transport and metabolism | Core | Pro |
| 41 | MGCS36044_03462 | ***rbfA*** |  | 30S ribosome-binding factor RbfA | J | INFORMATION STORAGE AND PROCESSING; translation, ribosomal structure and biogenesis | Core | Pro |
| 42 | MGCS36044_03488 | ***-*** |  | Cps2a family anaionic cell wall polymer biosynthesis enxyme | K | INFORMATION STORAGE AND PROCESSING; transcription | Core | Pro |
| 43 | MGCS36044_03492 | ***tsaE*** |  | tRNA (adenosine(37)-N6)-threonylcarbamoyltransferase TsaE | J | INFORMATION STORAGE AND PROCESSING; translation, ribosomal structure and biogenesis | Core | Pro |
| 44 | MGCS36044_03526 | ***fabG_2*** |  | 3-ketoacyl-(acyl-carrier-protein) reductase protein FabG | I | METABOLISM; lipid metabolism | Core | Pro |
| 45 | MGCS36044_03532 | ***acpP_2*** |  | acyl carrier protein AcpP | I | METABOLISM; lipid metabolism | Core | Pro |
| 46 | MGCS36044_03534 | ***fabH*** |  | 3-oxoacyl-[acyl-carrier-protein] synthase protein FabH | I | METABOLISM; lipid metabolism | Core | Pro |
| 47 | MGCS36044_03624 | ***-*** |  | IS30 family transposase |  |  | Core | Pro |
| 48 | MGCS36044_03648 | ***-*** |  | CorA family divalent cation transport protein | P | METABOLISM; inorganic ion transport and metabolism | Core | Pro |
| 49 | MGCS36044_03798 | ***glnA*** |  | glutamine synthetase GlnA | E | METABOLISM; amino acid transport and metabolism | Core | Pro |
| 50 | MGCS36044_03910 | ***-*** |  | IS30 family transposase |  |  | Core | Pro |
| 51 | MGCS36044_03918 | ***-*** |  | disrupted IS30 family transposase encoding gene | | | Core | Pro |
| 52 | MGCS36044_03932 | ***tadA*** |  | tRNA adenosine(34) deaminase TadA | J | INFORMATION STORAGE AND PROCESSING; translation, ribosomal structure and biogenesis | Core | Pro |
| 53 | MGCS36044_03950 | ***polA*** |  | DNA polymerase I PolA | L | INFORMATION STORAGE AND PROCESSING; replication, recombination and repair | Core | Pro |
| 54 | MGCS36044_04178 | ***-*** |  | IS982 family transposase |  |  | ROD.9 | Pro |
| 55 | MGCS36044_04242 | ***rodZ*** |  | cytoskeltal protein RodZ | O | CELLULAR PROCESSES AND SIGNALING; post-translational modification, protein turnover, and chaperones | Core | Pro |
| 56 | MGCS36044_04250 | ***recF*** |  | DNA replication/repair protein RecF | L | INFORMATION STORAGE AND PROCESSING; replication, recombination and repair | Core | Pro |

† Genes are ordered ascendingly based on the locus tag numbers.

‡ Putative virulence genes (Eraso et al., 2024, *mBio*).

††Genomic location. Core: the core genome, Acc: the accessory genome, ROD: regions of difference.

‡‡ Pro: protein.
