## Supplementary material for "Gene contribution of *Streptococcus dysgalactiae* subspecies *equisimilis*, an emerging pathogen, to experimental primate necrotizing myositis": Table S2A

**Table S2A. Essential MGCS36044 genes shared during growth *in vivo* and *in vitro***

| **No.^†^** | **Locus tag** | **Gene** | **Virulence^‡^** | **Exported^¶^** | **Function** | **COG class** | | **COG description** | **Core/**  **Acc^††^** | **Pro/ RNA^‡‡^** |
| --- | --- | --- | --- | --- | --- | --- | --- | --- | --- | --- |
| 1 | MGCS36044_00002 | ***dnaA*** |  |  | chromosomal replication initiator protein DnaA | L | | INFORMATION STORAGE AND PROCESSING; replication, recombination and repair | Core | Pro |
| 2 | MGCS36044_00004 | ***dnaN*** |  |  | DNA polymerase III subunit beta protein DnaN | L | | INFORMATION STORAGE AND PROCESSING; replication, recombination and repair | Core | Pro |
| 3 | MGCS36044_00010 | ***engD*** |  |  | redox-regulated ATPase EngD | J | | INFORMATION STORAGE AND PROCESSING; translation, ribosomal structure and biogenesis | Core | Pro |
| 4 | MGCS36044_00012 | ***pth*** |  |  | aminoacyl-tRNA hydrolase Pth | J | | INFORMATION STORAGE AND PROCESSING; translation, ribosomal structure and biogenesis | Core | Pro |
| 5 | MGCS36044_00018 | ***-*** |  |  | RNA-binding S4 domain-containing protein | J | | INFORMATION STORAGE AND PROCESSING; translation, ribosomal structure and biogenesis | Core | Pro |
| 6 | MGCS36044_00020 | ***divIC*** |  |  | septum formation initiator family protein FtsB/DivIC | D | | CELLULAR PROCESSES AND SIGNALING; cell division and chromosome partitioning | Core | Pro |
| 7 | MGCS36044_00024 | ***-*** |  | Secreted | class A beta-lactamase-related serine hydrolase | M | | CELLULAR PROCESSES AND SIGNALING; cell envelope biogenesis, outermembrane | Core | Pro |
| 8 | MGCS36044_00026 | ***tilS*** |  |  | tRNA lysidine(34) synthetase TilS | J | | INFORMATION STORAGE AND PROCESSING; translation, ribosomal structure and biogenesis | Core | Pro |
| 9 | MGCS36044_00030 | ***ftsH*** |  |  | ATP-dependent zinc metalloprotease FtsH | O | | CELLULAR PROCESSES AND SIGNALING; post-translational modification, protein turnover, and chaperones | Core | Pro |
| 10 | MGCS36044_00104 | ***sibA*** |  | Secreted | CHAP domain-containing protein/secreted immunoglobulin-binding protein SibA | U | | CELLULAR PROCESSES AND SIGNALING; intracellular trafficking, secretion, and vesicular transport | Core | Pro |
| 11 | MGCS36044_00106 | ***prs*** |  |  | ribose-phosphate pyrophosphokinase PrsA | G | | METABOLISM; carbohydrate transport and metabolism | Core | Pro |
| 12 | MGCS36044_00110 | ***plsX*** |  |  | phosphate acyltransferase PlsX | I | | METABOLISM; lipid metabolism | Core | Pro |
| 13 | MGCS36044_00140 | ***purB*** |  |  | adenylosuccinate lyase PurB | F | | METABOLISM; nucleotide transport and metabolism | Core | Pro |
| 14 | MGCS36044_00146 | ***ruvB*** |  |  | Holliday junction branch migration DNA helicase RvuB | L | | INFORMATION STORAGE AND PROCESSING; replication, recombination and repair | Core | Pro |
| 15 | MGCS36044_00152 | ***oatA*** |  |  | acetyltransferase OatA | M | | CELLULAR PROCESSES AND SIGNALING; cell envelope biogenesis, outermembrane | Core | Pro |
| 16 | MGCS36044_00164 | ***-*** |  |  | IS30 family transposase | | |  | Core | Pro |
| 17 | MGCS36044_00168 | ***rplC*** |  |  | 50S ribosomal L3 protein RplC | J | | INFORMATION STORAGE AND PROCESSING; translation, ribosomal structure and biogenesis | Core | Pro |
| 18 | MGCS36044_00170 | ***rplD*** |  |  | 50S ribosomal L4 protein RplD | J | | INFORMATION STORAGE AND PROCESSING; translation, ribosomal structure and biogenesis | Core | Pro |
| 19 | MGCS36044_00172 | ***rplW*** |  |  | 50S ribosomal L23 protein RplW | J | | INFORMATION STORAGE AND PROCESSING; translation, ribosomal structure and biogenesis | Core | Pro |
| 20 | MGCS36044_00174 | ***rplB*** |  |  | 50S ribosomal L2 protein RplB | J | | INFORMATION STORAGE AND PROCESSING; translation, ribosomal structure and biogenesis | Core | Pro |
| 21 | MGCS36044_00176 | ***rpsS*** |  |  | 30S ribosomal S19 protein RpsS | J | | INFORMATION STORAGE AND PROCESSING; translation, ribosomal structure and biogenesis | Core | Pro |
| 22 | MGCS36044_00178 | ***rplV*** |  |  | 50S ribosomal L22 protein RplV | J | | INFORMATION STORAGE AND PROCESSING; translation, ribosomal structure and biogenesis | Core | Pro |
| 23 | MGCS36044_00180 | ***rpsC*** |  |  | 30S ribosomal S3 protein RpsC | J | | INFORMATION STORAGE AND PROCESSING; translation, ribosomal structure and biogenesis | Core | Pro |
| 24 | MGCS36044_00184 | ***rpmC*** |  |  | 50S ribosomal L16 protein RpmC | J | | INFORMATION STORAGE AND PROCESSING; translation, ribosomal structure and biogenesis | Core | Pro |
| 25 | MGCS36044_00186 | ***rpsQ*** |  |  | 30S ribosomal S17 protein RpsQ | J | | INFORMATION STORAGE AND PROCESSING; translation, ribosomal structure and biogenesis | Core | Pro |
| 26 | MGCS36044_00188 | ***rplN*** |  |  | 50S ribosomal L14 protein RplN | J | | INFORMATION STORAGE AND PROCESSING; translation, ribosomal structure and biogenesis | Core | Pro |
| 27 | MGCS36044_00190 | ***rplX*** |  |  | 50S ribosomal L24 protein RplX | J | | INFORMATION STORAGE AND PROCESSING; translation, ribosomal structure and biogenesis | Core | Pro |
| 28 | MGCS36044_00192 | ***rplE*** |  |  | 50S ribosomal L5 protein RplE | J | INFORMATION STORAGE AND PROCESSING; translation, ribosomal structure and biogenesis | | Core | Pro |
| 29 | MGCS36044_00196 | ***rpsH*** |  |  | 30S ribosomal S8 protein RpsH | J | INFORMATION STORAGE AND PROCESSING; translation, ribosomal structure and biogenesis | | Core | Pro |
| 30 | MGCS36044_00198 | ***rplF*** |  |  | 50S ribosomal L6 protein RplF | J | INFORMATION STORAGE AND PROCESSING; translation, ribosomal structure and biogenesis | | Core | Pro |
| 31 | MGCS36044_00200 | ***rplR*** |  |  | 50S ribosomal L18 protein RplR | J | INFORMATION STORAGE AND PROCESSING; translation, ribosomal structure and biogenesis | | Core | Pro |
| 32 | MGCS36044_00202 | ***rpsE*** |  |  | 30S ribosomal S5 protein RpsE | J | INFORMATION STORAGE AND PROCESSING; translation, ribosomal structure and biogenesis | | Core | Pro |
| 33 | MGCS36044_00204 | ***rpmD*** |  |  | 50S ribosomal L30 protein RpmD | J | INFORMATION STORAGE AND PROCESSING; translation, ribosomal structure and biogenesis | | Core | Pro |
| 34 | MGCS36044_00206 | ***rplO*** |  |  | 50S ribosomal L15 protein RplO | J | INFORMATION STORAGE AND PROCESSING; translation, ribosomal structure and biogenesis | | Core | Pro |
| 35 | MGCS36044_00208 | ***secY*** |  |  | preprotein translocase subunit SecY | U | CELLULAR PROCESSES AND SIGNALING; intracellular trafficking, secretion, and vesicular transport | | Core | Pro |
| 36 | MGCS36044_00210 | ***adk*** |  |  | adenylate kinase protein Adk | F | METABOLISM; nucleotide transport and metabolism | | Core | Pro |
| 37 | MGCS36044_00212 | ***infA*** |  |  | translation initiation factor IF-1 protein InfA | J | INFORMATION STORAGE AND PROCESSING; translation, ribosomal structure and biogenesis | | Core | Pro |
| 38 | MGCS36044_00216 | ***rpsM*** |  |  | 30S ribosomal S13 protein RpsM | J | INFORMATION STORAGE AND PROCESSING; translation, ribosomal structure and biogenesis | | Core | Pro |
| 39 | MGCS36044_00218 | ***rpsK*** |  |  | 30S ribosomal S11 protein RpsK | J | INFORMATION STORAGE AND PROCESSING; translation, ribosomal structure and biogenesis | | Core | Pro |
| 40 | MGCS36044_00220 | ***rpoA*** |  |  | DNA-directed RNA polymerase subunit alpha RpoA | K | INFORMATION STORAGE AND PROCESSING; transcription | | Core | Pro |
| 41 | MGCS36044_00222 | ***rplQ*** |  |  | 50S ribosomal L17 protein RplQ | J | INFORMATION STORAGE AND PROCESSING; translation, ribosomal structure and biogenesis | | Core | Pro |
| 42 | MGCS36044_00226 | ***-*** |  |  | disrupted IS1239 transposase encoding gene | | | | Core | Pro |
| 43 | MGCS36044_00228 | ***-*** |  |  | IS30 family transposase | |  | | Core | Pro |
| 44 | MGCS36044_00308 | ***tyrS*** |  |  | tyrosyl-tRNA synthetase TyrS | J | INFORMATION STORAGE AND PROCESSING; translation, ribosomal structure and biogenesis | | Core | Pro |
| 45 | MGCS36044_00310 | ***pbp1B*** |  |  | bifunctional PG transglycosylase-transpeptidase, penicillin-binding protein PBP1B | M | CELLULAR PROCESSES AND SIGNALING; cell envelope biogenesis, outermembrane | | Core | Pro |
| 46 | MGCS36044_00312 | ***-*** |  |  | Lacto-rpoB |  |  | | Core | RNA |
| 47 | MGCS36044_00314 | ***rpoB*** |  |  | DNA-directed RNA polymerase subunit beta RpoB | K | INFORMATION STORAGE AND PROCESSING; transcription | | Core | Pro |
| 48 | MGCS36044_00316 | ***rpoC*** |  |  | DNA-directed RNA polymerase subunit beta' RpoC | K | INFORMATION STORAGE AND PROCESSING; transcription | | Core | Pro |
| 49 | MGCS36044_00336 | ***ackA*** |  |  | acetate kinase AckA | C | METABOLISM; energy production and conversion | | Core | Pro |
| 50 | MGCS36044_00542 | ***uppS*** |  |  | UDP pyrophosphate synthase UppS | I | METABOLISM; lipid metabolism | | Core | Pro |
| 51 | MGCS36044_00544 | ***cdsA*** |  |  | phosphatidate cytidylyltransferase CdsA | I | METABOLISM; lipid metabolism | | Core | Pro |
| 52 | MGCS36044_00548 | ***proS*** |  |  | prolyl-tRNA synthetase ProS | J | INFORMATION STORAGE AND PROCESSING; translation, ribosomal structure and biogenesis | | Core | Pro |
| 53 | MGCS36044_00552 | ***polC*** |  |  | DNA polymerase III PolC | L | INFORMATION STORAGE AND PROCESSING; replication, recombination and repair | | Core | Pro |
| 54 | MGCS36044_00558 | ***def*** |  |  | peptide deformylase Def | J | INFORMATION STORAGE AND PROCESSING; translation, ribosomal structure and biogenesis | | Core | Pro |
| 55 | MGCS36044_00564 | ***rpsO*** |  |  | 30S ribosomal S15 protein RpsO | J | INFORMATION STORAGE AND PROCESSING; translation, ribosomal structure and biogenesis | | Core | Pro |
| 56 | MGCS36044_00574 | ***pnp*** |  |  | polyribonucleotide nucleotidyltransferase Pnp | J | INFORMATION STORAGE AND PROCESSING; translation, ribosomal structure and biogenesis | | Core | Pro |
| 57 | MGCS36044_00576 | ***-*** |  |  | polynucleotide phosphorylase/polyadenylase | L | INFORMATION STORAGE AND PROCESSING; replication, recombination and repair | | Core | Pro |
| 58 | MGCS36044_00578 | ***cysE*** |  |  | serine O-acetyltransferase CysE | E | METABOLISM; amino acid transport and metabolism | | Core | Pro |
| 59 | MGCS36044_00582 | ***cysS*** |  |  | cysteine--tRNA synthetase CysS | J | INFORMATION STORAGE AND PROCESSING; translation, ribosomal structure and biogenesis | | Core | Pro |
| 60 | MGCS36044_00600 | ***-*** |  |  | IS30 family transposase | |  | | Core | Pro |
| 61 | MGCS36044_00602 | ***-*** |  |  | L13_leader |  |  | | Core | RNA |
| 62 | MGCS36044_00604 | ***rplM*** |  |  | 50S ribosomal L13 protein RplM | J | INFORMATION STORAGE AND PROCESSING; translation, ribosomal structure and biogenesis | | Core | Pro |
| 63 | MGCS36044_00606 | ***rpsI*** |  |  | 30S ribosomal S9 protein RpsI | J | INFORMATION STORAGE AND PROCESSING; translation, ribosomal structure and biogenesis | | Core | Pro |
| 64 | MGCS36044_00610 | ***-*** |  |  | helix-turn-helix transcriptional regulator | K | INFORMATION STORAGE AND PROCESSING; transcription | | ROD.3 | Pro |
| 65 | MGCS36044_00622 | ***-*** |  |  | DNA cytosine methyltransferase | L | INFORMATION STORAGE AND PROCESSING; replication, recombination and repair | | ROD.3 | Pro |
| 66 | MGCS36044_00668 | ***-*** |  |  | Cro/CI family transcriptional regulator | K | INFORMATION STORAGE AND PROCESSING; transcription | | ROD.3 | Pro |
| 67 | MGCS36044_00708 | ***-*** |  |  | IS30 family transposase | |  | | Core | Pro |
| 68 | MGCS36044_00722 | ***comX_1*** |  |  | competence protein ComX | K | INFORMATION STORAGE AND PROCESSING; transcription | | Core | Pro |
| 69 | MGCS36044_00740 | ***tig*** |  |  | trigger factor molecular chaperone Tig | O | CELLULAR PROCESSES AND SIGNALING; post-translational modification, protein turnover, and chaperones | | Core | Pro |
| 70 | MGCS36044_00742 | ***rpoE*** |  |  | DNA-directed RNA polymerase subunit delta RpoE | K | INFORMATION STORAGE AND PROCESSING; transcription | | Core | Pro |
| 71 | MGCS36044_00758 | ***fba_2*** |  |  | fructose-bisphosphate aldolase | G | METABOLISM; carbohydrate transport and metabolism | | Core | Pro |
| 72 | MGCS36044_00760 | ***rpmB*** |  |  | 50S ribosomal L28 protein RpmB | J | INFORMATION STORAGE AND PROCESSING; translation, ribosomal structure and biogenesis | | Core | Pro |
| 73 | MGCS36044_00762 | ***-*** |  |  | IS1548 family transposase | L | INFORMATION STORAGE AND PROCESSING; replication, recombination and repair | | Core | Pro |
| 74 | MGCS36044_00764 | ***-*** |  |  | Asp23/Gls24 family envelope stress response protein | M | CELLULAR PROCESSES AND SIGNALING; cell envelope biogenesis, outermembrane | | Core | Pro |
| 75 | MGCS36044_00766 | ***-*** |  |  | DAK2 domain-containing protein | S | POORLY CHARACTERIZED; function unknown | | Core | Pro |
| 76 | MGCS36044_00780 | ***mecA*** |  |  | negative regulator of genetic competence, adaptor protein MecA | O | CELLULAR PROCESSES AND SIGNALING; post-translational modification, protein turnover, and chaperones | | Core | Pro |
| 77 | MGCS36044_00782 | ***rgpG*** |  |  | undecaprenyl/decaprenyl-phosphate alpha-N-acetylglucosaminyl 1-phosphate transferase RgpG | M | CELLULAR PROCESSES AND SIGNALING; cell envelope biogenesis, outermembrane | | Core | Pro |
| 78 | MGCS36044_00788 | ***sufS*** |  |  | cysteine desulfurase SufS | H | METABOLISM; coenzyme transport and metabolism | | Core | Pro |
| 79 | MGCS36044_00822 | ***comX_2*** |  |  | competence protein ComX | K | INFORMATION STORAGE AND PROCESSING; transcription | | Core | Pro |
| 80 | MGCS36044_00824 | ***yqeG*** |  |  | HAD IIIA-type phosphatase YqeG | F | METABOLISM; nucleotide transport and metabolism | | Core | Pro |
| 81 | MGCS36044_00826 | ***yqeH*** |  |  | ribosome biogenesis GTPase YqeH | P | METABOLISM; inorganic ion transport and metabolism | | Core | Pro |
| 82 | MGCS36044_00830 | ***nadD*** |  |  | nicotinate-nucleotide adenylyltransferase NadD | H | METABOLISM; coenzyme transport and metabolism | | Core | Pro |
| 83 | MGCS36044_00864 | ***sstT*** |  |  | serine/threonine transporter SstT | E | METABOLISM; amino acid transport and metabolism | | Core | Pro |
| 84 | MGCS36044_00876 | ***yceD*** |  |  | large ribosomal RNA subunit accumulation protein YceD | M | CELLULAR PROCESSES AND SIGNALING; cell envelope biogenesis, outermembrane | | Core | Pro |
| 85 | MGCS36044_00878 | ***covR*** | Virulence |  | TCS DNA-binding response regulator CovR | T | | CELLULAR PROCESSES AND SIGNALING; signal transduction mechanisms | Core | Pro |
| 86 | MGCS36044_00884 | ***dnaB*** |  |  | replication initiation and membrane attachment protein DnaB | L | | INFORMATION STORAGE AND PROCESSING; replication, recombination and repair | Core | Pro |
| 87 | MGCS36044_00886 | ***dnaI*** |  |  | primosomal protein DnaI | L | | INFORMATION STORAGE AND PROCESSING; replication, recombination and repair | Core | Pro |
| 88 | MGCS36044_00888 | ***der*** |  |  | ribosome biogenesis GTPase Der | J | | INFORMATION STORAGE AND PROCESSING; translation, ribosomal structure and biogenesis | Core | Pro |
| 89 | MGCS36044_00894 | ***murC*** |  |  | UDP-N-acetylmuramate--L-alanine ligase MurC | M | | CELLULAR PROCESSES AND SIGNALING; cell envelope biogenesis, outermembrane | Core | Pro |
| 90 | MGCS36044_00898 | ***mltG*** |  |  | endolytic transglycosylase MltG | F | | METABOLISM; nucleotide transport and metabolism | Core | Pro |
| 91 | MGCS36044_00900 | ***greA*** |  |  | transcription elongation factor GreA | K | | INFORMATION STORAGE AND PROCESSING; transcription | Core | Pro |
| 92 | MGCS36044_00902 | ***yidC_1*** |  | Lipo | membrane protein insertase lipoprotein YidC | M | | CELLULAR PROCESSES AND SIGNALING; cell envelope biogenesis, outermembrane | Core | Pro |
| 93 | MGCS36044_00912 | ***yneF*** |  |  | YneF family protein | S | | POORLY CHARACTERIZED; function unknown | Core | Pro |
| 94 | MGCS36044_00914 | ***murI*** |  |  | glutamate racemase MurI | M | | CELLULAR PROCESSES AND SIGNALING; cell envelope biogenesis, outermembrane | Core | Pro |
| 95 | MGCS36044_00938 | ***-*** |  |  | ECF transporter S component | I | | METABOLISM; lipid metabolism | Core | Pro |
| 96 | MGCS36044_00950 | ***ppaC*** |  |  | manganese-dependent inorganic pyrophosphatase PpaC | R | | General function prediction only | Core | Pro |
| 97 | MGCS36044_00962 | ***murE_1*** |  |  | UDP-N-acetylmuramoyl-L-alanyl-D-glutamate--L- lysine ligase MurE | M | | CELLULAR PROCESSES AND SIGNALING; cell envelope biogenesis, outermembrane | Core | Pro |
| 98 | MGCS36044_00964 | ***murJ*** |  |  | peptidoglycan lipid-II intermdiate flippase MurJ | O | | CELLULAR PROCESSES AND SIGNALING; post-translational modification, protein turnover, and chaperones | Core | Pro |
| 99 | MGCS36044_00968 | ***clpP*** |  |  | ATP-dependent Clp protease proteolytic subunit ClpP | O | | CELLULAR PROCESSES AND SIGNALING; post-translational modification, protein turnover, and chaperones | Core | Pro |
| 100 | MGCS36044_00988 | ***tmk*** |  |  | thymidylate kinase Tmk | F | | METABOLISM; nucleotide transport and metabolism | Core | Pro |
| 101 | MGCS36044_00990 | ***holB*** |  |  | DNA polymerase III subunit delta' HolB | L | | INFORMATION STORAGE AND PROCESSING; replication, recombination and repair | Core | Pro |
| 102 | MGCS36044_00994 | ***yabA*** |  |  | DNA replication initiation control protein YabA | L | | INFORMATION STORAGE AND PROCESSING; replication, recombination and repair | Core | Pro |
| 103 | MGCS36044_01028 | ***metS*** |  |  | methionine--tRNA synthase MetS | J | | INFORMATION STORAGE AND PROCESSING; translation, ribosomal structure and biogenesis | Core | Pro |
| 104 | MGCS36044_01054 | ***glmU*** |  |  | bifunctional UDP-N-acetylglucosamine diphosphorylase/glucosamine-1-phosphate N-acetyltransferase GlmU | G | | METABOLISM; carbohydrate transport and metabolism | Core | Pro |
| 105 | MGCS36044_01060 | ***mtnN*** |  |  | 5'-methylthioadenosine/adenosylhomocysteine nucleosidase MtnN | F | | METABOLISM; nucleotide transport and metabolism | Core | Pro |
| 106 | MGCS36044_01070 | ***mtsC*** |  |  | metal ABC transporter permease MtsC | P | | METABOLISM; inorganic ion transport and metabolism | Core | Pro |
| 107 | MGCS36044_01078 | ***rplK*** |  |  | 50S ribosomal L11P protein RplK | J | | INFORMATION STORAGE AND PROCESSING; translation, ribosomal structure and biogenesis | Core | Pro |
| 108 | MGCS36044_01080 | ***rplA*** |  |  | 50S ribosomal L1 protein RplA | J | | INFORMATION STORAGE AND PROCESSING; translation, ribosomal structure and biogenesis | Core | Pro |
| 109 | MGCS36044_01084 | ***pyrH*** |  |  | UMP kinase PyrH | F | | METABOLISM; nucleotide transport and metabolism | Core | Pro |
| 110 | MGCS36044_01086 | ***frr*** |  |  | ribosome recycling factor Frr | J | | INFORMATION STORAGE AND PROCESSING; translation, ribosomal structure and biogenesis | Core | Pro |
| 111 | MGCS36044_01102 | ***ybeY*** |  |  | rRNA maturation RNase YbeY | J | | INFORMATION STORAGE AND PROCESSING; translation, ribosomal structure and biogenesis | Core | Pro |
| 112 | MGCS36044_01104 | ***dgkA*** |  |  | diacylglycerol kinase DgkA | M | | CELLULAR PROCESSES AND SIGNALING; cell envelope biogenesis, outermembrane | Core | Pro |
| 113 | MGCS36044_01106 | ***era*** |  |  | GTPase Era | J | | INFORMATION STORAGE AND PROCESSING; translation, ribosomal structure and biogenesis | Core | Pro |
| 114 | MGCS36044_01122 | ***-*** |  |  | IS1182 family transposase | | |  | Core | Pro |
| 115 | MGCS36044_01124 | ***-*** |  |  | disrupted IS3 family transposase encoding gene | | | | Core | Pro |
| 116 | MGCS36044_01130 | ***-*** |  |  | IS1548 family transposase | L | | INFORMATION STORAGE AND PROCESSING; replication, recombination and repair | Core | Pro |
| 117 | MGCS36044_01164 | ***coaE*** |  |  | dephospho-CoA kinase CoaE | H | | METABOLISM; coenzyme transport and metabolism | Core | Pro |
| 118 | MGCS36044_01178 | ***smpB*** |  |  | SsrA(tmRNA)-binding protein SmpB | O | | CELLULAR PROCESSES AND SIGNALING; post-translational modification, protein turnover, and chaperones | Core | Pro |
| 119 | MGCS36044_01182 | ***-*** |  |  | disrupted IS3 family transposase encoding gene | | | | Core | Pro |
| 120 | MGCS36044_01194 | ***ccpA*** | Virulence |  | catabolite control protein CcpA | K | | INFORMATION STORAGE AND PROCESSING; transcription | Core | Pro |
| 121 | MGCS36044_01198 | ***-*** |  |  | glycosyltransferase | R | | General function prediction only | Core | Pro |
| 122 | MGCS36044_01200 | ***-*** |  |  | glycosyltransferase | M | | CELLULAR PROCESSES AND SIGNALING; cell envelope biogenesis, outermembrane | Core | Pro |
| 123 | MGCS36044_01202 | ***thrS*** |  |  | threonyl-tRNA synthetase ThrS | J | | INFORMATION STORAGE AND PROCESSING; translation, ribosomal structure and biogenesis | Core | Pro |
| 124 | MGCS36044_01238 | ***vicR*** | Virulence |  | TCS DNA-binding response regulator VicR | T | | CELLULAR PROCESSES AND SIGNALING; signal transduction mechanisms | Core | Pro |
| 125 | MGCS36044_01240 | ***vicK*** | Virulence |  | TCS signal transduction sensor kinase VicK | T | | CELLULAR PROCESSES AND SIGNALING; signal transduction mechanisms | Core | Pro |
| 126 | MGCS36044_01244 | ***rnc*** |  |  | ribonuclease III Rnc | A | | INFORMATION STORAGE AND PROCESSING; RNA processing and modification | Core | Pro |
| 127 | MGCS36044_01246 | ***smc*** |  |  | chromosome segregation protein Smc | D | | CELLULAR PROCESSES AND SIGNALING; cell division and chromosome partitioning | Core | Pro |
| 128 | MGCS36044_01250 | ***-*** |  |  | IS1548 family transposase | L | | INFORMATION STORAGE AND PROCESSING; replication, recombination and repair | Core | Pro |
| 129 | MGCS36044_01256 | ***ftsY*** |  |  | signal recognition particle-docking protein FtsY | U | | CELLULAR PROCESSES AND SIGNALING; intracellular trafficking, secretion, and vesicular transport | Core | Pro |
| 130 | MGCS36044_01300 | ***hprK*** |  |  | HPr(Ser) kinase/phosphatase HprK | T | | CELLULAR PROCESSES AND SIGNALING; signal transduction mechanisms | Core | Pro |
| 131 | MGCS36044_01308 | ***-*** |  |  | DUF3270 domain-containing protein | S | | POORLY CHARACTERIZED; function unknown | Core | Pro |
| 132 | MGCS36044_01320 | ***lysS*** |  |  | lysyl-tRNA synthetase LysS | J | | INFORMATION STORAGE AND PROCESSING; translation, ribosomal structure and biogenesis | Core | Pro |
| 133 | MGCS36044_01330 | ***thiT*** |  |  | energy-coupled thiamine transporter ThiT | H | | METABOLISM; coenzyme transport and metabolism | Core | Pro |
| 134 | MGCS36044_01336 | ***-*** |  |  | transposase IS116/IS110/IS902 family protein | | | | Core | Pro |
| 135 | MGCS36044_01348 | ***ftsW*** |  |  | cell division protein FtsW | D | | CELLULAR PROCESSES AND SIGNALING; cell division and chromosome partitioning | Core | Pro |
| 136 | MGCS36044_01354 | ***tufA*** |  |  | translation elongation factor Tu protein TufA | J | | INFORMATION STORAGE AND PROCESSING; translation, ribosomal structure and biogenesis | Core | Pro |
| 137 | MGCS36044_01364 | ***tpiA*** |  |  | triose-phosphate isomerase TpiA | G | | METABOLISM; carbohydrate transport and metabolism | Core | Pro |
| 138 | MGCS36044_01366 | ***-*** |  |  | disrupted IS3 family transposase encoding gene | | | | Core | Pro |
| 139 | MGCS36044_01368 | ***-*** |  |  | disrupted IS3 family transposase encoding gene | | | | Core | Pro |
| 140 | MGCS36044_01372 | ***murN*** |  |  | peptidoglycan lipid II-Ala--L-alanine ligase protein MurN | M | | CELLULAR PROCESSES AND SIGNALING; cell envelope biogenesis, outermembrane | Core | Pro |
| 141 | MGCS36044_01374 | ***murM*** |  |  | peptidoglycan lipid II--L-alanine ligase protein MurM | M | | CELLULAR PROCESSES AND SIGNALING; cell envelope biogenesis, outermembrane | Core | Pro |
| 142 | MGCS36044_01376 | ***-*** |  |  | sugar-phosphatase | R | | General function prediction only | Core | Pro |
| 143 | MGCS36044_01384 | ***mgtA*** |  |  | MgtA superfmily cation-translocating P-type ATPase | P | | METABOLISM; inorganic ion transport and metabolism | Core | Pro |
| 144 | MGCS36044_01426 | ***prfB*** |  |  | peptide chain release factor 2 PrfB | J | | INFORMATION STORAGE AND PROCESSING; translation, ribosomal structure and biogenesis | Core | Pro |
| 145 | MGCS36044_01428 | ***ftsE*** |  |  | cell division ATP-binding protein FtsE | D | | CELLULAR PROCESSES AND SIGNALING; cell division and chromosome partitioning | Core | Pro |
| 146 | MGCS36044_01430 | ***ftsX*** |  |  | cell division permease-like protein FtsX | D | | CELLULAR PROCESSES AND SIGNALING; cell division and chromosome partitioning | Core | Pro |
| 147 | MGCS36044_01436 | ***aspC*** |  |  | aspartate aminotransferase protein AspC | E | | METABOLISM; amino acid transport and metabolism | Core | Pro |
| 148 | MGCS36044_01438 | ***asnC*** |  |  | asparaginyl-tRNA synthetase protein AsnC | J | | INFORMATION STORAGE AND PROCESSING; translation, ribosomal structure and biogenesis | Core | Pro |
| 149 | MGCS36044_01456 | ***lacD_1*** |  |  | tagatose-bisphosphate aldolase LacD-like | G | | METABOLISM; carbohydrate transport and metabolism | Core | Pro |
| 150 | MGCS36044_01470 | ***rpmE*** |  |  | 50S ribosomal L31 type B protein RpmE | J | | INFORMATION STORAGE AND PROCESSING; translation, ribosomal structure and biogenesis | Core | Pro |
| 151 | MGCS36044_01472 | ***-*** |  |  | IS30 family transposase | | |  | Core | Pro |
| 152 | MGCS36044_01474 | ***nrnA*** |  |  | bifunctional oligoribonuclease/PAP phosphatase NrnA | J | | INFORMATION STORAGE AND PROCESSING; translation, ribosomal structure and biogenesis | Core | Pro |
| 153 | MGCS36044_01482 | ***-*** |  |  | IS1548 family transposase | L | | INFORMATION STORAGE AND PROCESSING; replication, recombination and repair | Core | Pro |
| 154 | MGCS36044_01484 | ***fldA*** |  |  | flavodoxin FldA | C | | METABOLISM; energy production and conversion | Core | Pro |
| 155 | MGCS36044_01490 | ***rplS*** |  |  | 50S ribosomal L19 protein RpsL | J | | INFORMATION STORAGE AND PROCESSING; translation, ribosomal structure and biogenesis | Core | Pro |
| 156 | MGCS36044_01496 | ***-*** |  |  | HAD-IA family hydrolase | R | | General function prediction only | Core | Pro |
| 157 | MGCS36044_01498 | ***gyrB*** |  |  | DNA topoisomerase ATP-hydrolyzing B subunit GyrB | L | | INFORMATION STORAGE AND PROCESSING; replication, recombination and repair | Core | Pro |
| 158 | MGCS36044_01500 | ***ezrA*** |  |  | cell division septation ring formation regulator EzrA | D | | CELLULAR PROCESSES AND SIGNALING; cell division and chromosome partitioning | Core | Pro |
| 159 | MGCS36044_01506 | ***eno*** |  |  | phosphopyruvate hydratase -- enolase protein Eno | G | | METABOLISM; carbohydrate transport and metabolism | Core | Pro |
| 160 | MGCS36044_01524 | ***sagG*** | Virulence |  | streptolysin S export protein SagG | U | | CELLULAR PROCESSES AND SIGNALING; intracellular trafficking, secretion, and vesicular transport | Core | Pro |
| 161 | MGCS36044_01526 | ***sagH*** | Virulence |  | streptolysin S export permease protein SagH | U | | CELLULAR PROCESSES AND SIGNALING; intracellular trafficking, secretion, and vesicular transport | Core | Pro |
| 162 | MGCS36044_01528 | ***sagI*** | Virulence |  | streptolysin S export permease protein SagI | U | | CELLULAR PROCESSES AND SIGNALING; intracellular trafficking, secretion, and vesicular transport | Core | Pro |
| 163 | MGCS36044_01536 | ***ligA*** |  |  | NAD-dependent DNA ligase LigA | L | | INFORMATION STORAGE AND PROCESSING; replication, recombination and repair | Core | Pro |
| 164 | MGCS36044_01538 | ***dagK*** |  |  | diacylglycerol kinase family lipid kinase | I | | METABOLISM; lipid metabolism | Core | Pro |
| 165 | MGCS36044_01552 | ***atpB*** |  |  | ATP synthase A subunit AtpB | C | | METABOLISM; energy production and conversion | Core | Pro |
| 166 | MGCS36044_01554 | ***atpF*** |  |  | ATP synthase B subunit AtpF | C | | METABOLISM; energy production and conversion | Core | Pro |
| 167 | MGCS36044_01556 | ***atpH*** |  |  | ATP synthase delta subunit AtpH | C | | METABOLISM; energy production and conversion | Core | Pro |
| 168 | MGCS36044_01558 | ***atpA*** |  |  | ATP synthase alpha chain, AtpA | C | | METABOLISM; energy production and conversion | Core | Pro |
| 169 | MGCS36044_01560 | ***atpG*** |  |  | ATP synthase gamma subunit AtpG | C | | METABOLISM; energy production and conversion | Core | Pro |
| 170 | MGCS36044_01562 | ***atpD*** |  |  | ATP synthase beta subunit AtpD | C | | METABOLISM; energy production and conversion | Core | Pro |
| 171 | MGCS36044_01564 | ***atpC*** |  |  | ATP synthase epsilon subunit AtpC | C | | METABOLISM; energy production and conversion | Core | Pro |
| 172 | MGCS36044_01574 | ***pheS*** |  |  | phenylalanyl-tRNA synthetase alpha subunit PheS | J | | INFORMATION STORAGE AND PROCESSING; translation, ribosomal structure and biogenesis | Core | Pro |
| 173 | MGCS36044_01576 | ***pheT*** |  |  | phenylalanyl-tRNA synthetase beta subunit PheT | J | | INFORMATION STORAGE AND PROCESSING; translation, ribosomal structure and biogenesis | Core | Pro |
| 174 | MGCS36044_01588 | ***rexB*** |  |  | ATP-dependent nuclease B subunit RexB | L | | INFORMATION STORAGE AND PROCESSING; replication, recombination and repair | Core | Pro |
| 175 | MGCS36044_01590 | ***rexA*** |  |  | ATP-dependent nuclease A subunit RexA | L | | INFORMATION STORAGE AND PROCESSING; replication, recombination and repair | Core | Pro |
| 176 | MGCS36044_01596 | ***-*** |  |  | IS1548 family transposase | L | | INFORMATION STORAGE AND PROCESSING; replication, recombination and repair | Core | Pro |
| 177 | MGCS36044_01602 | ***dnaG*** |  |  | DNA primase protein DnaG | L | | INFORMATION STORAGE AND PROCESSING; replication, recombination and repair | Core | Pro |
| 178 | MGCS36044_01604 | ***rpoD*** |  |  | RNA polymerase sigma factor RpoD | K | | INFORMATION STORAGE AND PROCESSING; transcription | Core | Pro |
| 179 | MGCS36044_01608 | ***rmlD*** |  |  | dTDP-4-dehydrorhamnose reductase protein RmlD | G | | METABOLISM; carbohydrate transport and metabolism | Core | Pro |
| 180 | MGCS36044_01610 | ***rgpA*** |  |  | glycosyltransferase family 1 protein RgpA | G | | METABOLISM; carbohydrate transport and metabolism | Core | Pro |
| 181 | MGCS36044_01612 | ***rgpB*** |  |  | glycosyltransferase family GT2 protein RgpB | G | | METABOLISM; carbohydrate transport and metabolism | Core | Pro |
| 182 | MGCS36044_01614 | ***rgpC*** |  |  | ABC transporter polysaccharide/polyol phosphate export permease RgpC | G | | METABOLISM; carbohydrate transport and metabolism | Core | Pro |
| 183 | MGCS36044_01616 | ***rgpD*** |  |  | ABC transporter polysaccharide/polyol phosphate ATPase component RgpD | G | | METABOLISM; carbohydrate transport and metabolism | Core | Pro |
| 184 | MGCS36044_01618 | ***rgpE*** |  |  | glycosyltransferase family GT2 protein RgpE | G | | METABOLISM; carbohydrate transport and metabolism | Core | Pro |
| 185 | MGCS36044_01620 | ***rgpF*** |  |  | alpha-L-Rha alpha-1,3-L-rhamnosyltransferase RgpF | G | | METABOLISM; carbohydrate transport and metabolism | Core | Pro |
| 186 | MGCS36044_01626 | ***-*** |  |  | DUF2142 domain-containing protein | S | | POORLY CHARACTERIZED; function unknown | Core | Pro |
| 187 | MGCS36044_01634 | ***-*** |  |  | RfbX superfamily lipopolysaccharide biosynthesis protein | M | | CELLULAR PROCESSES AND SIGNALING; cell envelope biogenesis, outermembrane | Core | Pro |
| 188 | MGCS36044_01642 | ***cmk*** |  |  | CMP kinase Cmk | F | | METABOLISM; nucleotide transport and metabolism | Core | Pro |
| 189 | MGCS36044_01644 | ***-*** |  |  | L20_leader |  | |  | Core | RNA |
| 190 | MGCS36044_01646 | ***infC*** |  |  | translation initiation factor InfC | J | | INFORMATION STORAGE AND PROCESSING; translation, ribosomal structure and biogenesis | Core | Pro |
| 191 | MGCS36044_01648 | ***rpmI*** |  |  | 50S ribosomal L35 protein RpmL | J | | INFORMATION STORAGE AND PROCESSING; translation, ribosomal structure and biogenesis | Core | Pro |
| 192 | MGCS36044_01650 | ***rplT*** |  |  | 50S ribosomal L20 protein RplT | J | | INFORMATION STORAGE AND PROCESSING; translation, ribosomal structure and biogenesis | Core | Pro |
| 193 | MGCS36044_01652 | ***ltaS*** |  |  | LTA synthase LtaS | M | | CELLULAR PROCESSES AND SIGNALING; cell envelope biogenesis, outermembrane | Core | Pro |
| 194 | MGCS36044_01654 | ***rlmK*** |  |  | 23S rRNA methyltransferase RmlK | J | | INFORMATION STORAGE AND PROCESSING; translation, ribosomal structure and biogenesis | Core | Pro |
| 195 | MGCS36044_01656 | ***aroD*** |  |  | type I 3-dehydroquinate dehydratase AroD | E | | METABOLISM; amino acid transport and metabolism | Core | Pro |
| 196 | MGCS36044_01674 | ***-*** |  |  | IS30 family transposase | | |  | Core | Pro |
| 197 | MGCS36044_01676 | ***-*** |  |  | L21_leader |  | |  | Core | RNA |
| 198 | MGCS36044_01678 | ***rplU*** |  |  | 50S ribosomal L21 protein RplU | J | | INFORMATION STORAGE AND PROCESSING; translation, ribosomal structure and biogenesis | Core | Pro |
| 199 | MGCS36044_01680 | ***prp*** |  |  | ribosomal-processing cysteine protease Prp | J | | INFORMATION STORAGE AND PROCESSING; translation, ribosomal structure and biogenesis | Core | Pro |
| 200 | MGCS36044_01682 | ***rpmA*** |  |  | 50S ribosomal L27 protein RpmA | J | | INFORMATION STORAGE AND PROCESSING; translation, ribosomal structure and biogenesis | Core | Pro |
| 201 | MGCS36044_01684 | ***lysR*** |  |  | LysR family transcriptional regulator | K | | INFORMATION STORAGE AND PROCESSING; transcription | Core | Pro |
| 202 | MGCS36044_01714 | ***rpsP*** |  |  | 30S ribosomal S16 protein RpsP | J | | INFORMATION STORAGE AND PROCESSING; translation, ribosomal structure and biogenesis | Core | Pro |
| 203 | MGCS36044_01720 | ***rimM*** |  |  | ribosome maturation factor RimM | J | | INFORMATION STORAGE AND PROCESSING; translation, ribosomal structure and biogenesis | Core | Pro |
| 204 | MGCS36044_01722 | ***trmD*** |  |  | tRNA (guanosine(37)-N1)-methyltransferase TrmD | J | | INFORMATION STORAGE AND PROCESSING; translation, ribosomal structure and biogenesis | Core | Pro |
| 205 | MGCS36044_01746 | ***-*** |  |  | tRNA CCA-pyrophosphorylase | J | | INFORMATION STORAGE AND PROCESSING; translation, ribosomal structure and biogenesis | Core | Pro |
| 206 | MGCS36044_01766 | ***mvaK1*** |  |  | mevalonate kinase MvaK1 | I | | METABOLISM; lipid metabolism | Core | Pro |
| 207 | MGCS36044_01768 | ***mvaD*** |  |  | diphosphomevalonate decarboxylase MvaD | I | | METABOLISM; lipid metabolism | Core | Pro |
| 208 | MGCS36044_01770 | ***mvaK2*** |  |  | mevalonate kinase MvaK2 | I | | METABOLISM; lipid metabolism | Core | Pro |
| 209 | MGCS36044_01772 | ***-*** |  |  | isopentenyl-diphosphate delta-isomerase | I | | METABOLISM; lipid metabolism | Core | Pro |
| 210 | MGCS36044_01782 | ***mvaS1*** |  |  | hydroxymethylglutaryl-CoA reductase protein (1) MvaS1 | I | | METABOLISM; lipid metabolism | Core | Pro |
| 211 | MGCS36044_01784 | ***mvaS2*** |  |  | hydroxymethylglutaryl-CoA synthase protein (2) MvaS2 | I | | METABOLISM; lipid metabolism | Core | Pro |
| 212 | MGCS36044_01786 | ***thyA*** |  |  | thymidylate synthase ThyA | F | | METABOLISM; nucleotide transport and metabolism | Core | Pro |
| 213 | MGCS36044_01788 | ***dyr*** |  |  | dihydrofolate reductase Dyr | H | | METABOLISM; coenzyme transport and metabolism | Core | Pro |
| 214 | MGCS36044_01792 | ***clpX*** |  |  | ATP-dependent Clp protease, ATP-binding subunit ClpX | O | | CELLULAR PROCESSES AND SIGNALING; post-translational modification, protein turnover, and chaperones | Core | Pro |
| 215 | MGCS36044_01794 | ***engB*** |  |  | ribosome biogenesis GTP-binding protein EngB | J | | INFORMATION STORAGE AND PROCESSING; translation, ribosomal structure and biogenesis | Core | Pro |
| 216 | MGCS36044_01800 | ***-*** |  |  | L10_leader |  | |  | Core | RNA |
| 217 | MGCS36044_01802 | ***rplJ*** |  |  | 50S ribosomal L10 protein RplJ | J | | INFORMATION STORAGE AND PROCESSING; translation, ribosomal structure and biogenesis | Core | Pro |
| 218 | MGCS36044_01804 | ***rplL*** |  |  | 50S ribosomal L7/L12 protein RplL | J | | INFORMATION STORAGE AND PROCESSING; translation, ribosomal structure and biogenesis | Core | Pro |
| 219 | MGCS36044_01806 | ***-*** |  |  | rli38 |  | |  | Core | RNA |
| 220 | MGCS36044_01838 | ***-*** |  |  | type IV toxin-antitoxin system AbiEi family antitoxin | V | | CELLULAR PROCESSES AND SIGNALING; defense mechanisms | ROD.4 | Pro |
| 221 | MGCS36044_01920 | ***folC*** |  |  | dihydrofolate synthase FolC | H | | METABOLISM; coenzyme transport and metabolism | Core | Pro |
| 222 | MGCS36044_01922 | ***folE*** |  |  | GTP cyclohydrolase I protein FolE | H | | METABOLISM; coenzyme transport and metabolism | Core | Pro |
| 223 | MGCS36044_01924 | ***folP*** |  |  | dihydropteroate synthase protein FolP | H | | METABOLISM; coenzyme transport and metabolism | Core | Pro |
| 224 | MGCS36044_01926 | ***folQ*** |  |  | dihydroneopterin aldolase protein FolB | H | | METABOLISM; coenzyme transport and metabolism | Core | Pro |
| 225 | MGCS36044_01928 | ***folK*** |  |  | 2-amino-4-hydroxy-6- hydroxymethyldihydropteridine pyrophosphokinase protein FolK | H | | METABOLISM; coenzyme transport and metabolism | Core | Pro |
| 226 | MGCS36044_01930 | ***murB*** |  |  | UDP-N-acetylmuramate dehydrogenase MurB | M | | CELLULAR PROCESSES AND SIGNALING; cell envelope biogenesis, outermembrane | Core | Pro |
| 227 | MGCS36044_01960 | ***rex*** |  |  | redox-sensing transcriptional repressor Rex | K | | INFORMATION STORAGE AND PROCESSING; transcription | Core | Pro |
| 228 | MGCS36044_01966 | ***nifS_2*** |  |  | NifS superfamily cysteine desulfurase | E | | METABOLISM; amino acid transport and metabolism | Core | Pro |
| 229 | MGCS36044_01968 | ***ribP*** |  |  | ribose-phosphate pyrophosphokinase RibP | F | | METABOLISM; nucleotide transport and metabolism | Core | Pro |
| 230 | MGCS36044_01974 | ***nadK*** |  |  | NAD kinase NadK | H | | METABOLISM; coenzyme transport and metabolism | Core | Pro |
| 231 | MGCS36044_01978 | ***eutD*** |  |  | phosphate acetyltransferase EutD | C | | METABOLISM; energy production and conversion | Core | Pro |
| 232 | MGCS36044_01982 | ***-*** |  |  | Na+ driven multidrug efflux pump | P | | METABOLISM; inorganic ion transport and metabolism | Core | Pro |
| 233 | MGCS36044_02002 | ***prfA*** |  |  | peptide chain release factor 1 PrfA | J | | INFORMATION STORAGE AND PROCESSING; translation, ribosomal structure and biogenesis | Core | Pro |
| 234 | MGCS36044_02004 | ***prmC*** |  |  | peptide chain release factor N(5)-glutamine methyltransferase PrmC | J | | INFORMATION STORAGE AND PROCESSING; translation, ribosomal structure and biogenesis | Core | Pro |
| 235 | MGCS36044_02006 | ***-*** |  |  | Sua5/YciO/YrdC/YwlC family protein ribosome maturation factor | J | | INFORMATION STORAGE AND PROCESSING; translation, ribosomal structure and biogenesis | Core | Pro |
| 236 | MGCS36044_02024 | ***ldh*** |  |  | L-lactate dehydrogenase Ldh | C | | METABOLISM; energy production and conversion | Core | Pro |
| 237 | MGCS36044_02026 | ***gyrA*** |  |  | DNA gyrase subunit A GyrA | L | | INFORMATION STORAGE AND PROCESSING; replication, recombination and repair | Core | Pro |
| 238 | MGCS36044_02028 | ***srtA*** |  |  | class A sortase SrtA | M | | CELLULAR PROCESSES AND SIGNALING; cell envelope biogenesis, outermembrane | Core | Pro |
| 239 | MGCS36044_02038 | ***-*** |  |  | disrupted IS3 family transposase encoding gene | | | | Core | Pro |
| 240 | MGCS36044_02078 | ***-*** |  |  | disrupted IS3 family transposase encoding gene | | | | Core | Pro |
| 241 | MGCS36044_02080 | ***-*** |  |  | IS30 family transposase | | |  | Core | Pro |
| 242 | MGCS36044_02086 | ***-*** |  |  | disrupted IS1182 family transposase encoding gene | | | | Core | Pro |
| 243 | MGCS36044_02136 | ***ffh*** |  |  | signal recognition particle protein | U | | CELLULAR PROCESSES AND SIGNALING; intracellular trafficking, secretion, and vesicular transport | Core | Pro |
| 244 | MGCS36044_02154 | ***xerS*** |  |  | site-specific tyrosine recombinase XerS | L | | INFORMATION STORAGE AND PROCESSING; replication, recombination and repair | ROD.6 | Pro |
| 245 | MGCS36044_02200 | ***topA*** |  |  | type I DNA topoisomerase TopA | L | | INFORMATION STORAGE AND PROCESSING; replication, recombination and repair | Core | Pro |
| 246 | MGCS36044_02206 | ***ylqF*** |  |  | ribosome biogenesis GTPase YlqF | J | | INFORMATION STORAGE AND PROCESSING; translation, ribosomal structure and biogenesis | Core | Pro |
| 247 | MGCS36044_02212 | ***yqfA*** |  |  | membrane channel forming/hemolysin III protein YqfA | K | | INFORMATION STORAGE AND PROCESSING; transcription | Core | Pro |
| 248 | MGCS36044_02242 | ***-*** |  |  | disrupted IS3 family transposase encoding gene | | | | Core | Pro |
| 249 | MGCS36044_02262 | ***coaB*** |  |  | phosphopantothenate--cysteine ligase CoaB | H | | METABOLISM; coenzyme transport and metabolism | Core | Pro |
| 250 | MGCS36044_02264 | ***coaC*** |  |  | phosphopantothenoylcysteine decarboxylase CoaC | H | | METABOLISM; coenzyme transport and metabolism | Core | Pro |
| 251 | MGCS36044_02266 | ***panT*** |  |  | pantothenic acid transporter PanT | T | | CELLULAR PROCESSES AND SIGNALING; signal transduction mechanisms | Core | Pro |
| 252 | MGCS36044_02268 | ***pgmA*** |  |  | phospho-sugar mutase PgmA | G | | METABOLISM; carbohydrate transport and metabolism | Core | Pro |
| 253 | MGCS36044_02286 | ***coaA*** |  |  | type I pantothenate kinase | H | | METABOLISM; coenzyme transport and metabolism | Core | Pro |
| 254 | MGCS36044_02296 | ***phoU_2*** |  |  | phosphate signaling complex protein PhoU | P | | METABOLISM; inorganic ion transport and metabolism | Core | Pro |
| 255 | MGCS36044_02298 | ***ptsB1*** |  |  | phosphate ABC transporter ATP-binding protein PstB1 | P | | METABOLISM; inorganic ion transport and metabolism | Core | Pro |
| 256 | MGCS36044_02300 | ***ptsB2*** |  |  | phosphate ABC transporter ATP-binding protein PstB2 | P | | METABOLISM; inorganic ion transport and metabolism | Core | Pro |
| 257 | MGCS36044_02314 | ***spxA_1*** |  |  | transcriptional regulator SpxA | K | | INFORMATION STORAGE AND PROCESSING; transcription | Core | Pro |
| 258 | MGCS36044_02316 | ***ribF*** |  |  | bifunctional riboflavin kinase/FAD synthetase RibF | H | | METABOLISM; coenzyme transport and metabolism | Core | Pro |
| 259 | MGCS36044_02332 | ***-*** |  |  | GntR family transcriptional regulator | K | | INFORMATION STORAGE AND PROCESSING; transcription | Core | Pro |
| 260 | MGCS36044_02346 | ***pcrA*** |  |  | DNA helicase PcrA | L | | INFORMATION STORAGE AND PROCESSING; replication, recombination and repair | Core | Pro |
| 261 | MGCS36044_02348 | ***-*** |  |  | glycine |  | |  | Core | RNA |
| 262 | MGCS36044_02354 | ***-*** |  |  | IS1182 family transposase | | |  | Core | Pro |
| 263 | MGCS36044_02364 | ***glmS*** |  |  | glutamine--fructose-6-phosphate transaminase (isomerizing) | E | | METABOLISM; amino acid transport and metabolism | Core | Pro |
| 264 | MGCS36044_02370 | ***pyk*** |  |  | pyruvate kinase Pyk | G | | METABOLISM; carbohydrate transport and metabolism | Core | Pro |
| 265 | MGCS36044_02372 | ***pfkA*** |  |  | 6-phosphofructokinase PfkA | G | | METABOLISM; carbohydrate transport and metabolism | Core | Pro |
| 266 | MGCS36044_02374 | ***dnaE*** |  |  | DNA polymerase III subunit alpha DnaE | L | | INFORMATION STORAGE AND PROCESSING; replication, recombination and repair | Core | Pro |
| 267 | MGCS36044_02376 | ***yhcF*** |  |  | YhcF family transcriptional regulator | K | | INFORMATION STORAGE AND PROCESSING; transcription | Core | Pro |
| 268 | MGCS36044_02384 | ***ssrA*** |  |  | transfer-messenger RNA | | |  | Core | RNA |
| 269 | MGCS36044_02388 | ***-*** |  |  | IS1182 family transposase | | |  | Core | Pro |
| 270 | MGCS36044_02392 | ***rpsA*** |  |  | 30S ribosomal S1 protein RpsA | J | | INFORMATION STORAGE AND PROCESSING; translation, ribosomal structure and biogenesis | Core | Pro |
| 271 | MGCS36044_02402 | ***parC*** |  |  | DNA topoisomerase IV subunit A ParC | L | | INFORMATION STORAGE AND PROCESSING; replication, recombination and repair | Core | Pro |
| 272 | MGCS36044_02406 | ***parE*** |  |  | DNA topoisomerase IV subunit B ParE | L | | INFORMATION STORAGE AND PROCESSING; replication, recombination and repair | Core | Pro |
| 273 | MGCS36044_02408 | ***plsY*** |  |  | glycerol-3-phosphate 1-O-acyltransferase PlsY | L | | INFORMATION STORAGE AND PROCESSING; replication, recombination and repair | Core | Pro |
| 274 | MGCS36044_02444 | ***rpiA*** |  |  | ribose-5-phosphate isomerase RpiA | G | | METABOLISM; carbohydrate transport and metabolism | Core | Pro |
| 275 | MGCS36044_02446 | ***mnmE*** |  |  | MnmE family tRNA uridine-5-carboxymethylaminomethyl(34) synthesis GTPase | J | | INFORMATION STORAGE AND PROCESSING; translation, ribosomal structure and biogenesis | Core | Pro |
| 276 | MGCS36044_02448 | ***pepV*** |  |  | dipeptidase PepV | E | | METABOLISM; amino acid transport and metabolism | Core | Pro |
| 277 | MGCS36044_02528 | ***glmM*** |  |  | phosphoglucosamine mutase GlmM | G | | METABOLISM; carbohydrate transport and metabolism | Core | Pro |
| 278 | MGCS36044_02532 | ***-*** |  |  | DisA N domain-containing diadenylate cyclase | G | | METABOLISM; carbohydrate transport and metabolism | Core | Pro |
| 279 | MGCS36044_02534 | ***murE_2*** |  |  | UDP-N-acetylmuramoylalanyl-D-glutamate-2, 6-diaminopimelate ligase MurE | M | | CELLULAR PROCESSES AND SIGNALING; cell envelope biogenesis, outermembrane | Core | Pro |
| 280 | MGCS36044_02536 | ***-*** |  |  | CobQ-like type 1 glutamine amidotransferase | M | | CELLULAR PROCESSES AND SIGNALING; cell envelope biogenesis, outermembrane | Core | Pro |
| 281 | MGCS36044_02538 | ***lplA_2*** |  |  | lipoate--protein ligase LplA | H | | METABOLISM; coenzyme transport and metabolism | Core | Pro |
| 282 | MGCS36044_02544 | ***acoL*** |  |  | dihydrolipoyl dehydrogenase AcoL | C | | METABOLISM; energy production and conversion | Core | Pro |
| 283 | MGCS36044_02548 | ***acoC*** |  |  | dihydrolipoamide acetyltransferase AcoC | C | | METABOLISM; energy production and conversion | Core | Pro |
| 284 | MGCS36044_02550 | ***acoB*** |  |  | pyruvate dehydrogenase E1 component beta subunit AcoB | C | | METABOLISM; energy production and conversion | Core | Pro |
| 285 | MGCS36044_02552 | ***acoA*** |  |  | Pyruvate dehydrogenase E1 component alpha subunit AcoA | C | | METABOLISM; energy production and conversion | Core | Pro |
| 286 | MGCS36044_02560 | ***rnjA_1*** |  |  | mRNA degradation ribonuclease RnjA | A | | INFORMATION STORAGE AND PROCESSING; RNA processing and modification | Core | Pro |
| 287 | MGCS36044_02578 | ***-*** |  |  | disrupted IS3 family transposase encoding gene | | | | Core | Pro |
| 288 | MGCS36044_02596 | ***rfbB*** |  |  | dTDP-glucose 4,6-dehydratase RfbB | M | | CELLULAR PROCESSES AND SIGNALING; cell envelope biogenesis, outermembrane | Core | Pro |
| 289 | MGCS36044_02598 | ***rfbC*** |  |  | dTDP-4-dehydrorhamnose 3,5-epimerase RfbC | G | | METABOLISM; carbohydrate transport and metabolism | Core | Pro |
| 290 | MGCS36044_02600 | ***rfbA*** |  |  | glucose-1-phosphate thymidylyltransferase RfbA | M | | CELLULAR PROCESSES AND SIGNALING; cell envelope biogenesis, outermembrane | Core | Pro |
| 291 | MGCS36044_02608 | ***trmK*** |  |  | tRNA (adenine(22)-N(1))-methyltransferase TrmK | L | | INFORMATION STORAGE AND PROCESSING; replication, recombination and repair | Core | Pro |
| 292 | MGCS36044_02612 | ***dnaD*** |  |  | DNA replication protein DnaD | L | | INFORMATION STORAGE AND PROCESSING; replication, recombination and repair | Core | Pro |
| 293 | MGCS36044_02620 | ***rnz*** |  |  | ribonuclease Rnz | J | | INFORMATION STORAGE AND PROCESSING; translation, ribosomal structure and biogenesis | Core | Pro |
| 294 | MGCS36044_02644 | ***malQ*** |  |  | 4-alpha-glucanotransferase (amylomaltase) protein MalQ | G | | METABOLISM; carbohydrate transport and metabolism | Core | Pro |
| 295 | MGCS36044_02648 | ***-*** |  |  | IS30 family transposase | | |  | Core | Pro |
| 296 | MGCS36044_02668 | ***dltD*** |  |  | D-alanyl-lipoteichoic acid biosynthesis protein DltD | M | | CELLULAR PROCESSES AND SIGNALING; cell envelope biogenesis, outermembrane | Core | Pro |
| 297 | MGCS36044_02672 | ***dltB*** |  |  | D-alanyl-lipoteichoic acid biosynthesis protein DltB | M | | CELLULAR PROCESSES AND SIGNALING; cell envelope biogenesis, outermembrane | Core | Pro |
| 298 | MGCS36044_02674 | ***dltA*** |  |  | D-alanine--poly(phosphoribitol) ligase subunit DltA | M | | CELLULAR PROCESSES AND SIGNALING; cell envelope biogenesis, outermembrane | Core | Pro |
| 299 | MGCS36044_02704 | ***obgE*** |  |  | GTPase ObgE | L | | INFORMATION STORAGE AND PROCESSING; replication, recombination and repair | Core | Pro |
| 300 | MGCS36044_02730 | ***-*** |  |  | disrupted IS30 family transposase encoding gene | | | | Core | Pro |
| 301 | MGCS36044_02834 | ***map*** |  |  | methionyl aminopeptidase Map | J | | INFORMATION STORAGE AND PROCESSING; translation, ribosomal structure and biogenesis | Core | Pro |
| 302 | MGCS36044_02864 | ***metK*** |  |  | methionine adenosyltransferase MetK | J | | INFORMATION STORAGE AND PROCESSING; translation, ribosomal structure and biogenesis | Core | Pro |
| 303 | MGCS36044_02870 | ***dnaX*** |  |  | DNA polymerase III gamma/tau subunit DnaX | L | | INFORMATION STORAGE AND PROCESSING; replication, recombination and repair | Core | Pro |
| 304 | MGCS36044_02880 | ***srmB*** |  |  | SrmB superfailily II DNA and RNA helicase | L | | INFORMATION STORAGE AND PROCESSING; replication, recombination and repair | Core | Pro |
| 305 | MGCS36044_02884 | ***gapN*** |  |  | NADP-dependent glyceraldehyde-3-phosphate dehydrogenase GapN | G | | METABOLISM; carbohydrate transport and metabolism | Core | Pro |
| 306 | MGCS36044_02886 | ***ptsI*** |  |  | phosphoenolpyruvate--protein phosphotransferase PtsI | G | | METABOLISM; carbohydrate transport and metabolism | Core | Pro |
| 307 | MGCS36044_02888 | ***ptsH*** |  |  | PTS transporter phosphocarrier protein PtsH | G | | METABOLISM; carbohydrate transport and metabolism | Core | Pro |
| 308 | MGCS36044_02890 | ***nrdH*** |  |  | glutaredoxin-like protein NrdH | O | | CELLULAR PROCESSES AND SIGNALING; post-translational modification, protein turnover, and chaperones | Core | Pro |
| 309 | MGCS36044_02892 | ***nrdE_2*** |  |  | class 1b ribonucleoside-diphosphate reductase alpha subunit NrdE | F | | METABOLISM; nucleotide transport and metabolism | Core | Pro |
| 310 | MGCS36044_02894 | ***nrdF_2*** |  |  | class 1b ribonucleoside-diphosphate reductase beta subunit NrdF | F | | METABOLISM; nucleotide transport and metabolism | Core | Pro |
| 311 | MGCS36044_02904 | ***alaS*** |  |  | alanine--tRNA synthetase AlaS | J | | INFORMATION STORAGE AND PROCESSING; translation, ribosomal structure and biogenesis | Core | Pro |
| 312 | MGCS36044_02936 | ***sodA*** |  |  | superoxide dismutase SodA | V | | CELLULAR PROCESSES AND SIGNALING; defense mechanisms | Core | Pro |
| 313 | MGCS36044_02938 | ***holA*** |  |  | DNA polymerase III delta subunit HolA | L | | INFORMATION STORAGE AND PROCESSING; replication, recombination and repair | Core | Pro |
| 314 | MGCS36044_02944 | ***plsC*** |  | Secreted | secreted 1-acyl-sn-glycerol-3-phosphate acyltransferase PlsC | I | | METABOLISM; lipid metabolism | Core | Pro |
| 315 | MGCS36044_02952 | ***deaD*** |  |  | DEAD/DEAH box helicase | L | | INFORMATION STORAGE AND PROCESSING; replication, recombination and repair | Core | Pro |
| 316 | MGCS36044_02966 | ***murF*** |  |  | UDP-N-acetylmuramoyl-tripeptide--D-alanyl-D- alanine ligase MurF | M | | CELLULAR PROCESSES AND SIGNALING; cell envelope biogenesis, outermembrane | Core | Pro |
| 317 | MGCS36044_02968 | ***ddl*** |  |  | D-alanine--D-alanine ligase Ddl | M | | CELLULAR PROCESSES AND SIGNALING; cell envelope biogenesis, outermembrane | Core | Pro |
| 318 | MGCS36044_02970 | ***recR*** |  |  | recombination mediator RecR | L | | INFORMATION STORAGE AND PROCESSING; replication, recombination and repair | Core | Pro |
| 319 | MGCS36044_02980 | ***gpmA*** |  |  | phosphoglycerate mutase GpmA | G | | METABOLISM; carbohydrate transport and metabolism | Core | Pro |
| 320 | MGCS36044_02982 | ***-*** |  |  | IS982 family transposase | | |  | Core | Pro |
| 321 | MGCS36044_02994 | ***-*** |  |  | DNA-binding protein HU | L | | INFORMATION STORAGE AND PROCESSING; replication, recombination and repair | Core | Pro |
| 322 | MGCS36044_03008 | ***tlyA*** |  |  | TlyA family RNA methyltransferase | J | | INFORMATION STORAGE AND PROCESSING; translation, ribosomal structure and biogenesis | Core | Pro |
| 323 | MGCS36044_03014 | ***xseA*** |  |  | exodeoxyribonuclease VII large subunit XseA | L | | INFORMATION STORAGE AND PROCESSING; replication, recombination and repair | Core | Pro |
| 324 | MGCS36044_03018 | ***folD*** |  |  | bifunctional methylenetetrahydrofolate dehydrogenase/methenyltetrahydrofolate cyclohydrolase FolD | E | | METABOLISM; amino acid transport and metabolism | Core | Pro |
| 325 | MGCS36044_03044 | ***ileS*** |  |  | isoleucine--tRNA synthetase IleS | J | | INFORMATION STORAGE AND PROCESSING; translation, ribosomal structure and biogenesis | Core | Pro |
| 326 | MGCS36044_03046 | ***divIVA*** |  |  | cell division protein DivIVA | D | | CELLULAR PROCESSES AND SIGNALING; cell division and chromosome partitioning | Core | Pro |
| 327 | MGCS36044_03048 | ***-*** |  |  | RNA-binding protein | R | | General function prediction only | Core | Pro |
| 328 | MGCS36044_03050 | ***-*** |  |  | YggT family protein | S | | POORLY CHARACTERIZED; function unknown | Core | Pro |
| 329 | MGCS36044_03054 | ***yggS*** |  |  | YggS family pyridoxal phosphate-dependent enzyme | F | | METABOLISM; nucleotide transport and metabolism | Core | Pro |
| 330 | MGCS36044_03056 | ***ftsZ*** |  |  | cell division protein FtsZ | D | | CELLULAR PROCESSES AND SIGNALING; cell division and chromosome partitioning | Core | Pro |
| 331 | MGCS36044_03058 | ***ftsA*** |  |  | cell division protein FtsA | D | | CELLULAR PROCESSES AND SIGNALING; cell division and chromosome partitioning | Core | Pro |
| 332 | MGCS36044_03060 | ***ftsQ*** |  |  | cell division protein FtsQ/DivIB | D | | CELLULAR PROCESSES AND SIGNALING; cell division and chromosome partitioning | Core | Pro |
| 333 | MGCS36044_03062 | ***murG*** |  |  | UDP-N-acetylglucosamine--N-acetylmuramyl- (pentapeptide) pyrophosphoryl-undecaprenol N-acetylglucosamine transferase MurG | M | | CELLULAR PROCESSES AND SIGNALING; cell envelope biogenesis, outermembrane | Core | Pro |
| 334 | MGCS36044_03064 | ***murD*** |  |  | UDP-N-acetylmuramoyl-L-alanine--D-glutamate ligase MurD | M | | CELLULAR PROCESSES AND SIGNALING; cell envelope biogenesis, outermembrane | Core | Pro |
| 335 | MGCS36044_03070 | ***typA*** |  |  | translational GTPase TypA | T | | CELLULAR PROCESSES AND SIGNALING; signal transduction mechanisms | Core | Pro |
| 336 | MGCS36044_03092 | ***-*** |  |  | IS1548 family transposase | L | | INFORMATION STORAGE AND PROCESSING; replication, recombination and repair | Core | Pro |
| 337 | MGCS36044_03096 | ***-*** |  |  | IS110 family transposase | | |  | Core | Pro |
| 338 | MGCS36044_03112 | ***coaD*** |  |  | pantetheine-phosphate adenylyltransferase CoaD | H | | METABOLISM; coenzyme transport and metabolism | Core | Pro |
| 339 | MGCS36044_03120 | ***-*** |  |  | IS110 family transposase | | |  | Core | Pro |
| 340 | MGCS36044_03170 | ***valS*** |  |  | valine--tRNA synthetase ValS | J | | INFORMATION STORAGE AND PROCESSING; translation, ribosomal structure and biogenesis | Core | Pro |
| 341 | MGCS36044_03180 | ***-*** |  |  | DUF1912 family protein | S | | POORLY CHARACTERIZED; function unknown | Core | Pro |
| 342 | MGCS36044_03278 | ***-*** |  |  | IS30 family transposase | | |  | Core | Pro |
| 343 | MGCS36044_03288 | ***cysK*** |  |  | cysteine synthase A CysK | E | | METABOLISM; amino acid transport and metabolism | Core | Pro |
| 344 | MGCS36044_03294 | ***liaR*** | Virulence |  | three component system signal transduction response regulator protein LiaR | K | | INFORMATION STORAGE AND PROCESSING; transcription | Core | Pro |
| 345 | MGCS36044_03300 | ***pknB*** |  |  | Stk1 family PASTA domain-containing Ser/Thr kinase | T | | CELLULAR PROCESSES AND SIGNALING; signal transduction mechanisms | Core | Pro |
| 346 | MGCS36044_03302 | ***pppL*** |  |  | Stp1/IreP family PP2C-type Ser/Thr phosphatase | T | | CELLULAR PROCESSES AND SIGNALING; signal transduction mechanisms | Core | Pro |
| 347 | MGCS36044_03306 | ***fmt*** |  |  | methionyl-tRNA formyl transferase Fmt | J | | INFORMATION STORAGE AND PROCESSING; translation, ribosomal structure and biogenesis | Core | Pro |
| 348 | MGCS36044_03308 | ***priA*** |  |  | primosomal protein PriA | L | | INFORMATION STORAGE AND PROCESSING; replication, recombination and repair | Core | Pro |
| 349 | MGCS36044_03310 | ***rpoZ*** |  |  | DNA-directed RNA polymerase omega subunit RpoZ | K | | INFORMATION STORAGE AND PROCESSING; transcription | Core | Pro |
| 350 | MGCS36044_03312 | ***gmk*** |  |  | guanylate kinase Gmk | F | | METABOLISM; nucleotide transport and metabolism | Core | Pro |
| 351 | MGCS36044_03314 | ***rny*** |  |  | ribonuclease (Y) Rny | D | | CELLULAR PROCESSES AND SIGNALING; cell division and chromosome partitioning | Core | Pro |
| 352 | MGCS36044_03330 | ***mapZ*** |  |  | MapZ family cell division site-positioning protein | D | | CELLULAR PROCESSES AND SIGNALING; cell division and chromosome partitioning | Core | Pro |
| 353 | MGCS36044_03334 | ***-*** |  |  | RNaseP_bact_b |  | |  | Core | RNA |
| 354 | MGCS36044_03336 | ***gpsB*** |  |  | cell division regulator GpsB | D | | CELLULAR PROCESSES AND SIGNALING; cell division and chromosome partitioning | Core | Pro |
| 355 | MGCS36044_03340 | ***recU*** |  |  | Holliday junction resolvase RecU | L | | INFORMATION STORAGE AND PROCESSING; replication, recombination and repair | Core | Pro |
| 356 | MGCS36044_03342 | ***pbp1A*** |  |  | bifunctional PG transglycosylase-transpeptidase classA penicillin-binding protein PBP1A | M | | CELLULAR PROCESSES AND SIGNALING; cell envelope biogenesis, outermembrane | Core | Pro |
| 357 | MGCS36044_03346 | ***nadE*** |  |  | ammonia-dependent NAD(+) synthetase NadE | H | | METABOLISM; coenzyme transport and metabolism | Core | Pro |
| 358 | MGCS36044_03348 | ***pncB*** |  |  | nicotinate phosphoribosyltransferase PncB | H | | METABOLISM; coenzyme transport and metabolism | Core | Pro |
| 359 | MGCS36044_03352 | ***trxB_2*** |  |  | thioredoxin-disulfide reductase TrxB | C | | METABOLISM; energy production and conversion | Core | Pro |
| 360 | MGCS36044_03354 | ***-*** |  |  | DUF4059 family protein | S | | POORLY CHARACTERIZED; function unknown | Core | Pro |
| 361 | MGCS36044_03356 | ***-*** |  |  | GlnQ family polar amino acid ABC transporter ATPase | E | | METABOLISM; amino acid transport and metabolism | Core | Pro |
| 362 | MGCS36044_03358 | ***hisM*** |  |  | HisM family amino acid ABC transporter permease | P | | METABOLISM; inorganic ion transport and metabolism | Core | Pro |
| 363 | MGCS36044_03362 | ***cshB*** |  |  | DEAD/DEAH box helicase | L | | INFORMATION STORAGE AND PROCESSING; replication, recombination and repair | Core | Pro |
| 364 | MGCS36044_03364 | ***mraY*** |  |  | phospho-N-acetylmuramoyl-pentapeptide- translocase MraY | M | | CELLULAR PROCESSES AND SIGNALING; cell envelope biogenesis, outermembrane | Core | Pro |
| 365 | MGCS36044_03366 | ***pbp2X*** |  |  | PG transpeptidase class B penicillin-binding protein PBP2X/FtsI | M | | CELLULAR PROCESSES AND SIGNALING; cell envelope biogenesis, outermembrane | Core | Pro |
| 366 | MGCS36044_03370 | ***mraW*** |  |  | S-adenosyl-methyltransferase MraW | D | | CELLULAR PROCESSES AND SIGNALING; cell division and chromosome partitioning | Core | Pro |
| 367 | MGCS36044_03380 | ***-*** |  |  | disrupted IS3 family transposase encoding gene | | | | Core | Pro |
| 368 | MGCS36044_03382 | ***tkt*** |  |  | transketolase Tkt | G | | METABOLISM; carbohydrate transport and metabolism | Core | Pro |
| 369 | MGCS36044_03398 | ***ynzC*** |  |  | DUF896 family protein | S | | POORLY CHARACTERIZED; function unknown | Core | Pro |
| 370 | MGCS36044_03400 | ***glyS*** |  |  | glycine--tRNA ligase beta subunit GlyS | J | | INFORMATION STORAGE AND PROCESSING; translation, ribosomal structure and biogenesis | Core | Pro |
| 371 | MGCS36044_03402 | ***glyQ*** |  |  | glycine--tRNA ligase alpha subunit GlyQ | J | | INFORMATION STORAGE AND PROCESSING; translation, ribosomal structure and biogenesis | Core | Pro |
| 372 | MGCS36044_03414 | ***degV_2*** |  |  | DegV family fatty acid-binding protein | I | | METABOLISM; lipid metabolism | Core | Pro |
| 373 | MGCS36044_03464 | ***infB*** |  |  | translation initiation factor IF-2 | J | | INFORMATION STORAGE AND PROCESSING; translation, ribosomal structure and biogenesis | Core | Pro |
| 374 | MGCS36044_03466 | ***-*** |  |  | YlxQ-related RNA-binding protein | J | | INFORMATION STORAGE AND PROCESSING; translation, ribosomal structure and biogenesis | Core | Pro |
| 375 | MGCS36044_03468 | ***-*** |  |  | YlxR family putative RNA-binding protein | K | | INFORMATION STORAGE AND PROCESSING; transcription | Core | Pro |
| 376 | MGCS36044_03470 | ***nusA*** |  |  | transcription termination factor NusA | K | | INFORMATION STORAGE AND PROCESSING; transcription | Core | Pro |
| 377 | MGCS36044_03472 | ***rimP*** |  |  | ribosome maturation factor RimP | J | | INFORMATION STORAGE AND PROCESSING; translation, ribosomal structure and biogenesis | Core | Pro |
| 378 | MGCS36044_03478 | ***cotS*** |  |  | CotS family thaimine kinase | M | | CELLULAR PROCESSES AND SIGNALING; cell envelope biogenesis, outermembrane | Core | Pro |
| 379 | MGCS36044_03512 | ***serS*** |  |  | seryl-tRNA synthetase SerS | J | | INFORMATION STORAGE AND PROCESSING; translation, ribosomal structure and biogenesis | Core | Pro |
| 380 | MGCS36044_03514 | ***accD*** |  |  | acetyl-CoA carboxylase carboxyl transferase alpha subunit AccD | I | | METABOLISM; lipid metabolism | Core | Pro |
| 381 | MGCS36044_03516 | ***accA*** |  |  | acetyl-CoA carboxylase, carboxyltransferase beta subunit AccA | I | | METABOLISM; lipid metabolism | Core | Pro |
| 382 | MGCS36044_03518 | ***accC*** |  |  | acetyl-CoA carboxylase biotin carboxylase subunit AccC | I | | METABOLISM; lipid metabolism | Core | Pro |
| 383 | MGCS36044_03520 | ***fabZ*** |  |  | 3-hydroxyacyl-ACP dehydratase FabZ | I | | METABOLISM; lipid metabolism | Core | Pro |
| 384 | MGCS36044_03522 | ***accB*** |  |  | acetyl-CoA carboxylase biotin carboxyl carrier protein AccB | I | | METABOLISM; lipid metabolism | Core | Pro |
| 385 | MGCS36044_03524 | ***fabF*** |  |  | 3-oxoacyl-[acyl-carrier-protein] synthase protein FabF | I | | METABOLISM; lipid metabolism | Core | Pro |
| 386 | MGCS36044_03528 | ***fabD*** |  |  | malonyl CoA-acyl carrier protein transacylase protein FabD | I | | METABOLISM; lipid metabolism | Core | Pro |
| 387 | MGCS36044_03530 | ***fabK*** |  |  | Enoyl-[acyl-carrier-protein] reductase protein FabK | I | | METABOLISM; lipid metabolism | Core | Pro |
| 388 | MGCS36044_03536 | ***fabT*** |  |  | transcriptional regulatory protein FabT | I | | METABOLISM; lipid metabolism | Core | Pro |
| 389 | MGCS36044_03538 | ***phaB*** |  |  | enoyl-CoA hydratase protein PhaB | I | | METABOLISM; lipid metabolism | Core | Pro |
| 390 | MGCS36044_03540 | ***dnaJ*** |  |  | chaperone protein DnaJ | O | | CELLULAR PROCESSES AND SIGNALING; post-translational modification, protein turnover, and chaperones | Core | Pro |
| 391 | MGCS36044_03544 | ***dnaK*** |  |  | molecular chaperone DnaK | O | | CELLULAR PROCESSES AND SIGNALING; post-translational modification, protein turnover, and chaperones | Core | Pro |
| 392 | MGCS36044_03546 | ***grpE*** |  |  | heat shock protein/nucleotide exchange factor GrpE | O | | CELLULAR PROCESSES AND SIGNALING; post-translational modification, protein turnover, and chaperones | Core | Pro |
| 393 | MGCS36044_03548 | ***hrcA*** |  |  | heat-inducible transcriptional repressor HrcA | K | | INFORMATION STORAGE AND PROCESSING; transcription | Core | Pro |
| 394 | MGCS36044_03560 | ***gatB_2*** |  |  | aspartyl-tRNA(Asn) or glutamyl-tRNA(Gln) amidotransferase B subunit GatB | J | | INFORMATION STORAGE AND PROCESSING; translation, ribosomal structure and biogenesis | Core | Pro |
| 395 | MGCS36044_03562 | ***gatA_2*** |  |  | aspartyl-tRNA(Asn) or glutamyl-tRNA(Gln) amidotransferase A subunit GatA | J | | INFORMATION STORAGE AND PROCESSING; translation, ribosomal structure and biogenesis | Core | Pro |
| 396 | MGCS36044_03564 | ***gatC_2*** |  |  | aspartyl-tRNA(Asn) or glutamyl-tRNA(Gln) amidotransferase C subunit GatC | J | | INFORMATION STORAGE AND PROCESSING; translation, ribosomal structure and biogenesis | Core | Pro |
| 397 | MGCS36044_03582 | ***codY*** |  |  | CodY family GTP-sensing pleiotropic transcriptional regulator | K | | INFORMATION STORAGE AND PROCESSING; transcription | Core | Pro |
| 398 | MGCS36044_03594 | ***recG*** |  |  | ATP-dependent DNA helicase RecG | L | | INFORMATION STORAGE AND PROCESSING; replication, recombination and repair | Core | Pro |
| 399 | MGCS36044_03618 | ***alr*** |  |  | alanine racemase Alr | E | | METABOLISM; amino acid transport and metabolism | Core | Pro |
| 400 | MGCS36044_03620 | ***acpS*** |  |  | AcpS family provisional 4'-phosphopantetheinyl transferase | I | | METABOLISM; lipid metabolism | Core | Pro |
| 401 | MGCS36044_03622 | ***secA*** |  |  | preprotein translocase subunit SecA | U | | CELLULAR PROCESSES AND SIGNALING; intracellular trafficking, secretion, and vesicular transport | Core | Pro |
| 402 | MGCS36044_03632 | ***scrB*** |  |  | sucrose-6-phosphate hydrolase ScrB | G | | METABOLISM; carbohydrate transport and metabolism | Core | Pro |
| 403 | MGCS36044_03636 | ***nusB*** |  |  | transcription termination protein NusB | K | | INFORMATION STORAGE AND PROCESSING; transcription | Core | Pro |
| 404 | MGCS36044_03638 | ***-*** |  |  | Asp23/Gls24 family envelope stress response protein | R | | General function prediction only | Core | Pro |
| 405 | MGCS36044_03640 | ***efp*** |  |  | translation elongation factor (P) Efp | J | | INFORMATION STORAGE AND PROCESSING; translation, ribosomal structure and biogenesis | Core | Pro |
| 406 | MGCS36044_03652 | ***rpsR*** |  |  | 30S ribosomal S18 protein RpsR | J | | INFORMATION STORAGE AND PROCESSING; translation, ribosomal structure and biogenesis | Core | Pro |
| 407 | MGCS36044_03654 | ***ssb_2*** |  |  | single-stranded DNA-binding protein | L | | INFORMATION STORAGE AND PROCESSING; replication, recombination and repair | Core | Pro |
| 408 | MGCS36044_03656 | ***rpsF*** |  |  | 30S ribosomal S6 protein RpsF | J | | INFORMATION STORAGE AND PROCESSING; translation, ribosomal structure and biogenesis | Core | Pro |
| 409 | MGCS36044_03662 | ***trxA_2*** |  |  | thioredoxin TrxA | O | | CELLULAR PROCESSES AND SIGNALING; post-translational modification, protein turnover, and chaperones | Core | Pro |
| 410 | MGCS36044_03668 | ***-*** |  |  | colicin V production family protein | O | | CELLULAR PROCESSES AND SIGNALING; post-translational modification, protein turnover, and chaperones | Core | Pro |
| 411 | MGCS36044_03672 | ***rnhC*** |  |  | HIII ribonuclease RnhC | L | | INFORMATION STORAGE AND PROCESSING; replication, recombination and repair | Core | Pro |
| 412 | MGCS36044_03674 | ***lepB_2*** |  |  | signal peptidase I LepB | U | | CELLULAR PROCESSES AND SIGNALING; intracellular trafficking, secretion, and vesicular transport | Core | Pro |
| 413 | MGCS36044_03704 | ***glpF_2*** |  |  | glycerol uptake facilitator GlpF | G | | METABOLISM; carbohydrate transport and metabolism | Core | Pro |
| 414 | MGCS36044_03738 | ***-*** |  |  | disrupted IS3 family transposase encoding gene | | | | ROD.8 | Pro |
| 415 | MGCS36044_03770 | ***-*** |  |  | DUF536 domain-containing protein | S | | POORLY CHARACTERIZED; function unknown | Core | Pro |
| 416 | MGCS36044_03778 | ***tsaD*** |  |  | tRNA (adenosine(37)-N6)-threonylcarbamoyltransferase complex transferase subunit TsaD | J | | INFORMATION STORAGE AND PROCESSING; translation, ribosomal structure and biogenesis | Core | Pro |
| 417 | MGCS36044_03782 | ***tsaB*** |  |  | tRNA (adenosine(37)-N6)-threonylcarbamoyltransferase complex dimerization subunit type 1 TsaB | O | | CELLULAR PROCESSES AND SIGNALING; post-translational modification, protein turnover, and chaperones | Core | Pro |
| 418 | MGCS36044_03784 | ***-*** |  |  | DUF1447 family protein | S | | POORLY CHARACTERIZED; function unknown | Core | Pro |
| 419 | MGCS36044_03786 | ***rnjA_2*** |  |  | mRNA degradation ribonuclease RnjA | A | | INFORMATION STORAGE AND PROCESSING; RNA processing and modification | Core | Pro |
| 420 | MGCS36044_03804 | ***pgk*** |  |  | phosphoglycerate kinase Pgk | G | | METABOLISM; carbohydrate transport and metabolism | Core | Pro |
| 421 | MGCS36044_03808 | ***gapA*** |  |  | glyceraldehyde-3-phosphate dehydrogenase GapA | G | | METABOLISM; carbohydrate transport and metabolism | Core | Pro |
| 422 | MGCS36044_03810 | ***fusA*** |  |  | FusA family elongation factor EF-G | J | | INFORMATION STORAGE AND PROCESSING; translation, ribosomal structure and biogenesis | Core | Pro |
| 423 | MGCS36044_03812 | ***rpsG*** |  |  | 30S ribosomal S7 protein RpsG | J | | INFORMATION STORAGE AND PROCESSING; translation, ribosomal structure and biogenesis | Core | Pro |
| 424 | MGCS36044_03814 | ***rpsL*** |  |  | 30S ribosomal S12 protein RpsL | J | | INFORMATION STORAGE AND PROCESSING; translation, ribosomal structure and biogenesis | Core | Pro |
| 425 | MGCS36044_03824 | ***thiN*** |  |  | thiamine diphosphokinase ThiN | H | | METABOLISM; coenzyme transport and metabolism | Core | Pro |
| 426 | MGCS36044_03826 | ***rpe*** |  |  | ribulose-phosphate 3-epimerase Rpe | G | | METABOLISM; carbohydrate transport and metabolism | Core | Pro |
| 427 | MGCS36044_03828 | ***rsgA*** |  |  | ribosome small subunit-dependent GTPase (A) RgsA | J | | INFORMATION STORAGE AND PROCESSING; translation, ribosomal structure and biogenesis | Core | Pro |
| 428 | MGCS36044_03834 | ***rrmV*** |  |  | 5S rRNA maturation endonuclease RnmV | J | | INFORMATION STORAGE AND PROCESSING; translation, ribosomal structure and biogenesis | Core | Pro |
| 429 | MGCS36044_03858 | ***rpmH*** |  |  | 50S ribosomal L34 protein RpmH | J | | INFORMATION STORAGE AND PROCESSING; translation, ribosomal structure and biogenesis | Core | Pro |
| 430 | MGCS36044_03864 | ***rnpA*** |  |  | ribonuclease P protein component RnpA | J | | INFORMATION STORAGE AND PROCESSING; translation, ribosomal structure and biogenesis | Core | Pro |
| 431 | MGCS36044_03874 | ***gltX*** |  |  | glutamate--tRNA ligase | J | | INFORMATION STORAGE AND PROCESSING; translation, ribosomal structure and biogenesis | Core | Pro |
| 432 | MGCS36044_03884 | ***-*** |  |  | carbonic anhydrase | P | | METABOLISM; inorganic ion transport and metabolism | Core | Pro |
| 433 | MGCS36044_03898 | ***gpsA*** |  |  | NAD(P)H-dependent glycerol-3-phosphate dehydrogenase GpsA | I | | METABOLISM; lipid metabolism | Core | Pro |
| 434 | MGCS36044_03900 | ***galU*** |  |  | UTP--glucose-1-phosphate uridylyltransferase GalU | M | | CELLULAR PROCESSES AND SIGNALING; cell envelope biogenesis, outermembrane | Core | Pro |
| 435 | MGCS36044_03920 | ***pgi*** |  |  | Pgi family glucose-6-phosphate isomerase | G | | METABOLISM; carbohydrate transport and metabolism | Core | Pro |
| 436 | MGCS36044_03928 | ***-*** |  |  | Bacteria_small_SRP | | |  | Core | RNA |
| 437 | MGCS36044_03974 | ***leuS*** |  |  | leucine--tRNA synthase LeuS | J | | INFORMATION STORAGE AND PROCESSING; translation, ribosomal structure and biogenesis | Core | Pro |
| 438 | MGCS36044_03978 | ***-*** |  |  | IS30 family transposase | | |  | Core | Pro |
| 439 | MGCS36044_03980 | ***-*** |  |  | disrupted IS3 family transposase encoding gene | | | | Core | Pro |
| 440 | MGCS36044_03990 | ***-*** |  |  | IS982 family transposase | | |  | Core | Pro |
| 441 | MGCS36044_03996 | ***ifs*** |  |  | nicotine adenine dinucleotide glycohydrolase inhibitor Ifs | R | | General function prediction only | Core | Pro |
| 442 | MGCS36044_04000 | ***nusG*** |  |  | transcription antitermination protein NusG | K | | INFORMATION STORAGE AND PROCESSING; transcription | Core | Pro |
| 443 | MGCS36044_04002 | ***secE*** |  |  | preprotein translocase subunit protein SecE | U | | CELLULAR PROCESSES AND SIGNALING; intracellular trafficking, secretion, and vesicular transport | Core | Pro |
| 444 | MGCS36044_04018 | ***groEL*** |  |  | chaperonin GroEL | O | | CELLULAR PROCESSES AND SIGNALING; post-translational modification, protein turnover, and chaperones | Core | Pro |
| 445 | MGCS36044_04020 | ***groES*** |  |  | co-chaperone GroES | O | | CELLULAR PROCESSES AND SIGNALING; post-translational modification, protein turnover, and chaperones | Core | Pro |
| 446 | MGCS36044_04032 | ***-*** |  |  | disrupted IS1182 family transposase encoding gene | | | | Core | Pro |
| 447 | MGCS36044_04060 | ***rpsB*** |  |  | 30S ribosomal S2 protein RpsB | J | | INFORMATION STORAGE AND PROCESSING; translation, ribosomal structure and biogenesis | Core | Pro |
| 448 | MGCS36044_04062 | ***tsf*** |  |  | translation elongation factor Tsf | J | | INFORMATION STORAGE AND PROCESSING; translation, ribosomal structure and biogenesis | Core | Pro |
| 449 | MGCS36044_04066 | ***treC*** |  |  | trehalose-6-phosphate hydrolase TreC | G | | METABOLISM; carbohydrate transport and metabolism | Core | Pro |
| 450 | MGCS36044_04096 | ***-*** |  |  | DUF1292 domain-containing protein | S | | POORLY CHARACTERIZED; function unknown | Core | Pro |
| 451 | MGCS36044_04098 | ***ruvX*** |  |  | Holliday junction resolvase RuvX | K | | INFORMATION STORAGE AND PROCESSING; transcription | Core | Pro |
| 452 | MGCS36044_04100 | ***-*** |  |  | IreB-related regulatory phosphoprotein | S | | POORLY CHARACTERIZED; function unknown | Core | Pro |
| 453 | MGCS36044_04104 | ***spxA_2*** |  |  | transcriptional regulator SpxA | P | | METABOLISM; inorganic ion transport and metabolism | Core | Pro |
| 454 | MGCS36044_04108 | ***recA*** |  |  | recombinase RecA | L | | INFORMATION STORAGE AND PROCESSING; replication, recombination and repair | Core | Pro |
| 455 | MGCS36044_04116 | ***ruvA*** |  |  | Holliday junction ATP-dependent DNA helicase RuvA | L | | INFORMATION STORAGE AND PROCESSING; replication, recombination and repair | Core | Pro |
| 456 | MGCS36044_04128 | ***argS*** |  |  | arginine--tRNA synthase ArgS | J | | INFORMATION STORAGE AND PROCESSING; translation, ribosomal structure and biogenesis | Core | Pro |
| 457 | MGCS36044_04138 | ***aspS*** |  |  | aspartyl-tRNA synthetase | J | | INFORMATION STORAGE AND PROCESSING; translation, ribosomal structure and biogenesis | Core | Pro |
| 458 | MGCS36044_04140 | ***hisS*** |  |  | histidine--tRNA synthase HisS | J | | INFORMATION STORAGE AND PROCESSING; translation, ribosomal structure and biogenesis | Core | Pro |
| 459 | MGCS36044_04142 | ***rpmF*** |  |  | 50S ribosomal L32 protein RpmF | J | | INFORMATION STORAGE AND PROCESSING; translation, ribosomal structure and biogenesis | Core | Pro |
| 460 | MGCS36044_04200 | ***-*** |  |  | IS1182 family transposase | | |  | Core | Pro |
| 461 | MGCS36044_04202 | ***rpsD*** |  |  | 30S ribosomal S4 protein RpsD | J | | INFORMATION STORAGE AND PROCESSING; translation, ribosomal structure and biogenesis | Core | Pro |
| 462 | MGCS36044_04206 | ***-*** |  |  | Veg family protein | R | | General function prediction only | Core | Pro |
| 463 | MGCS36044_04208 | ***dnaC*** |  |  | replicative DNA helicase DnaC | L | | INFORMATION STORAGE AND PROCESSING; replication, recombination and repair | Core | Pro |
| 464 | MGCS36044_04210 | ***rplI*** |  |  | 50S ribosomal L9 protein RplI | J | | INFORMATION STORAGE AND PROCESSING; translation, ribosomal structure and biogenesis | Core | Pro |
| 465 | MGCS36044_04212 | ***-*** |  |  | DHH family phosphoesterase | R | | General function prediction only | Core | Pro |
| 466 | MGCS36044_04214 | ***-*** |  |  | IS1548 family transposase | L | | INFORMATION STORAGE AND PROCESSING; replication, recombination and repair | Core | Pro |
| 467 | MGCS36044_04216 | ***mnmG*** |  |  | tRNA uridine-5-carboxymethylaminomethyl(34) synthesis enzyme MnmG | J | | INFORMATION STORAGE AND PROCESSING; translation, ribosomal structure and biogenesis | Core | Pro |
| 468 | MGCS36044_04220 | ***-*** |  |  | Spd-sr37 |  | |  | Core | RNA |
| 469 | MGCS36044_04224 | ***mnmA*** |  |  | tRNA 2-thiouridine(34) synthase MnmA | J | | INFORMATION STORAGE AND PROCESSING; translation, ribosomal structure and biogenesis | Core | Pro |
| 470 | MGCS36044_04234 | ***cbiQ_2*** |  |  | cobalt ABC transporter permease CbiQ | P | | METABOLISM; inorganic ion transport and metabolism | Core | Pro |
| 471 | MGCS36044_04236 | ***cbiO2*** |  |  | cobalt ABC transporter ATPase CbiO1 | P | | METABOLISM; inorganic ion transport and metabolism | Core | Pro |
| 472 | MGCS36044_04238 | ***cbiO1*** |  |  | cobalt ABC transporter ATPase CbiO2 | P | | METABOLISM; inorganic ion transport and metabolism | Core | Pro |
| 473 | MGCS36044_04240 | ***pgsA*** |  |  | CDP-diacylglycerol--glycerol-3-phosphate 3-phosphatidyltransferase PgsA | I | | METABOLISM; lipid metabolism | Core | Pro |
| 474 | MGCS36044_04256 | ***trpS*** |  |  | tryptophanyl-tRNA synthetase | J | | INFORMATION STORAGE AND PROCESSING; translation, ribosomal structure and biogenesis | Core | Pro |
| 475 | MGCS36044_04268 | ***-*** |  |  | disrupted IS30 family transposase encoding gene | | | | Core | Pro |

† Genes are ordered ascendingly based on the locus tag numbers.

‡ Putative virulence genes (Eraso et al., 2024, *mBio*).

¶ As defined by the Prokka annotator (signal P.6); Secreted: secreted outside the cell, Lipo: membrane lipoprotein.

††Genomic location. Core: the core genome, Acc: the accessory genome, ROD: regions of difference.

‡‡ Pro: protein.
