## Supplementary material for "Gene contribution of *Streptococcus dysgalactiae* subspecies *equisimilis*, an emerging pathogen, to experimental primate necrotizing myositis": Table S1

**Table S1. Primers used in this study**

| **Primer** | **DNA sequence** |
| --- | --- |
| A1^1^ | [PHOS]-GAT CGG AAG AGC ACA CGT CT |
| A2^2^ | ACA CTC TTT CCC TAC ACG ACG CTC TTC CGA TC*T |
| ISS*1* | AAT GAT ACG GCG ACC ACC GAG ATC TAC ACG TTC ATT GAT ATA TCC TCG CTG |
| Read 1 SEQ | GTT CAT TGA TAT ATC CTC GCT GTC ATT TTT ATT CAT TTT ACA CTA AAA TAG ACT TAT |
| Index SEQ | AGA TCG GAA GAG CGT CGT GTA GGG AAA GAG TGT |
| Index primer 1 | CAA GCA GAA GAC GGC ATA CGA GAT CGG TTC GCC TTA ACA CTC TTT C |
| Index primer 2 | CAA GCA GAA GAC GGC ATA CGA GAT CGG TCT AGT ACG ACA CTC TTT C |
| Index primer 3 | CAA GCA GAA GAC GGC ATA CGA GAT CGG TTT CTG CCT ACA CTC TTT C |
| Index primer 4 | CAA GCA GAA GAC GGC ATA CGA GAT CGG TGC TCA GGA ACA CTC TTT C |
| Index primer 5 | CAA GCA GAA GAC GGC ATA CGA GAT CGG TAG GAG TCC ACA CTC TTT C |
| Index primer 6 | CAA GCA GAA GAC GGC ATA CGA GAT CGG TCA TGC CTA ACA CTC TTT C |
| Index primer 7 | CAA GCA GAA GAC GGC ATA CGA GAT CGG TGT AGA GAG ACA CTC TTT C |
| Index primer 8 | CAA GCA GAA GAC GGC ATA CGA GAT CGG TCC TCT CTG ACA CTC TTT C |
| 1. [PHOS], phosphorylated 2. ^*,^ Phosphorothioate bond | |
