## Supplementary material for "Gene contribution of *Streptococcus dysgalactiae* subspecies *equisimilis*, an emerging pathogen, to experimental primate necrotizing myositis": Table S4E

**Table S4E. Conditionally essential genes in MGCS36044 and MGCS36089 *in vitro*, but not *in vivo*.**

| **No. ^†^** | **Locus tag 044^∫^** | **Locus tag 089^∫∫^** | **Gene** | **Function** | **COG class** | **COG description** | **Core/ Acc^††^** | **Pro/ RNA^‡‡^** |
| --- | --- | --- | --- | --- | --- | --- | --- | --- |
| 1 | MGCS36044_00034 | MGCS36089_00034 | ***-*** | IS30 family transposase |  |  | Core | Pro |
| 2 | MGCS36044_00150 | MGCS36089_00150 | ***-*** | hypothetical protein | S | POORLY CHARACTERIZED; function unknown | Core | Pro |
| 3 | MGCS36044_00166 | MGCS36089_00166 | ***rpsJ*** | 30S ribosomal S10 protein RpsJ | J | INFORMATION STORAGE AND PROCESSING; translation, ribosomal structure and biogenesis | Core | Pro |
| 4 | MGCS36044_00182 | MGCS36089_00182 | ***rplP*** | 50S ribosomal L29 protein RplP | J | INFORMATION STORAGE AND PROCESSING; translation, ribosomal structure and biogenesis | Core | Pro |
| 5 | MGCS36044_00468 | MGCS36089_00468 | ***-*** | hypothetical protein | S | POORLY CHARACTERIZED; function unknown | ROD.2 | Pro |
| 6 | MGCS36044_00512 | MGCS36089_00512 | ***nrdI_1*** | class Ib ribonucleoside-diphosphate reductase assembly flavoprotein NrdI | F | METABOLISM; nucleotide transport and metabolism | Core | Pro |
| 7 | MGCS36044_01074 | MGCS36089_01060 | ***ftsK*** | cell division protein FtsK | D | CELLULAR PROCESSES AND SIGNALING; cell division and chromosome partitioning | Core | Pro |
| 8 | MGCS36044_02134 | MGCS36089_02126 | ***ylxM*** | YlxM superfamily signal recognition particle associated DNA-binding protein | J | INFORMATION STORAGE AND PROCESSING; translation, ribosomal structure and biogenesis | Core | Pro |
| 9 | MGCS36044_02350 | MGCS36089_02342 | ***alsT*** | sodium:alanine symporter family protein | E | METABOLISM; amino acid transport and metabolism | Core | Pro |
| 10 | MGCS36044_02676 | MGCS36089_02688 | ***dltX*** | teichoic acid D-Ala incorporation-associated protein DltX | M | CELLULAR PROCESSES AND SIGNALING; cell wall/membrane/envelope biogenesis | Core | Pro |
| 11 | MGCS36044_03052 | MGCS36089_03064 | ***sepF*** | cell division protein SepF | D | CELLULAR PROCESSES AND SIGNALING; cell division and chromosome partitioning | Core | Pro |
| 12 | MGCS36044_03422 | MGCS36089_03434 | ***lacD_2*** | tagatose-bisphosphate aldolase LacD | G | METABOLISM; carbohydrate transport and metabolism | Core | Pro |
| 13 | MGCS36044_03624 | MGCS36089_03636 | ***-*** | IS30 family transposase |  |  | Core | Pro |
| 14 | MGCS36044_03798 | MGCS36089_03810 | ***glnA*** | glutamine synthetase GlnA | E | METABOLISM; amino acid transport and metabolism | Core | Pro |
| 15 | MGCS36044_03910 | MGCS36089_03922 | ***-*** | IS30 family transposase |  |  | Core | Pro |
| 16 | MGCS36044_03918 | MGCS36089_03930 | ***-*** | disrupted IS30 family transposase encoding gene | |  | Core | Pro |
| 17 | MGCS36044_04178 | MGCS36089_04192 | ***-*** | IS982 family transposase |  |  | ROD.9 | Pro |

† Genes are ordered ascendingly based on the locus tag numbers.

**^∫^** Refers to MGCS36044

**^∫∫^** Refers to MGCS36089

††Genomic location. Core: the core genome, Acc: the accessory genome, ROD: regions of difference.

‡‡ Pro: protein.
