## Supplementary material for "Gene contribution of *Streptococcus dysgalactiae* subspecies *equisimilis*, an emerging pathogen, to experimental primate necrotizing myositis": Table S3B

**Table S3B. Essential MGCS36089 genes during growth *in vitro* but not in *vivo***

| **No. ^†^** | **Locus tag** | **Gene** | **Virulence^‡^** | **Function** | **COG class** | **COG description** | **Core/ Acc^††^** | **Pro/ RNA^‡‡^** |
| --- | --- | --- | --- | --- | --- | --- | --- | --- |
| 1 | MGCS36089_00034 | ***-*** |  | IS30 family transposase | |  | Core | Pro |
| 2 | MGCS36089_00146 | ***ruvB*** |  | Holliday junction branch migration DNA helicase | L | INFORMATION STORAGE AND PROCESSING; Dna replication, recombination and repair | Core | Pro |
| 3 | MGCS36089_00150 | ***-*** |  | hypothetical protein | S | POORLY CHARACTERIZED; function unknown | Core | Pro |
| 4 | MGCS36089_00164 | ***-*** |  | IS30 family transposase | |  | Core | Pro |
| 5 | MGCS36089_00166 | ***rpsJ*** |  | 30S ribosomal S10 protein RpsJ | J | INFORMATION STORAGE AND PROCESSING; translation, ribosomal structure and biogenesis | Core | Pro |
| 6 | MGCS36089_00182 | ***rplP*** |  | 50S ribosomal L29 protein RplP | J | INFORMATION STORAGE AND PROCESSING; translation, ribosomal structure and biogenesis | Core | Pro |
| 7 | MGCS36089_00202 | ***rpsE*** |  | 30S ribosomal S5 protein RpsE | J | INFORMATION STORAGE AND PROCESSING; translation, ribosomal structure and biogenesis | Core | Pro |
| 8 | MGCS36089_00206 | ***rplO*** |  | 50S ribosomal L15 protein RplO | J | INFORMATION STORAGE AND PROCESSING; translation, ribosomal structure and biogenesis | Core | Pro |
| 9 | MGCS36089_00210 | ***adk*** |  | adenylate kinase protein Adk | F | METABOLISM; nucleotide transport and metabolism | Core | Pro |
| 10 | MGCS36089_00390 | ***purA*** |  | adenylosuccinate synthase PurA | F | METABOLISM; nucleotide transport and metabolism | Core | Pro |
| 11 | MGCS36089_00468 | ***-*** |  | hypothetical protein | S | POORLY CHARACTERIZED; function unknown | ROD.2 | Pro |
| 12 | MGCS36089_00512 | ***nrdI_1*** |  | class Ib ribonucleoside-diphosphate reductase | F | METABOLISM; nucleotide transport and metabolism | Core | Pro |
| 13 | MGCS36089_00542 | ***cdsA*** |  | phosphatidate cytidylyltransferase CdsA | I | METABOLISM; lipid metabolism | Core | Pro |
| 14 | MGCS36089_00758 | ***-*** |  | IS1548 family transposase | L | INFORMATION STORAGE AND PROCESSING; Dna replication, recombination and repair | Core | Pro |
| 15 | MGCS36089_00810 | ***yqeG*** |  | HAD IIIA-type phosphatase YqeG | F | METABOLISM; nucleotide transport and metabolism | Core | Pro |
| 16 | MGCS36089_00816 | ***nadD*** |  | nicotinate-nucleotide adenylyltransferase NadD | H | METABOLISM; coenzyme transport and metabolism | Core | Pro |
| 17 | MGCS36089_00862 | ***yceD*** |  | large ribosomal RNA subunit accumulation protein | M | CELLULAR PROCESSES AND SIGNALING; cell envelope biogenesis, outermembrane | Core | Pro |
| 18 | MGCS36089_00864 | ***covR*** | Virulence | TCS DNA-binding response regulator CovR | T | CELLULAR PROCESSES AND SIGNALING; signal transduction mechanisms | Core | Pro |
| 19 | MGCS36089_00886 | ***greA*** |  | transcription elongation factor GreA | K | INFORMATION STORAGE AND PROCESSING; transcription | Core | Pro |
| 20 | MGCS36089_00924 | ***-*** |  | ECF transporter S component | I | METABOLISM; lipid metabolism | Core | Pro |
| 21 | MGCS36089_00954 | ***clpP*** |  | ATP-dependent Clp protease proteolytic subunit | O | CELLULAR PROCESSES AND SIGNALING; post-translational modification, protein turnover, and chaperones | Core | Pro |
| 22 | MGCS36089_01046 | ***mtnN*** |  | 5'-methylthioadenosine/adenosylhomocysteine | F | METABOLISM; nucleotide transport and metabolism | Core | Pro |
| 23 | MGCS36089_01056 | ***mtsC*** |  | metal ABC transporter permease MtsC | P | METABOLISM; inorganic ion transport and metabolism | Core | Pro |
| 24 | MGCS36089_01060 | ***ftsK*** |  | cell division protein FtsK | D | CELLULAR PROCESSES AND SIGNALING; cell division and chromosome partitioning | Core | Pro |
| 25 | MGCS36089_01062 | ***-*** |  | DUF3397 domain-containing protein | S | POORLY CHARACTERIZED; function unknown | Core | Pro |
| 26 | MGCS36089_01066 | ***rplA*** |  | 50S ribosomal L1 protein RplA | J | INFORMATION STORAGE AND PROCESSING; translation, ribosomal structure and biogenesis | Core | Pro |
| 27 | MGCS36089_01116 | ***-*** |  | IS1548 family transposase | L | INFORMATION STORAGE AND PROCESSING; Dna replication, recombination and repair | Core | Pro |
| 28 | MGCS36089_01226 | ***vicK*** | Virulence | TCS signal transduction sensor kinase VicK | T | CELLULAR PROCESSES AND SIGNALING; signal transduction mechanisms | Core | Pro |
| 29 | MGCS36089_01230 | ***rnc*** |  | ribonuclease III Rnc | A | INFORMATION STORAGE AND PROCESSING; RNA processing and modification | Core | Pro |
| 30 | MGCS36089_01238 | ***-*** |  | IS1548 family transposase | L | INFORMATION STORAGE AND PROCESSING; Dna replication, recombination and repair | Core | Pro |
| 31 | MGCS36089_01318 | ***thiT*** |  | energy-coupled thiamine transporter ThiT | H | METABOLISM; coenzyme transport and metabolism | Core | Pro |
| 32 | MGCS36089_01324 | ***-*** |  | IS110 family transposase | |  | Core | Pro |
| 33 | MGCS36089_01336 | ***ftsW*** |  | cell division protein FtsW | D | CELLULAR PROCESSES AND SIGNALING; cell division and chromosome partitioning | Core | Pro |
| 34 | MGCS36089_01426 | ***asnC*** |  | asparaginyl-tRNA synthetase protein AsnC | J | INFORMATION STORAGE AND PROCESSING; translation, ribosomal structure and biogenesis | Core | Pro |
| 35 | MGCS36089_01458 | ***rpmE*** |  | 50S ribosomal L31 type B protein RpmE | J | INFORMATION STORAGE AND PROCESSING; translation, ribosomal structure and biogenesis | Core | Pro |
| 36 | MGCS36089_01552 | ***atpC*** |  | ATP synthase epsilon subunit AtpC | C | METABOLISM; energy production and conversion | Core | Pro |
| 37 | MGCS36089_01576 | ***rexB*** |  | ATP-dependent nuclease B subunit RexB | L | INFORMATION STORAGE AND PROCESSING; Dna replication, recombination and repair | Core | Pro |
| 38 | MGCS36089_01584 | ***-*** |  | IS1548 family transposase | L | INFORMATION STORAGE AND PROCESSING; Dna replication, recombination and repair | Core | Pro |
| 39 | MGCS36089_01622 | ***-*** |  | RfbX superfamily lipopolysaccharide biosynthesis | M | CELLULAR PROCESSES AND SIGNALING; cell envelope biogenesis, outermembrane | Core | Pro |
| 40 | MGCS36089_01644 | ***aroD*** |  | type I 3-dehydroquinate dehydratase AroD | E | METABOLISM; amino acid transport and metabolism | Core | Pro |
| 41 | MGCS36089_01792 | ***rplJ*** |  | 50S ribosomal L10 protein RplJ | J | INFORMATION STORAGE AND PROCESSING; translation, ribosomal structure and biogenesis | Core | Pro |
| 42 | MGCS36089_02072 | ***-*** |  | IS30 family transposase | |  | Core | Pro |
| 43 | MGCS36089_02126 | ***ylxM*** |  | YlxM superfamily signal recognition particle | J | INFORMATION STORAGE AND PROCESSING; translation, ribosomal structure and biogenesis | Core | Pro |
| 44 | MGCS36089_02340 | ***-*** |  | Glycine. Infernal predicted noncoding RNA, Rfam accession | | | Core | RNA |
| 45 | MGCS36089_02342 | ***alsT*** |  | sodium:alanine symporter family protein | E | METABOLISM; amino acid transport and metabolism | Core | Pro |
| 46 | MGCS36089_02378 | ***-*** |  | IS1182 family transposase | |  | Core | Pro |
| 47 | MGCS36089_02434 | ***rpiA*** |  | ribose-5-phosphate isomerase RpiA | G | METABOLISM; carbohydrate transport and metabolism; Pentose-P pathway | Core | Pro |
| 48 | MGCS36089_02506 | ***-*** |  | CRISPR-DR22. Infernal predicted noncoding RNA, Rfam accession | | | Core | Pro |
| 49 | MGCS36089_02620 | ***trmK*** |  | tRNA (adenine(22)-N(1))-methyltransferase TrmK | L | INFORMATION STORAGE AND PROCESSING; Dna replication, recombination and repair | Core | Pro |
| 50 | MGCS36089_02624 | ***dnaD*** |  | DNA replication protein DnaD | L | INFORMATION STORAGE AND PROCESSING; Dna replication, recombination and repair | Core | Pro |
| 51 | MGCS36089_02688 | ***dltX*** |  | teichoic acid D-Ala incorporation-associated | M | CELLULAR PROCESSES AND SIGNALING; cell envelope biogenesis, outermembrane | Core | Pro |
| 52 | MGCS36089_02742 | ***-*** |  | disrupted IS30 transposase encoding gene | | | Core | Pro |
| 53 | MGCS36089_02844 | ***brkB*** |  | BrkB family protein | |  | Core | Pro |
| 54 | MGCS36089_02898 | ***ptsI*** |  | phosphoenolpyruvate--protein phosphotransferase | G | METABOLISM; carbohydrate transport and metabolism; Pentose-P pathway | Core | Pro |
| 55 | MGCS36089_02950 | ***holA*** |  | DNA polymerase III delta subunit HolA | L | INFORMATION STORAGE AND PROCESSING; Dna replication, recombination and repair | Core | Pro |
| 56 | MGCS36089_03030 | ***folD*** |  | bifunctional methylenetetrahydrofolate dehydrogenase/methenyltetrahydrofolate cyclohydrolase FolD protein | E | METABOLISM; amino acid transport and metabolism | Core | Pro |
| 57 | MGCS36089_03064 | ***sepF*** |  | cell division protein SepF | D | CELLULAR PROCESSES AND SIGNALING; cell division and chromosome partitioning | Core | Pro |
| 58 | MGCS36089_03076 | ***murD*** |  | UDP-N-acetylmuramoyl-L-alanine--D-glutamate | M | CELLULAR PROCESSES AND SIGNALING; cell envelope biogenesis, outermembrane | Core | Pro |
| 59 | MGCS36089_03124 | ***coaD*** |  | pantetheine-phosphate adenylyltransferase CoaD | H | METABOLISM; coenzyme transport and metabolism | Core | Pro |
| 60 | MGCS36089_03300 | ***cysK*** |  | cysteine synthase A CysK | E | METABOLISM; amino acid transport and metabolism | Core | Pro |
| 61 | MGCS36089_03306 | ***liaR*** | Virulence | three component system signal transduction | K | INFORMATION STORAGE AND PROCESSING; transcription | Core | Pro |
| 62 | MGCS36089_03314 | ***pppL*** |  | Stp1/IreP family PP2C-type Ser/Thr phosphatase | T | CELLULAR PROCESSES AND SIGNALING; signal transduction mechanisms | Core | Pro |
| 63 | MGCS36089_03434 | ***lacD_2*** |  | tagatose-bisphosphate aldolase LacD | G | METABOLISM; carbohydrate transport and metabolism; Pentose-P pathway | Core | Pro |
| 64 | MGCS36089_03484 | ***rimP*** |  | ribosome maturation factor RimP | J | INFORMATION STORAGE AND PROCESSING; translation, ribosomal structure and biogenesis | Core | Pro |
| 65 | MGCS36089_03534 | ***accB*** |  | acetyl-CoA carboxylase biotin carboxyl carrier | I | METABOLISM; lipid metabolism | Core | Pro |
| 66 | MGCS36089_03554 | ***-*** |  | PfpI family predicted protease/amidase | | | Core | Pro |
| 67 | MGCS36089_03630 | ***alr*** |  | alanine racemase Alr | E | METABOLISM; amino acid transport and metabolism | Core | Pro |
| 68 | MGCS36089_03636 | ***-*** |  | IS30 family transposase | |  | Core | Pro |
| 69 | MGCS36089_03648 | ***nusB*** |  | transcription termination protein NusB | K | INFORMATION STORAGE AND PROCESSING; transcription | Core | Pro |
| 70 | MGCS36089_03650 | ***-*** |  | Asp23/Gls24 family envelope stress response | R | POORLY CHARACTERIZED; General function prediction only | Core | Pro |
| 71 | MGCS36089_03750 | ***-*** |  | disrupted IS3 family transposase encoding gene | | | ROD.8 | Pro |
| 72 | MGCS36089_03810 | ***glnA*** |  | glutamine synthetase GlnA | E | METABOLISM; amino acid transport and metabolism | Core | Pro |
| 73 | MGCS36089_03836 | ***thiN*** |  | thiamine diphosphokinase ThiN | H | METABOLISM; coenzyme transport and metabolism | Core | Pro |
| 74 | MGCS36089_03838 | ***rpe*** |  | ribulose-phosphate 3-epimerase Rpe | G | METABOLISM; carbohydrate transport and metabolism; Pentose-P pathway | Core | Pro |
| 75 | MGCS36089_03922 | ***-*** |  | IS30 family transposase | |  | Core | Pro |
| 76 | MGCS36089_03930 | ***-*** |  | disrupted IS30 family transposase encoding gene | | | Core | Pro |
| 77 | MGCS36089_03994 | ***-*** |  | disrupted IS3 family transposase encoding gene | | | Core | Pro |
| 78 | MGCS36089_04004 | ***-*** |  | IS982 family transposase | |  | Core | Pro |
| 79 | MGCS36089_04014 | ***nusG*** |  | transcription antitermination protein NusG | K | INFORMATION STORAGE AND PROCESSING; transcription | Core | Pro |
| 80 | MGCS36089_04122 | ***recA*** |  | recombinase RecA | L | INFORMATION STORAGE AND PROCESSING; Dna replication, recombination and repair | Core | Pro |
| 81 | MGCS36089_04192 | ***-*** |  | IS982 family transposase | |  | ROD.9 | Pro |
| 82 | MGCS36089_04216 | ***rpsD*** |  | 30S ribosomal S4 protein RpsD | J | INFORMATION STORAGE AND PROCESSING; translation, ribosomal structure and biogenesis | Core | Pro |

† Genes are ordered ascendingly based on the locus tag numbers.

‡ Putative virulence genes (Eraso et al., 2024, *mBio*).

††Genomic location. Core: the core genome, Acc: the accessory genome, ROD: regions of difference.

‡‡ Pro: protein.
